# Chromosome-Level Genome Assembly and Circadian Gene Repertoire of the Patagonia blennie *Eleginops maclovinus* - the closest ancestral proxy of Antarctic cryonotothenioids

**DOI:** 10.1101/2023.04.24.537882

**Authors:** C.-H. Christina Cheng, Angel G. Rivera-Colón, Bushra Fazal Minhas, Loralee Wilson, Niraj Rayamajhi, Luis Vargas-Chacoff, Julian M. Catchen

## Abstract

The basal S. American notothenioid *Eleginops maclovinus* (Patagonia blennie) occupies a uniquely important phylogenetic position in Notothenioidei as the singular closest sister species to the Antarctic cryonotothenioid fishes. Its genome and the traits encoded therein would be nearest representatives of the temperate ancestor from which the Antarctic clade arose, providing an ancestral reference for deducing polar derived changes. In this study, we generated a gene- and chromosome-complete assembly of *E. maclovinus* genome using long read sequencing and HiC scaffolding. We compared its genome architecture with the more basally divergent *Cottoperca gobio* and the derived genomes of nine cryonotothenioids representing all five Antarctic families. We additionally curated its repertoire of circadian rhythm genes, ascertained their functionality by transcriptome sequencing, and compared its pattern of gene retention with *C. gobio* and the derived cryonotothenioids. Both analyses found *E. maclovinus* to share greater conservation with the Antarctic clade, solidifying its evolutionary status as direct sister and best suited ancestral proxy of cryonotothenioids. The high quality genome of *E. maclovinus* will facilitate inquiries into cold derived traits in temperate to polar evolution, as well as inform on the paths of readaptation to non-freezing habitats in various secondarily temperate cryonotothenioids through comparative genomic analyses.

## 1. Introduction

The perciform suborder Notothenioidei consists of non-Antarctic and Antarctic lineages, historically classified in eight taxonomic families [1]. Three basal families -Bovichtidae, Pseudaphritidae, and Eleginopidae diverged along the southern coasts of Southern Hemisphere land masses and never experienced polar climate adaptation. The other five families – Nototheniidae, Artedidraconidae, Harpagiferidae, Bathydraconidae, and Channichthyidae evolved and diversified *in situ* in the oceanographically and thermally isolated Southern Ocean (SO), forming an adaptive radiation and became the dominant fish group endemic to the freezing SO [2,3]. As a rare marine species flock [4,5] that successfully overcame and flourished in otherwise uninhabitable icy freezing conditions for ectothermic teleosts, the Antarctic clade of notothenioid fishes (cryonotothenioids) have been the subject of decades of numerous studies covering all aspects of its biology and evolution.

The extraordinary ecological success of cryonotothenioids resulted from the evolutionary innovation of macromolecular antifreeze glycoproteins (AFGP) [6,7], which avert death from freezing by preventing internal ice nucleation [8,9], along with biochemical and physiological adaptive changes that enable cellular and life processes at rate-depressive and protein denaturing subzero temperatures [10-12]. Becoming highly adapted and specialized to chronic cold apparently came at the cost of developing extreme stenothermy, with extant cryonotothenioids exhibiting greatly reduced thermal tolerance [13,14]. In parallel, certain essential traits have been lost, hypothesized to result from relaxation of selective pressure in the polar environment for their maintenance. The most ostensible are the loss of the classic inducible heat shock response (HSR) across cryonotothenioids [15-17] presumably due to the lack of large thermal variations in the cold-stable SO, and the loss of hemoglobin and red blood cells in the icefish family Channichthyidae [18,19], where stably cold and oxygen-rich SO waters presumably provide a reliable oxygen source that made simple diffusion a viable mechanism for oxygen transport.

Understanding the cold adaptive and specialized states in cryonotothenioids continue to be active pursuits, particularly in light of the vulnerability of these stenothermal fishes to increasing threats of climate change. Thus added to the historical interest of how cryonotothenioids evolved cold adaptation and specialization to dominate the SO fish fauna, are pressing questions regarding whether their derived polar character harbors sufficient plasticity and adaptive potential to survive the changing contemporary SO environment. To this end, the fast trajectory of generating high quality genome assemblies of species across Notothenioidei promises to be an accelerator in our ability to address a wide range of important questions, including variations in genetic diversity, mutation rates, transposable element mobilization [12,20,21], genome structures [22], protein genes experiencing positive or relaxed selective pressures [23], and historical and extant demography (by whole genome resequencing of populations) shaped by Antarctic glacial cycles [24], among others, that would inform on the degree of resilience of the cryonotothenioids under climate change scenarios. Seventeen draft genomes of notothenioids with various degree of completeness have been published since 2014. A large step advance in genome resources in form of an additional two dozen new draft genomes are becoming publicly available [21].

Investigations of derived polar characters in cryonotothenioids by necessity would require having an appropriately related non-Antarctic species representing the ancestral state for comparison, to discern and/or test hypotheses of Antarctic-specific trait changes. Species of the three basally divergent non-Antarctic notothenioid families -Bovichtidae, Pseudaphritidae, and Eleginopidae all constitute candidates as ancestral proxy. A chromosome level genome of one bovichtid species *Cottoperca gobio* has been published [25], but it represents the most basally divergent lineage, which may be less informative in deciphering the temperate to polar transition. The Patagonia blennie (locally known as róbalo), *E. maclovinus*, the monotypic species of the family Eleginopidae, long recognized as the taxonomic and phylogenetic direct sister species of the Antarctic clade would serve as the most fitting closest ancestral state for evaluating trait changes and reconfigurations shaped by Southern Ocean selective pressures during cryonotothenioid evolution. A few studies have utilized *E. maclovinus* as the ancestral reference, namely the nature of the native chaperome [26], loss of the classical heat shock response [15], proliferation of transposable elements and a number of developmental traits during cold adaptation [20], and the state of the AFGP orthologous genomic region prior to the emergence of the AFGP genotype [22]. Apart from these more visible traits, there are expectedly myriad subtler trait changes across the genetic blueprint, requiring a well assembled and gene-complete genome to decode.

A draft genome of the Patagonia blennie has previously been reported using Illumina short reads, and is thus lacking contiguity [20]. In this study, we generated a highly contiguous and gene-complete, chromosome-level genome assembly of *E. maclovinus* using PacBio CLR (continuous long read) sequencing and HiC scaffolding. We found substantial difference in the genome structures between *C. gobio* and *E. maclovinus*, while *E. maclovinus* genome and those of nine cryonotothenioids species representing all five families of the Antarctic clade share larger degree of structural conservation. We further evaluated *E. maclovinus* as ancestral reference for addressing functional trait change in cryonotothenioids by characterizing the network of genes that choreograph the circadian rhythm, a trait that is likely distinct due to disparate light regimes between temperate and high polar latitudes. We found *E. maclovinus* to share a more similar pattern of retention of functional circadian genes with cryonotothenioids than *C. gobio*. These genome scale findings solidify *E. maclovinus* as an evolutionarily strategic sister species of the Antarctic clade, and its chromosome-level genome an important resource, in furthering our inquiries into cryonotothenioid evolution, past and present.

## 2. Materials and Methods

### 2.1 Specimen collection and sample preservation

Specimens of *E. maclovinus* were collected from two Chilean coastal water locations. In June 2008, several individuals were collected using rod and reel from the Patagonian waters near Puerto Natales, Chile (51.82°S, 72.63°S). We are deeply thankful to Mathias Hune for the collection. Blood cell DNA of one male individual from this collection was used for whole genome sequencing in this study. Fresh blood cells were collected from the live specimen in the field, washed with notothenioid fish ringer (0.1M sodium phosphate buffer, pH8.0, adjusted to 420 mOsm with NaCl) to remove plasma proteins, then embedded in 1% low-melt agarose blocks (BioRad Plug Kit #1703591) to preserve DNA integrity. The blood cells were lysed exhaustively in situ using 1% lithium dodecyl sulfate in notothenioid ringer, then stored in 0.5M EDTA pH 8.0 at 4oC until use. The handling and sampling of fish complied with Protocol #07053 approved by the University of Illinois Institutional Animal Care and Use Committee (IACUC).

A cohort of about 30 small juvenile *E. maclovinus* specimens were collected for a prior study [15] using a small seine in the near-shore water of Valdivia, Chile during the austral winter of 2014. Specimens were used for the study at the Laboratorio Costero de Recursos Acuáticos Calfuco of the Universidad Austral de Chile. Collection and aquarium maintenance of specimens were conducted with approved protocol 11/10 by the Comité de Bioética Animal, Universidad Austral de Chile. The post-experiment specimens were dissected, preserved in >10X volume of −20°C pre-chilled 90% ethanol, and stored at −20°C. Handling and sampling of fish were carried out in compliance with Protocol #12123 approved by the University of Illinois IACUC. In this study, pectoral fin tissue from twelve of the ethanol preserved individuals were taken for RNA isolation to obtain full-length transcripts using PacBio (Pacific Biosciences) Iso-Seq sequencing.

### 2.2. HMW DNA preparation and CLR sequencing

To isolate agarose-embedded high molecular weight (HMW) DNA, each agarose block was melted (70oC, 10 min.), and digested with 2 units of β-agarase (NEB) (42°C, one hr.). The released DNA was extracted using the Nanobind CBB Big DNA Kit and protocol (Circulomics) with adjustments in reagent volumes. DNA concentration was determined with Qubit Broad Range DNA (Invitrogen) fluorometry, purity with Nanodrop One (Fisher Scientific), and MW and integrity with pulsed field electrophoresis (CHEF Mapper, BioRad). The recovered *E. maclovinus* DNA showed high quality preservation (>50 to >150 kbp) through 12 years of storage prior to sequencing.

PacBio CLR (Continuous Long Read) library was prepared using unsheared HMW DNA, size selected for ≥45 kbp inserts using the BluePippin (Sage Science), and sequenced on two SMRT cells on Sequel II for 30 hours of data collection at the University of Oregon Genomics & Cell Characterization Core Facility (GC3F). A total of 10.12 million reads were obtained, with an average length of 10.86 Kbp, and a read N50 of 22.07 Kbp. Total read length was 110.7 Gb, equivalent to 142× genome coverage, based on a genome size of ∼780Mb previously estimated with flow cytometry of erythrocytes (mean c-value 0.798 pg) [20]. Based on assembled length of 606 Mbp (see Results and Discussion), the genome coverage was 180×.

### 2.3. Hi-C library and sequencing

For genome scaffolding, a chromatin proximity ligation Hi-C library was prepared from ethanol preserved liver of the same male *E. maclovinus* specimen by Phase Genomics Inc. using its Proximo Hi-C kit. The library was sequenced for paired-end 150bp on Illumina NovaSeq6000 at the Roy J. Carver Biotechnology Center, University of Illinois, Urbana-Champaign, which generated 204.5 million paired-end reads.

### 2.4. Genome assembly

Following a published assembly optimization strategy [27], we generated subsamples of the raw PacBio CLR read data covering various distributions of read lengths and depths of coverage, and assembled each subsampled dataset to arrive at the most optimal contig-level assembly. These contig-level assemblies were generated using *WTDBG2* v2.5 [28]. The assemblies were then self-corrected using the *GENOMICCONSENSUS* arrow v2.3.3 tool from PacBio (https://github.com/PacificBiosciences/GenomicConsensus), which identifies and fixes small indels and other errors in the assembled genome based on the sequence consensus present in the aligned raw reads. After polishing, each separate assembly was assessed based on contiguity metrics using *QUAST* v4.4 [29] and gene completeness metrics using *BUSCO* v3.0.1 [30] with the actinopterygii_odb9 reference ortholog set. The best contig-level assembly, *i.e*., the most contiguous and gene-complete, was obtained from reads 10-40 Kb in length, and a depth of coverage of 64×.

The optimal contig-level assembly was then integrated with the Hi-C data to generate chromosome-level scaffolds. The 3d-dna program from *JUICER* v1.6.2 [31] was used to identify Hi-C junctions and perform integration and ordering of the contigs into super-scaffolds. This chromosome-level assembly was reassessed for contiguity using *QUAST* and for gene-completeness using *BUSCO* v5.1.3 [32] with the actinopterygii_odb10 reference ortholog set. Additional manual inspection and curation of this scaffolding process was done using a conserved synteny analysis (methods described below), *e.g*., verifying that structural variants were not limited to the boundaries of a contig/scaffold, or that the large-scale organization of the chromosomes was supported by patterns of within-species synteny. Any manual changes made to the assembly were propagated to the corresponding structure (AGP), annotation (GTF), and sequence files (FASTA) using a custom Python program, as described in [22].

### 2.5 Repeat and protein-coding gene annotation

Repeat elements in the genome were annotated by first building a *de novo* repeat library with the *REPEATMODELER* v2.0.2a pipeline [33], using the BuildDatabase option and the NCBI database as input (-engine ncbi). We then used the RepeatModeler option to generate the final repeat library, using *LTRHARVEST* [34] to perform the discovery of LTRs (-LTRStruct). We then extracted teleost-specific repeats using the famdb.py script from *REPEATMASKER* (-species Teleostei), and combined these teleost-specific repeats with the *de novo* repeat library generated by *REPEATMODELER*. Final annotation and masking were done using *REPEATMASKER* v4.1.2-p1, using our custom library (-lib).

To annotate protein coding genes, we used the RNAseq data available in NCBI SRA under accession number SRX2523921 we previously generated to build a *de novo* reference transcriptome for *E. maclovinus* [15]. The RNAseq reads were aligned to the reference assembly using *STAR* [36] v2.7.1.a (--runMode alignReads). These alignments were used as input to the *BRAKER* v2.1.6 annotation pipeline [37,38] in transcript mode (--bam). In addition, we also ran *BRAKER* in protein mode, using the *ORTHODB* v10.1[39] zebrafish reference protein sequences as input (--prot_seq). The output of both *BRAKER* runs were then aggregated using *TSEBRA* v1.0.1 [40], obtaining a curated set of protein-coding gene annotations supported by both transcript and protein evidence.

### 2.6 Conserved synteny analysis

We used the *SYNOLOG* software [41,42] to evaluate conserved synteny and determine orthologous chromosomes between *E. maclovinus* and other notothenioid species. A total of ten notothenioid genomes were used for comparisons against *E. maclovinus*, which include the basal, non-Antarctic *Cottoperca gobio* (Bovichtidae) [25], five red-blooded cryonotothenioids across four different families (*Trematomus bernacchii* (Nototheniidae), *Dissostichus mawsoni* (Nototheniidae) [43], *Gymnodraco acuticeps* (Bathydraconidae) [21], *Harpagifer antarcticus* (Harpagiferidae) [21], and *Pogonophryne albipinna* (Artedidraconidae) (NCBI GCA_028583405.1), and four white-blooded icefishes (Channichthyidae; *Champsocephalus esox* and *Champsocephalus gunnari* [22], *Chaenocephalus aceratus* (NCBI GCA_023974075.1), and *Pseudochaenichthys georgianus* [21]). To run *SYNOLOG*, we first reciprocally matched the annotated gene models for all species using the blastp algorithm in *BLAST+* v2.4.0 [44]. The resulting *BLAST* outputs and the annotation coordinates then serve as input to *SYNOLOG*, which finds orthologous genes using reciprocal *BLAST* best hits and identifies blocks of conserved synteny based on the coordinates of the matched orthologs along a sliding window. The matching of orthologs is further refined by sequential rounds of synteny block identification, which allows for a more accurate identification of orthologous genes between species in the presence of paralogs.

In addition to these ten notothenioid genomes, we also included comparisons against three teleost outgroup species, including zebrafish (*Danio rerio*), threespine stickleback (*Gasterosteus aculeatus*) and platyfish (*Xiphophorous maculatus*), in order to validate the annotation of genes of interest (genes participating in circadian rhythm). In addition, this conserved synteny approach was used to name the *E. maclovinus* chromosomal scaffolds. The *E. maclovinus* chromosomes were numbered according to the orthologous platyfish chromosomes (Fig. S1), as this outgroup species serves as an example of the ancestral teleost karyotype of 24 chromosomes [45]. Accession information for the genome assemblies used for the comparative synteny analysis and circadian gene curation (next section) is given in Table S1.

### 2.7. Curation of predicted circadian network genes from genome assemblies

We performed a comparative analysis on the presence and absence of 33 genes of known circadian function (Table S2) in *E. maclovinus* and the 10 abovementioned notothenioid genome assemblies. First, we identified the respective IDs and sequences of these genes in the zebrafish (*D. rerio*); two of the 33 are absent in the zebrafish annotation, thus references from the threespine stickleback (*G. aculeatus*) were used. For each query assembly, we located the orthologs of the 33 model teleost reference genes by comparative synteny analysis in *SYNOLOG. SYNOLOG* identifies orthologous sequences first by identifying reciprocal *BLAST* best hits between a query and subject species (e.g., *E. maclovinus* vs. *D. rerio*), which are then refined by sequential rounds of conserved synteny identification. A reference gene was denoted as absent from a query assembly when no orthologous sequence could by identified by *SYNOLOG*. Once orthologs were identified in each query assembly, we used a Python script to extract the corresponding peptide sequences and annotation coordinates for use in downstream analyses.

### 2.8 Fin RNA isolation, Iso-Seq transcriptome sequencing and curation of circadian network gene transcripts

Teleost fins are known sites of peripheral circadian clock [46,47]. We evaluated functionality of the predicted circadian network genes in *E. maclovinus* by PacBio Iso-Seq sequencing of fin transcriptome. About 5-10 mg of pectoral fin tissue from each of the 12 ethanol preserved juveniles mentioned above were homogenized in Trizol (Invitrogen) with 0.5mm zirconium oxide beads using a Bullet Blender^R^ (Next Advance Inc.), followed by extraction of total RNA according to Invitrogen protocol. RNA purity and concentrations were initially assessed with the Epoch Take 3 microplate spectrophotometer (BioTek Instruments), and RNA integrity by visualizing ∼1 μg of RNA electrophoresed on a 1% formaldehyde agarose gel. Equal amounts (2 μg) of RNA from each sample were pooled into a single sample and treated with DNaseI with added RNase Inhibitor (New England Biolabs), then purified using MonarchR RNA Cleanup Kit (New England Biolabs). The quality of the cleaned, pooled RNA sample was checked using Qubit Broad Range RNA fluorometry (Invitrogen) for concentration, Nanodrop One (Fisher Scientific) for purity, and Agilent 5200 Fragment Analyzer (AATI) for RNA Quality Number; the pooled RNA sample achieved a RQN over 8.

Iso-Seq library construction and sequencing were carried out at the Roy J. Carver Biotechnology Center, University of Illinois, Urbana-Champaign. About 300ng of the pooled RNA sample was converted to cDNA with PacBio Iso-SeqR Express Oligo Kit, and the sequencing library was constructed with PacBio SMRTBell Express Template Prep kit 3.0. The library was sequenced on one SMRT cell 8M on a PacBio Sequel IIe using the CCS (circular consensus sequencing) mode for 30 hours of data collection. Over 3.3 million HiFi reads totaling 6.18 million bases were obtained. Removal of primers, trimming of polyA tails and other clean ups of the HiFi reads, followed by de novo clustering and consensus calling of transcripts were performed using PacBio *SMRTLINK* v.11 software, producing a final set of 167,428 high-quality (99.99% accuracy) consensus transcript isoforms.

The coding sequences of the 33 predicted circadian network genes were used as queries to identify their expressed transcripts from the high-quality Iso-Seq transcript dataset by blastn search. Since almost all circadian network genes occur in two or more paralogs, and blastn search would match a given query to all its paralogs, the assignment of the transcript/s specific to each paralog was based on >99%-100% nucleotide identity to the query gene model. The transcripts were extracted and aligned with the query gene model to ascertain absence of mutations.

## 3. Results and Discussion

*A priori*, the recognized position of *E. maclovinus* in notothenioid taxonomy and molecular phylogeny as the closest non-Antarctic sister species to the Antarctic nototothenioid radiation indicates its suitability as proxy for the ancestral notothenioid state, useful for deciphering evolutionary changes in the Antarctic clade. Our study provides robust whole genome evidence that supports this hypothesis, through generating a long-read, chromosome-level assembly of *E. maclovinus* genome, comparing its genome architecture with the genomes of the basal *C. gobio* and nine derived notothenioids representing all five Antarctic families, and comparing its repertoire of circadian network genes against the same ten genomes.

### 3.1 Genome assembly and annotation

We generated a chromosome-level genome assembly of the Patagonian blennie using PacBio CLR sequencing and Hi-C scaffolding. The final assembly was 606Mb in length, smaller than the 744Mb scaffold-level *E. maclovinus* assembly first described by [20]. The longer length of the original, short read-based assembly could be attributed to its low contiguity (scaffold N50 of 695Kb), likely reflecting unassembled repetitive regions and alternative haplotypes. Our newly generated PacBio assembly is highly contiguous, with a contig and scaffold N50 of 7.6Mb and 26.7Mb, respectively (Table 1). A total of 99.98% of the total sequence length was assembled into 24 chromosome-level scaffolds, in agreement with the 2n=48 karyotype previously described for the species [48]. In addition to its contiguity, the *E. maclovinus* assembly is 96.5% gene complete, according to *BUSCO* v5.3.1 analysis. The combination of high contiguity and gene-completeness indicates the smaller size of the PacBio long-read assembly compared to the short-read *E. maclovinus* assembly is not likely caused by portions of the true genome missing. Instead, the updated PacBio assembly is a better representative of the genome architecture of *E. maclovinus*.

**Table 1.** Assembly and annotation metrics for the E. maclovinus genome

| Assembly Contiguity |  |  |
| --- | --- | --- |
| Scale | Contig | Scaffold |
| Total Size (bp) | 606,099,673 | 606,289,673 |
| Number of Fragments | 406 | 26 |
| Largest Fragment (bp) | 19,993,184 | 37,057,500 |
| N50 (bp) | 7,582,354 | 26,674,500 |
| L50 | 27 | 11 |
| Number of Chromosome-Scale Scaffolds | -- | 24 |
| Total Bases in Chromosomes (bp) | -- | 606,192,173 |
| Percent of Assembly in Chromosomes | -- | 99.98% |
| BUSCO v5.3.1 Gene Completeness |  |  |
| Complete | 3,513 (96.5%) |  |
| Complete & Single Copy | 3,476 (95.5%) |  |
| Complete & Duplicated | 37 (1.0%) |  |
| Fragmented | 12 (0.3%) |  |
| Missing | 115 (3.2%) |  |
| Total | 3,640 |  |
| Reference Gene Set | actinopterygii_odb10 |  |
| Gene Annotation |  |  |
| Number of Protein Coding Genes | 25,081 |  |
| Repeat Annotation |  |  |
| Total Repeats | 33.18% |  |
| LTR | 2.92% |  |
| SINE | 0.64% |  |
| LINE | 5.06% |  |
| DNA | 15.84 |  |
| Unclassified | 5.72% |  |

A total of 25 thousand protein-coding genes were annotated using a combined proteome- and transcriptome-based annotation. This number is comparable with the 20-30 thousand protein-coding genes described in other notothenioid assemblies [21,22,49], and the 23 thousand genes annotated for the Illumina short-read assembly [20]. With regard to repeat content, this assembly is composed of 33.2% repetitive sequences, characterized from a *de novo* repeat library. Nearly half of the repetitive fraction (15.8% of the total genome) was annotated as DNA mobile elements, with the second largest classified fraction (5.1% of the genome) being composed of LINE sequences (Table 1). The 33% repetitive fraction of the genome in *E. maclovinus* is smaller than observed in other cryonotothenioids [21,22,49]. In the context of genome evolution across Notothenioidei, this result reinforces the idea that expansion of repeat elements and associated increase in genome size observed in cryonotothenioids [50] is specific to the Antarctic clade. As the direct extant outgroup of the Antarctic clade, the *E. maclovinus* genome likely reflects the ancestral state of the genome architecture in this group.

### 3.2 Conserved Synteny

Using a comparative synteny analysis, we identified conservation in the large-scale genome organization between basal, non-Antarctic notothenioids and highly derived members of the Antarctic clade. The genomes of both *C. gobio* and *E. maclovinus*, both basal, non-Antarctic species, show conservation in size (∼600Mb) and one-to-one correspondence of 24 orthologous chromosomes (Fig. 1A). A similar relationship is observed when comparing these two basal species to representative members of the Antarctic clade. When compared to *E. maclovinus*, both the red-blooded *D. mawsoni* (Fig. 1A) and the white-blooded *C. gunnari* (Fig. 1B) also exhibit clear correspondence across 24 orthologous chromosomes; however, both species exhibit larger genomes sizes, which are linked to the expansion in repeat elements observed in the Antarctic clade [25,50]. This same relationship, one-to-one correspondence between 24 chromosomes and an increase in genome size in cryonotothenioids, is also observed when we compared the *E. maclovinus* genome against other Antarctic species with chromosome-level assemblies, namely *P. georgianus* and *C. esox*. While changes to the number of chromosomes have previously been observed, including independent evolution of sex chromosomes [51-53] and extensive chromosomal fusions leading to karyotype reduction [54], this one-to-one relationship is largely seen across both basal and derived notothenioid species, and provides further evidence for an ancestral karyotype of 24 chromosomes in the group.

**Figure 1:**
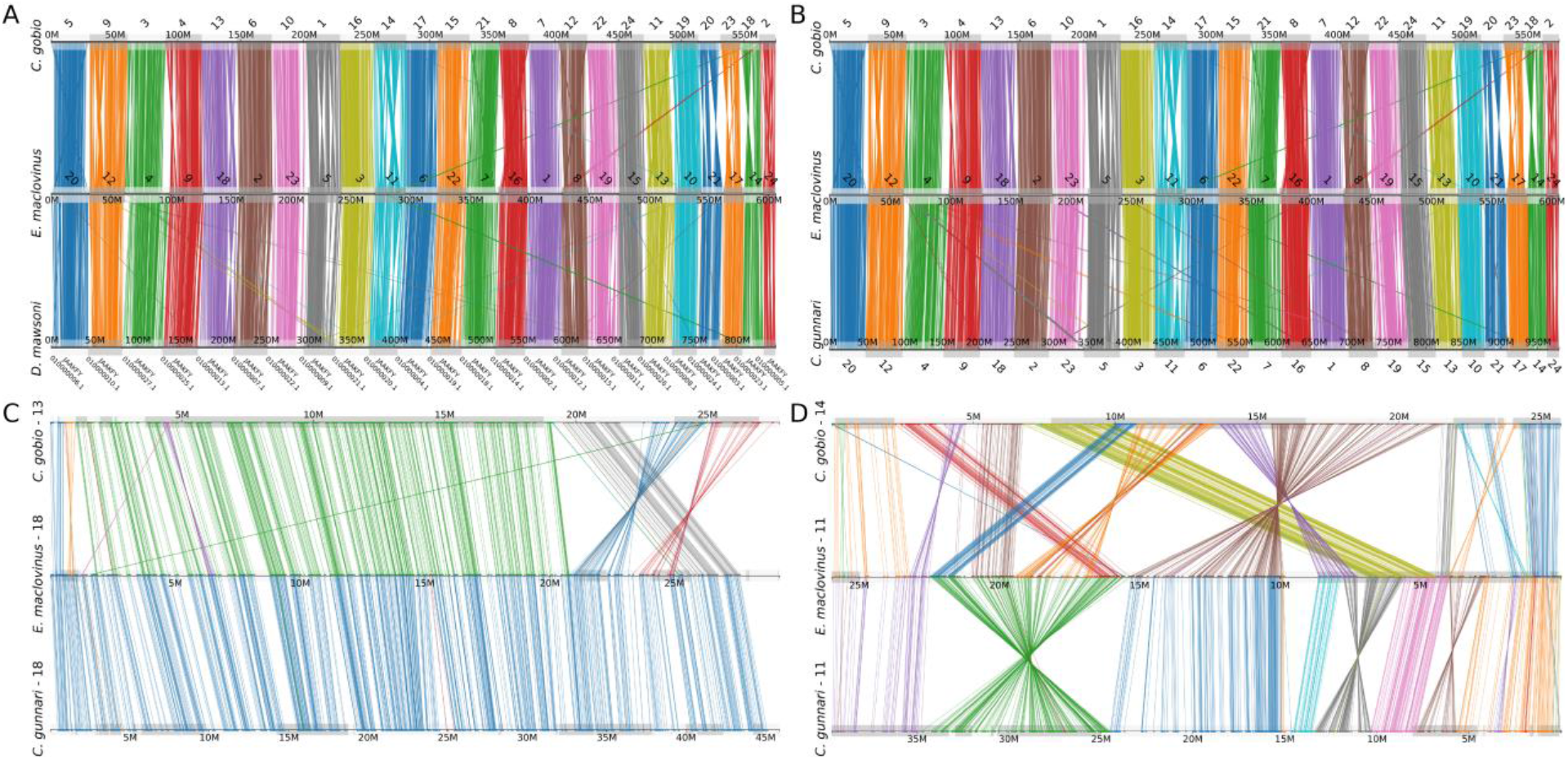
Patterns of conserved synteny across basal temperate and derived Antarctic notothenioid genomes. A) Genome-wide synteny shows a one-to-one correspondence between the 24 chromosomes of the basal temperate *C. gobio* (top), the basal temperate *E. maclovinus* (middle), and the Antarctic red-blooded *D. mawsoni* (bottom). Each line represents an orthologous gene between the compared genomes, color-coded according to their chromosome of origin. B) Genome-wide conserved synteny shows a one-to-one correspondence between the genomes of *C. gobio* (top), *E. maclovinus* (middle), and the Antarctic, white-blooded *C. gunnari* (bottom). Color annotation as described in panel A. C) Conserved synteny between orthologous chromosomes in *C. gobio*-13 (top), *E. maclovinus*-18 (middle), and *C. gunnari*-18 (bottom). Lines represent orthologous genes between the species, color coded according to their respective synteny clusters. Chromosomes are not drawn to scale. Several chromosomal inversions and translocations are observed between *C. gobio* and *E. maclovinus*, while the chromosome organization is fully observed between *E. maclovinus* and *C. gunnari*. D) Conserved synteny between orthologous

Focusing on individual chromosomes, we find evidence for numerous changes in the chromosomal structure between basal and derived notothenioids. Several chromosomal rearrangements, including inversions and translocations, are observed between orthologous chromosomes in *C. gobio* and *E. maclovinus* (Fig. 1C, D). In accordance with their well-recognized phylogenetic placement—in which *E. maclovinus* is the immediate sister taxon to cryonotothenioids [3,55]—we observe a higher degree of conservation in the chromosome structure and organization between *E. maclovinus* and *C. gunnari*, in which synteny is completely conserved along whole chromosomes (Fig. 1C). However, other orthologous chromosomes show extensive evidence of chromosomal rearrangements between these linages (Fig. 1D). This finding indicates that, while genome structure may be largely conserved, there is evidence for the independent evolution of chromosomal organization between basal and Antarctic notothenioids. Nonetheless, given the substantial chromosomal differences observed between *E. maclovinus* and *C. gobio*, and the relatively large degree of conservation between *E. maclovinus* and cryonotothenioid genomes, the *E. maclovinus* assembly may be better suited as an outgroup for future genome-wide comparisons in Antarctic notothenioids. chromosomes in *C. gobio*-14 (top), *E. maclovinus* -11 (middle), and *C. gunnari*-11 (bottom). Color notation as described in panel C. Several examples of chromosomal rearrangements are observed between the three species.

### 3.3 Circadian gene repertoire and expression in E. maclovinus as ancestral reference

In addition to establishing *E. maclovinus* as a better ancestral reference for whole genome architecture evolution, we further evaluate its suitability as reference for addressing functional trait evolution in the Antarctic clade. We analyzed the network of genes that choreograph the circadian rhythm, arguably the most crucial endogenous process guiding the tempo of cellular functioning, systems physiology and whole animal activity throughout life.

*E. maclovinus* and other basal non-Antarctic notothenioids are expected to have a usual circadian rhythm based on their evolutionary distribution in temperate latitudes where diurnal and seasonal light-dark cycles predictably occur. The evolutionary status of the circadian rhythm in Antarctic notothenioids, however, is at present completely unknown. The atypical light-dark regime in the polar region, particularly extreme in the high latitudes where prolonged sea ice cover further reduces, or completely excludes light penetration into marine habitats seasonally, means light-dark zeitgebers, which are major cues for the oscillation of the circadian network, would be irregular or protractedly absent for the local fishes.

Leveraging publicly available chromosomal or near-chromosomal scale genomes of 10 notothenioids, and *E. maclovinus* genome in this study, and using a set of 33 circadian network genes from the reference teleosts *D. rerio* and *G. aculeatus*, we curated the orthologs from these 11 notothenioid species that represent seven of the eight families of the Notothenioidei suborder (Table S2). Fig. 2B summarizes the curated circadian gene orthologs found in the notothenioids, and contrasts gene presence and absence relative to the teleost reference set, and between the basal species (*C. gobio* and *E. maclovinus*) and the nine derived cryonotothenioids.

**Figure 2.**
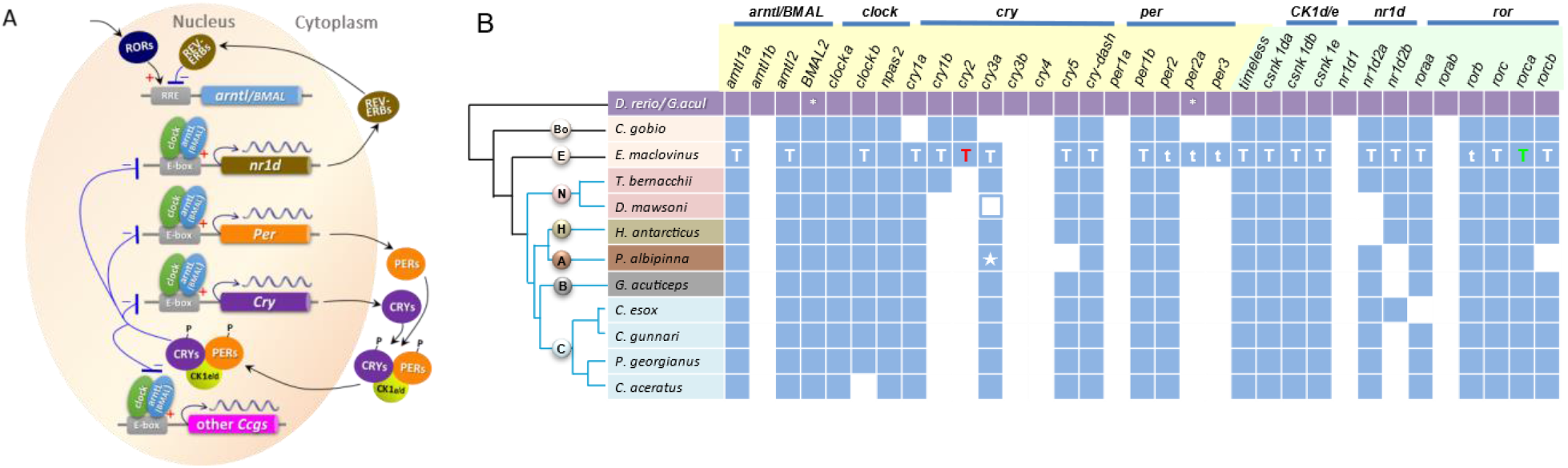
Schematic of circadian regulatory loops and the circadian network genes identified from the genome and transcriptomes of *E. maclovinus* and from genomes across Notothenioidei. **A)**. A simplified depiction of the circadian regulatory loops based on mammalian studies [59,60] relating the positions of the orthologous genes identified from notothenioids in the circadian network. Briefly, the activator core clock proteins arntl/BMAL (**a**ryl hydrocarbon **r**eceptor **n**uclear **t**ranslocator-**l**ike and **b**rain and **m**uscle **a**rnt-**l**ike are alternate names) and clock (**c**ircadian **l**ocomotor **o**utput **c**ycles protein **k**aput) heterodimerize and bind to regulatory elements containing E-box sequences of a large number of clock controlled genes (*Ccgs*) including the repressor genes *cry* and *per*. The cry and per proteins accumulate in the cytoplasm, interact with each other and the serine/threonine casein kinases *CK1e* and *CK1d*. The complex translocates back into the nucleus, and suppresses target gene activation by arntl/BMAL and clock, leading to their own transcriptional repression. *Timeless* protein (not shown) also interacts with *per* proteins and others to suppress arntl/BMAL activation of per1. The regulation of *arntl/BMAL* and *clock* transcription is in turn effected through a loop involving nuclear receptor proteins REV-ERB and ROR. Activation of *nr1d* by *arntl/BMAL* and *clock*, and of *ROR* through an intervening activator, produce REV-ERB (encoded by *nr1d* paralogs) and ROR proteins respectively. These compete for their common binding sequence element RORE in *arntl/BMAL* promotor, where ROR initiates and REV-ERB inhibits *arntl/BMAL* transcription. **B)**. Evolutionary status of circadian network genes in genomes of notothenioid fishes identified using 33 reference genes from zebrafish *D. rerio* and three-spined stickleback *G. aculeatus*. All reference genes (violet squares) except two are from *D. rerio*, with * indicating *G. aculeatus* as the source since it is absent in *D. rerio*. Names of the core loop circadian genes are highlighted in light yellow, and the secondary loop rhythmic genes in light green, with paralogs of each gene family grouped under a blue bar. Light blue squares indicate presence of notothenioid orthologs in the genome annotation of each species determined with *SYNOLOG*, and uncolored (white) squares indicate gene absence. A cladogram on the left depicts the phylogenetic relationship of the included notothenioid species, which represent seven of the eight families – Bo (Bovichtidae), E (Eleginopidae), N (Nototheniidae), H (Harpagiferidae), A (Artedidraconidae), B (Bathydraconidae), and C (Channichthyidae). *SYNOLOG* did not detect *cry3* in *D. mawsoni* or *P. albipinna*. Its presence in these two species was determined by manual BLAST search and annotation, which found *D. mawsoni cry3* (blue outlined white square) gene structure to be conserved except for two 1-nt frameshift mutations, and *P. albipinna cry3* (blue square with white star) to be complete and mutation-free. Circadian gene repertoire in the two basal non-Antarctic notothenioids differ, with *E. maclovinus* retaining more of the reference teleost set, while apparent *C. gobio*-specific loss of two *cry* (*cry1a, cry3a)* and two *per (per2a, per3*) paralogs had occurred since the two species diverged. Transcript evidence supporting functionality of the circadian network genes in *E. maclovinus* indicated by the letter “T” or “t” are as follows: T (white) – Iso-Seq transcript, complete cds; T (red) – complete cds from overlapping partial Iso-Seq + partial Trinity assembled RNAseq transcripts; T (green) – overlapping Trinity transcripts, complete cds; t (white) -partial Trinity assembled RNAseq transcript.

The 33 reference circadian genes/paralogs were selected based on their known function in the widely studied circadian system in Drosophila [56] and in mammals [57,58], and that orthologs are present in teleost fish. Fig. 2A provides a simplified depiction of the autonomous cellular transcription-translation negative feedback regulatory loops underlying circadian rhythmicity, and the positions of the 33 genes/paralogs in the circadian process. The core loop comprises the phasic interactions of the activator proteins of *arntl/BMAL* and *clock* with the suppressor proteins of *cry and per* paralogs (gene names highlighted in light yellow in Fig. 2B). Secondary loops involve *CK1e/CK1d*, and the nuclear receptors *nr1d* and *ror* paralogs (gene names highlighted in light green in Fig. 2B), whose proteins regulate the core clock genes. *Timeless* protein (not shown in Fig. 2A) also participates in a secondary loop, interacting with *per* proteins and others to suppress arntl/BMAL activation of *per*.

Relative to the teleost reference set, several paralogs of the core loop clock genes, namely *arntl1b, cry3b, cry4*, and *per1a*, as well as *nr1d1* and *rorab* of the secondary loops were absent across all basal and derived notothenioids (Fig. 2B), suggesting these six genes were absent in the common ancestor of the suborder. Within notothenioids, the two basal lineages differ in their circadian gene repertoire. The maintained set of genes (27) in *E. maclovinus* is similar to the teleost reference, while *C. gobio* has apparently undergone lineage-specific loss of additional *cry* (*cry1a, cry3a*) and *per* (*per2a, per3*) paralogs since the two basal taxa diverged. Between *E. maclovinus* and the derived cryonotothenioid species, the core clock genes *cry1b, cry2* have undergone Antarctic-specific loss. *Per2a* and *per3* were also lost in the cryonotothenioids, separately from their loss in *C. gobio*, since they are present in *E. maclovinus* the closest sister taxon to the Antarctic clade.

The remaining core clock genes are largely conserved between *E. maclovinus* and the derived cryonotothenioids. However, the *cryptochrome* paralog *cry3a* was not detected by *SYNOLOG* in *D. mawsoni* (nototheniid) or *P. albipinna* (artedidraconid) genome, suggesting varied losses in the Antarctic clade, or inaccuracy in the current state of the genome annotations of these two species. By manual *BLAST* searches we recovered a full length copy of *cry3a* from both species (Fig. S2). In manual annotation, we found *D. mawsoni cry3a* (in HiC_scaffold_10) to be completely conserved in gene structure, but contains a 1-nt indel in each of two exons, which would result in a frameshifted protein gene model with little sequence similarity to intact cry3a, thus eluding detection by *SYNOLOG. Cry3a* in the artedidraconid *P. albipinna* (in scaffold CM053253.1) however, is completely mutation free, and thus a structurally intact gene (Fig. S2). It has eluded detection likely because the current annotation has split up the gene into two halves, each with hypothetical protein as annotation. With the manual recovery of *D. mawsoni* and *P. albipinna cry3a*, it appears likely that this core clock gene paralog is preserved across the Antarctic clade. In contrast, manual search for *cry3a* in *C. gobio* draft genome only found an orphan exon 1 in an unplaced scaffold, verifying definitive loss.

Presence of the rhythmic genes of the secondary regulatory loops -*timeless*, and paralogs of *CK1d/e* (casein kinase 1) and *ror* (retinoic-acid-related orphan receptors) is conserved throughout notothenioids (Fig. 2B). In contrast, the two paralogs of *nr1d2* (nuclear receptor subfamily 1 group D) appear to have undergone varied loss in different Antarctic lineages. This *nr1d* evolutionary pattern is difficult to interpret, and how much bioinformatics accuracy play a part remains to be evaluated.

Curation of circadian network genes from a genome assembly serves to catalog the set of potentially active contributors to circadian rhythm generation. To ascertain functionality of these predicted genes requires transcriptional evidence at minimal. We thus obtained high-quality long read transcripts by PacBio Iso-Seq sequencing of fin tissue, where peripheral clock is known to operate in teleost fishes [46,47]. Where Iso-Seq transcript is missing for a given gene, we searched the multi-tissue reference transcriptome assembled from published Illumina RNAseq short reads [15].

In total, we found mRNA expression for 24 of the 27 predicted circadian genes/paralogs in *E. maclovinus* genome (Fig. 2B; sequences in File S1). Of the 24 expressed genes, 20 are supported by full-length cDNAs (complete coding sequences) in the transcriptomes. They are indicated with uppercase letter T (Fig. 2B) as follows: 18 (white letter T) are full-length Iso-Seq transcripts, one (red letter T) full-length is from overlapping partial Iso-Seq transcripts and partial transcripts from the RNAseq reference transcriptome, and one (orange letter T) full-length is from overlapping partial transcripts from the RNAseq reference transcriptome. The other four (indicated with white lowercase t) are three *per* paralogs (*per2, per2a, per3*) and one *ror* paralog (*rorb*), with expression evidence as partial transcripts in the RNAseq reference transcriptome.

Transcripts of the remaining three *E. maclovinus* clock genes – *BMAL2, clocka* and *npas2*-are lacking in both the Iso-Seq and RNAseq transcriptome. Their absence can either be due to the genes being inactive, or their mRNA not being represented in the RNA sample that was sequenced. *Npas2* (neuronal PAS domain Protein 2) is a *clock* paralog with demonstrated expression in teleost brain [61,62]. Whether it is expressed in a peripheral tissue such as fin is unknown, thus its absence in the fin Iso-Seq transcriptome may be due to tissue specificity of expression. The RNAseq reference transcriptome did include whole brain in the pooled RNA sample [15], but its small fractional input (1 of 11 tissues) might not have yielded sufficient sequence coverage to encompass the full complement of circadian network genes. An additional constraint in capturing the full circadian transcriptome is the interlocking phasic up- and down-regulation of gene expression in the different regulatory loops, such that genes that are transcriptionally activated at the time of tissue sampling are more likely to be captured, while those that are repressed or undergoing degradation are not. The tissues used for the two transcriptomes came from the same individuals sampled over an afternoon for the purpose of the published study [15]. To definitively ascertain transcriptional functionality of *BMAL2, clocka* and *npas2* will require using RNA from tissues (both fin and brain) sampled at regular intervals over a complete (24 hour) circadian time period for additional sequencing.

In sum, the *E. maclovinus* genome contains orthologs of 27 of the 33 reference circadian network genes/paralogs of model teleosts, with gene functionality of 24 validated by transcript evidence predominantly in full length Iso-Seq cDNAs. The five missing circadian genes are also absent in the more basally divergent bovichtid *C. gobio*, reflecting their loss in the common ancestor of Notothenioidei. *C. gobio* further lost *cry1a* and *cry3a*, and would be reliant on *cry1b* and *cry2* as the participating *cryptochrome* paralogs in the core regulatory loop. In contrast, functional *cry1a* and *cry3a* are maintained in *E. maclovinus* and gene models are present across species of the Antarctic clade. In the latter, *cry1a* and *cry3a* would likely become the core loop repressors, as *cry1b* and *cry2* were lost after their divergence from *E. maclovinus*. Overall, *E. maclovinus* and cryonotothenioids shares more of the same retained circadian genes than between the latter with *C. gobio*. This supports *E. maclovinus* as a better candidate as ancestral reference for investigating circadian and other functional trait changes during polar evolution of cryonotothenioids. It can also serve as reference for inquiries into potential changes in the re-colonization of non-freezing environment by secondarily temperate notothenioids.

## 4. Conclusion

The Patagonia blennie *E. maclovinus* occupies a strategically important phylogenetic position in Notothenioidei, as the monotypic species of the basal, non-Antarctic family Eleginopidae and immediate sister lineage to the Antarctic clade of cryonotothenioids. Using PacBio CLR sequencing and HiC scaffolding, our study provided a highly contiguous and gene-complete chromosomal-level assembly for this species. Through comparative whole genome architecture analyses with the genomes of ten other notothenioids, substantial structural divergence can be seen between the two basal species *C. gobio* and *E. maclovinus*, while there is a larger degree of conservation between *E. maclovinus* and derived cryonotothenioids. We documented a repertoire of 27 circadian network genes in the *E. maclovinus* genome and provided expression evidence supporting active transcription for most of them. Comparison of the pattern of circadian gene/paralog retention and loss among the same 10 species used for genome structure analysis, again illustrates greater similarity between and the cryonotothenioid than between *C. gobio* and the latter. These results further reinforce the suitability of *E. maclovinus* as ancestral proxy in investigating genome and functional trait evolution in the derived Antarctic clade during its evolution in extreme polar conditions. We additionally suggest that *E. maclovinus* can also serve as comparison in understanding the nature of the evolutionary path and mechanisms underlying secondary adaptation to non-freezing environments of various cryonotothenioid lineages that have undergone polar to temperate transition.

## Supplementary Materials

The following supporting information can be downloaded at: www.mdpi.com/xxx/s1, Figure S1: Conserved genome-wide synteny between *E. maclovinus* and the platyfish, *X. maculatus*; Figure S2: Manually curated and annotated *cry3a* from *D. mawsoni* and *P. albipinna* genome; Table S1: Genome assemblies used for conserved synteny analysis and curation of circadian genes; Table S2: Reference circadian genes from model teleosts zebrafish and threespine stickleback, and their orthologs in 11 notothenioid genomes with genomic locations indicated. File S1: Transcript sequences of circadian genes in *E. maclovinus* obtained from Iso-Seq and RNAseq transcriptomes.

## Author Contributions

Conceptualization, C.-H.C.C. and J.M.C.; methodology, formal analysis, investigation, C.-H.C.C., A.G.R.-C., B. F. M., L.W., N.R., J.M.C.; resources, C.-H.C.C., L.V.-C.; data curation, C.-H.C.C., A.G.R.-C.; writing—original draft preparation, C.-H.C.C. and A.G.R.-C.; writing—review and editing, C.-H.C.C. and A.G.R.-C.; supervision, project administration, funding acquisition, C.-H.C.C. and J.M.C. All authors have read and agreed on the manuscript.

## Funding

This research was funded by the United States National Science Foundation grant 1645087 to J.M.C. and C.-H.C.C., and NSF ANT Grant 11-42158 to C.-H.C.C. and Arthur L. DeVries. L.W. was supported by the University of Illinois Foundation Cheng-DeVries Research Fund. L.V.-C. was supported by Fondap-IDEAL 15150003 and ANID-Millennium Science Initiative Program-Center ICM-ANID ICN2021_002.

## Institutional Review Board Statement

The animal use in the study was approved by the Institutional Animal Care and Use Committee (IACUC) of University of Illinois, Urbana-Champaign on April 5, 2007, for Protocol no. 07053, and on June 11, 2012, for Protocol no.12123, and by Comité de Bioética Animal, Universidad Austral de Chile on August 2, 2010, for Protocol No 11/10.

## Data Availability Statement

The raw PacBio and Hi-C Illumina reads used for the *E. maclovinus* genome assembly are available on NCBI under BioProject PRJNA917608. A copy of the *E. maclovinus* reference assembly and annotation, alongside the gene models and annotation for all studied notothenioid species, and copies of all bioinformatic scripts used for analysis are available on DRYAD (https://doi.org/10.5061/dryad.qbzkh18nt).

## Acknowledgments

We sincerely thank Arthur L. DeVries for his assistance in sampling experimental specimens of *E. maclovinus* used in the study.

## Conflicts of Interest

The authors declare no conflict of interest.

## Notes

https://doi.org/10.5061/dryad.qbzkh18nt

### Competing Interest Statement

The authors have declared no competing interest.

## References

1. Eastman, J.T. The nature of the diversity of Antarctic fishes. Polar Biol. 2005, 28, 94–107, doi:10.1007/s00300-004-0667-4.

2. Matschiner, M.; Hanel, R.; Salzburger, W. On the Origin and Trigger of the Notothenioid Adaptive Radiation. PLoS One 2011, 6, e18911, doi:doi.org/10.1371/journal.pone.0018911.

3. Near, T.J.; Dornburg, A.; Kuhn, K.L.; Eastman, J.T.; Pennington, J.N.; Patarnello, T.; Zane, L.; Fernández, D.A.; Jones, C.D. Ancient climate change, antifreeze, and the evolutionary diversification of Antarctic fishes. Proc. Natl. Acad. Sci. U.S.A. 2012, 109, 3434–3439, doi:10.1073/pnas.1115169109.

4. Eastman, J.T.; McCune, A.R. Fishes on the Antarctic continental shelf: evolution of amarine species flock?*. J. Fish. Biol. 2000, 57, 84–102, doi:https://doi.org/10.1111/j.1095-8649.2000.tb02246.x.

5. Lecointre, G.; Améziane N Fau - Boisselier, M.-C.; Boisselier Mc Fau - Bonillo, C.; Bonillo C Fau - Busson, F.; Busson F Fau - Causse, R.; Causse R Fau - Chenuil, A.; Chenuil A Fau - Couloux, A.; Couloux A Fau - Coutanceau, J.-P.; Coutanceau Jp Fau - Cruaud, C.; Cruaud C Fau - d’Acoz, C.d.U.; et al. Is the species flock concept operational? The Antarctic shelf case. PLoS One 2013, 8, e68787, doi:10.1371/journal.pone.0068787.

6. Chen, L.; DeVries, A.L.; Cheng, C.-H.C. Evolution of antifreeze glycoprotein gene from a trypsinogen gene in Antarctic notothenioid fish. Proc. Natl. Acad. Sci. USA 1997, 94, 3811–3816, doi:doi.org/10.1073/pnas.94.8.3811.

7. Cheng, C.-H.C.; Zhuang, X. Molecular Origins and Mechanisms of Fish Antifreeze Evolution. In Antifreeze Proteins Volume 1: Environment, Systematics and Evolution, Ramløv, H., Friis, D.S., Eds.; Springer International Publishing: Cham, 2020; pp. 275–313. doi.org/10.1007/978-3-030-41929-5_9

8. DeVries, A.L. Glycoproteins as biological antifreeze agents in Antarctic fishes. Science 1971, 172, 1152–1155., doi:10.1126/science.172.3988.1152.

9. DeVries, A.L.; Cheng, C.-H.C. Antifreeze proteins and organismal freezing avoidance in polar fishes. In The physiology of polar fishes, Farrell, A.P., Steffensen, J.F., Eds.; Elsevier Academic Press: San Diego, 2005; Volume 22, pp. 155–201. doi.org/10.1016/S1546-5098(04)22004-0

10. Todgham, A.E.; Hoaglund, E.A.; Hofmann, G.E. Is cold the new hot? Elevated ubiquitin-conjugated protein levels in tissues of Antarctic fish as evidence for cold-denaturation of proteins in vivo. J. Comp. Physio. B 2007, 177, 857–866, doi:doi: 10.1007/s00360-007-0183-2.

11. Beers, J.M.; Jayasundara, N. Antarctic notothenioid fish: what are the future consequences of ‘losses’ and ‘gains’ acquired during long-term evolution at cold and stable temperatures? J. Exp. Biol. 2015, 218, 1834–1845, doi:10.1242/jeb.116129.

12. Daane, J.M.; Detrich, H.W. Adaptations and Diversity of Antarctic Fishes: A Genomic Perspective. Ann. Rev. Animal Biosci. 2022, 10, 39–62, doi:10.1146/annurev-animal-081221-064325.

13. Bilyk, K.T.; DeVries, A.L. Heat tolerance and its plasticity in Antarctic fishes. Comp Biochem Physiol A Mol Integr Physiol 2011, 158, 382–390, doi:10.1016/j.cbpa.2010.12.010.

14. Somero, G.N.; DeVries, A.L. Temperature tolerance of some Antarctic fishes. Science 1967, 156, 257–258. doi: 10.1126/science.156.3772.257

15. Bilyk, K.T.; Vargas-Chacoff, L.; Cheng, C.H.C. Evolution in chronic cold: varied loss of cellular response to heat in Antarctic notothenioid fish. BMC Evol Biol 2018, 18, 143, doi:10.1186/s12862-018-1254-6.

16. Hofmann, G.E.; Buckley, B.A.; Airaksinen, S.; Keen, J.; Somero, G.N. The Antarctic fish Trematomus bernacchii lacks heat-inducible heat shock protein synthesis. J. Exp. Biol. 2000, 203, 2331–2339, doi:doi.org/10.1242/jeb.203.15.2331.

17. Saravia, J.; Paschke, K.; Oyarzún-Salazar, R.; Cheng, C.H.C.; Navarro, J.M.; Vargas-Chacoff, L. Effects of warming rates on physiological and molecular components of response to CTMax heat stress in the Antarctic fish Harpagifer antarcticus. J. Thermal Biol. 2021, 99, 103021, doi:https://doi.org/10.1016/j.jtherbio.2021.103021.

18. di Prisco, G.; Cocca, E.; Parker, S.K.; Detrich, H.W., III. Tracking the Evolutionary Loss of Hemoglobin Expression by the White-blooded Antarctic Icefishes. Gene 2003, 295, 185–191, doi:10.1016/s0378-1119(02)00691-1.

19. Sidell, B.D.; O’Brien, K.M. When bad things happen to good fish: the loss of hemoglobin and myoglobin expression in Antarctic icefishes. J. Exp. Biol. 2006, doi:https://doi.org/10.1242/jeb.02091.

20. Chen, L.; Lu, Y.; Li, W.; Ren, Y.; Yu, M.; Jiang, S.; Fu, Y.; Wang, J.; Peng, S.; Bilyk, K.T.; et al. The genomic basis for colonizing the freezing Southern Ocean revealed by Antarctic toothfish and Patagonian robalo genomes. GigaScience 2019, 8, doi:10.1093/gigascience/giz016.

21. Bista, I.; Wood, J.M.D.; Desvignes, T.; McCarthy, S.A.; Matschiner, M.; Ning, Z.; Tracey, A.; Torrance, J.; Sims, Y.; Chow, W.; et al. Genomics of cold adaptations in the Antarctic notothenioid fish radiation. bioRxv 2022, doi:10.1101/2022.06.08.494096.

22. Rivera-Colón, A.G.; Rayamajhi, N.; Minhas, B.F.; Madrigal, G.; Bilyk, K.T.; Yoon, V.; Hüne, M.; Gregory, S.; Cheng, C.H.C.; Catchen, J.M. Genomics of Secondarily Temperate Adaptation in the Only Non-Antarctic Icefish. Mol. Biol. Evol. 2023, 40, msad029, doi:10.1093/molbev/msad029.

23. Bilyk, K.T.; Zhuang, X.; Papetti, C. Positive and Relaxed Selective Pressures Have Both Strongly Influenced the Evolution of Cryonotothenioid Fishes during Their Radiation in the Freezing Southern Ocean. Genome Biology and Evolution 2023, 15, evad049, doi:10.1093/gbe/evad049.

24. Lu, Y.; Li, W.; Li, Y.; Zhai, W.; Zhou, X.; Wu, Z.; Jiang, S.; Liu, T.; Wang, H.; Hu, R.; et al. Population genomics of an icefish reveals mechanisms of glacier-driven adaptive radiation in Antarctic notothenioids. BMC Biology 2022, 20, 231, doi:10.1186/s12915-022-01432-x.

25. Bista, I.; McCarthy, S.A.; Wood, J.; Ning, Z.; Detrich III, H.W.; Desvignes, T.; Postlethwait, J.; Chow, W.; Howe, K.; Torrance, J.; et al. The genome sequence of the channel bull blenny, Cottoperca gobio (Günther, 1861). Wellcome Open Research 2020, 5, 148, doi:10.12688/wellcomeopenres.16012.1.

26. Bilyk, K.T.; Zhuang, X.; Vargas-Chacoff, L.; Cheng, C.H.C. Evolution of chaperome gene expression and regulatory elements in the antarctic notothenioid fishes. Heredity 2021, 126, 424–441, doi:10.1038/s41437-020-00382-w.

27. Rayamajhi, N.; Cheng, C.-H.C.; Catchen, J.M. Evaluating Illumina-, Nanopore-, and PacBio-based genome assembly strategies with the bald notothen, Trematomus borchgrevinki. G3 Genes|Genomes|Genetics 2022, jkac192, doi:10.1093/g3journal/jkac192.

28. Ruan, J.; Li, H. Fast and accurate long-read assembly with wtdbg2. Nature Methods 2020, 17, 155–158, doi:10.1038/s41592-019-0669-3.

29. Gurevich, A.; Saveliev, V.; Vyahhi, N.; Tesler, G. QUAST: quality assessment tool for genome assemblies. Bioinformatics 2013, 29, 1072–1075, doi:10.1093/bioinformatics/btt086.

30. Simão, F.A.; Waterhouse, R.M.; Ioannidis, P.; Kriventseva, E.V.; Zdobnov, E.M. BUSCO: assessing genome assembly and annotation completeness with single-copy orthologs. Bioinformatics 2015, 31, 3210–3212, doi:10.1093/bioinformatics/btv351.

31. Durand, N.C.; Shamim, M.S.; Machol, I.; Rao, S.S.P.; Huntley, M.H.; Lander, E.S.; Aiden, E.L. Juicer Provides a One-Click System for Analyzing Loop-Resolution Hi-C Experiments. Cell Systems 2016, 3, 95–98, doi:10.1016/j.cels.2016.07.002.

32. Manni, M.; Berkeley, M.R.; Seppey, M.; Simão, F.A.; Zdobnov, E.M. BUSCO Update: Novel and Streamlined Workflows along with Broader and Deeper Phylogenetic Coverage for Scoring of Eukaryotic, Prokaryotic, and Viral Genomes. Mol. Biol. Evol. 2021, 38, 4647–4654, doi:10.1093/molbev/msab199.

33. Flynn, J.M.; Hubley, R.; Goubert, C.; Rosen, J.; Clark, A.G.; Feschotte, C.; Smit, A.F. RepeatModeler2 for automated genomic discovery of transposable element families. Proc. Natl. Acad. Sci. U.S.A. 2020, 117, 9451–9457, doi:10.1073/pnas.1921046117.

34. Ellinghaus, D.; Kurtz, S.; Willhoeft, U. LTRharvest, an efficient and flexible software for de novo detection of LTR retrotransposons. BMC Bioinformatics 2008, 9, 18, doi:10.1186/1471-2105-9-18.

35. Smit, A.F.; Hubley, R.; Green, P. RepeatMasker Open-4.0, 2013.

36. Dobin, A.; Davis, C.A.; Schlesinger, F.; Drenkow, J.; Zaleski, C.; Jha, S.; Batut, P.; Chaisson, M.; Gingeras, T.R. STAR: ultrafast universal RNA-seq aligner. Bioinformatics 2013, 29, 15–21, doi:10.1093/bioinformatics/bts635.

37. Brůna, T.; Hoff, K.J.; Lomsadze, A.; Stanke, M.; Borodovsky, M. BRAKER2: automatic eukaryotic genome annotation with GeneMark-EP+ and AUGUSTUS supported by a protein database. NAR Genomics and Bioinformatics 2021, 3, lqaa108, doi:10.1093/nargab/lqaa108.

38. Hoff, K.J.; Lange, S.; Lomsadze, A.; Borodovsky, M.; Stanke, M. BRAKER1: Unsupervised RNA-Seq-Based Genome Annotation with GeneMark-ET and AUGUSTUS: Table 1. Bioinformatics 2016, 32, 767–769, doi:10.1093/bioinformatics/btv661.

39. Kriventseva, E.V.; Kuznetsov, D.; Tegenfeldt, F.; Manni, M.; Dias, R.; Simão, F.A.; Zdobnov, E.M. OrthoDB v10: sampling the diversity of animal, plant, fungal, protist, bacterial and viral genomes for evolutionary and functional annotations of orthologs. Nucleic Acids Res. 2019, 47, D807–D811, doi:10.1093/nar/gky1053.

40. Gabriel, L.; Hoff, K.J.; Brůna, T.; Borodovsky, M.; Stanke, M. TSEBRA: transcript selector for BRAKER. BMC Bioinformatics 2021, 22, 566, doi:10.1186/s12859-021-04482-0.

41. Catchen, J.M.; Conery, J.S.; Postlethwait, J.H. Automated identification of conserved synteny after whole-genome duplication. Genome Res. 2009, 19, 1497–1505, doi:10.1101/gr.090480.108.

42. Small, C.M.; Bassham, S.; Catchen, J.; Amores, A.; Fuiten, A.M.; Brown, R.S.; Jones, A.G.; Cresko, W.A. The genome of the Gulf pipefish enables understanding of evolutionary innovations. Genome Biol 2016, 17, 258, doi:10.1186/s13059-016-1126-6.

43. Jae Lee, S.; Kim, J.H.; Jo, E.; Choi, E.; Kim, J.; Choi, S.G.; Chung, S.; Kim, H.W.; Park, H. Chromosomal assembly of the Antarctic toothfish (Dissostichus mawsoni) genome using third-generation DNA sequencing and Hi-C technology. Zool. Res. 2021, 42, 124–129, doi:10.24272/j.issn.2095-8137.2020.264.

44. Camacho, C.; Coulouris, G.; Avagyan, V.; Ma, N.; Papadopoulos, J.; Bealer, K.; Madden, T.L. BLAST+: architecture and applications. BMC Bioinformatics 2009, 10, 421, doi:10.1186/1471-2105-10-421.

45. Amores, A.; Catchen, J.; Nanda, I.; Warren, W.; Walter, R.; Schartl, M.; Postlethwait, J.H. A RAD-Tag Genetic Map for the Platyfish (Xiphophorus maculatus) Reveals Mechanisms of Karyotype Evolution Among Teleost Fish. Genetics 2014, 197, 625–641, doi:10.1534/genetics.114.164293.

46. Beale, A.; Guibal, C.; Tamai, T.K.; Klotz, L.; Cowen, S.; Peyric, E.; Reynoso, V.H.; Yamamoto, Y.; Whitmore, D. Circadian rhythms in Mexican blind cavefish Astyanax mexicanus in the lab and in the field. Nature Commun. 2013, 4, 2769, doi:10.1038/ncomms3769.

47. Foulkes, N.S.; Whitmore, D.; Vallone, D.; Bertolucci, C. Chapter One - Studying the Evolution of the Vertebrate Circadian Clock: The Power of Fish as Comparative Models. In Advances in Genetics, Foulkes, N.S., Ed.; Academic Press: 2016; Volume 95, pp. 1–30.

48. Mazzei, F.; Ghigliotti, L.; Coutanceau, J.-P.; Detrich, H.W.; Prirodina, V.; Ozouf-Costaz, C.; Pisano, E. Chromosomal characteristics of the temperate notothenioid fish Eleginops maclovinus (Cuvier). Polar Biology 2008, 31, 629, doi:10.1007/s00300-007-0399-3.

49. Kim, B.-M.; Amores, A.; Kang, S.; Ahn, D.-H.; Kim, J.-H.; Kim, I.-C.; Lee, J.H.; Lee, S.G.; Lee, H.; Lee, J.; et al. Antarctic blackfin icefish genome reveals adaptations to extreme environments. Nat Ecol Evol 2019, 3, 469–478, doi:10.1038/s41559-019-0812-7.

50. Auvinet, J.; Graça, P.; Belkadi, L.; Petit, L.; Bonnivard, E.; Dettaï, A.; Detrich, W.H.; Ozouf-Costaz, C.; Higuet, D. Mobilization of retrotransposons as a cause of chromosomal diversification and rapid speciation: the case for the Antarctic teleost genus Trematomus. BMC Genomics 2018, 19, 339, doi:10.1186/s12864-018-4714-x.

51. Ghigliotti, L.; Cheng, C.-H.C.; Pisano, E. Sex determination in Antarctic notothenioid fish: chromosomal clues and evolutionary hypotheses. Polar Biology 2016, 39, 11–22, doi:10.1007/s00300-014-1601-z.

52. Morescalchi, A.; Hureau, J.C.; Olmo, E.; Ozouf-Costaz, C.; Pisano, E.; Stanyon, R. A multiple sex-chromosome system in Antarctic ice-fishes. Polar Biology 1992, 11, doi:10.1007/BF00237962.

53. Morescalchi, A.; Pisano, E.; Stanyon, R.; Morescalchi, M.A. Cytotaxonomy of antarctic teleosts of the Pagothenia/Trematomus complex (Nototheniidae, Perciformes). Polar Biology 1992, 12, doi:10.1007/BF00236979.

54. Amores, A.; Wilson, C.A.; Allard, C.A.H.; Detrich, H.W.; Postlethwait, J.H. Cold Fusion: Massive Karyotype Evolution in the Antarctic Bullhead Notothen Notothenia coriiceps. G3 Genes|Genomes|Genetics 2017, 7, 2195–2207, doi:10.1534/g3.117.040063.

55. Lecointre, G.; Bonillo, C.; Ozouf-Costaz, C.; Hureau, J.C. Molecular evidence for the origins of Antarctic fishes: paraphyly of the Bovichtidae and no indication for the monophyly of the Notothenioidei (Teleostei). Polar Biology 1997, 18, 193–208, doi:10.1007/s003000050176.

56. Scully, A.L.; Kay, S.A. Time Flies for Drosophila. Cell 2000, 100, 297–300, doi:https://doi.org/10.1016/S0092-8674(00)80665-0.

57. Minami, Y.; Ode, K.L.; Ueda, H.R. Mammalian Circadian Clock: The Roles of Transcriptional Repression and Delay. In Circadian Clocks, Kramer, A., Merrow, M., Eds.; Springer Berlin Heidelberg: Berlin, Heidelberg, 2013; pp. 359–377.

58. Cox, K.H.; Takahashi, J.S. Circadian clock genes and the transcriptional architecture of the clock mechanism. J. Mol. Endocrinol. 2019, 63, R93–R102, doi:10.1530/JME-19-0153.

59. Buhr, E.D.; Takahashi, J.S. Molecular Components of the Mammalian Circadian Clock. In Circadian Clocks, Kramer, A., Merrow, M., Eds.; Springer Berlin Heidelberg: Berlin, Heidelberg, 2013; pp. 3–27.

60. Takahashi, J.S. Transcriptional architecture of the mammalian circadian clock. Nat. Rev. Genet. 2017, 18, 164–179, doi:10.1038/nrg.2016.150.

61. Bolton, C.M.; Bekaert, M.; Eilertsen, M.; Helvik, J.V.; Migaud, H. Rhythmic Clock Gene Expression in Atlantic Salmon Parr Brain. Front. Physiol. 2021, 12, 761109, doi:doi: 10.3389/fphys.2021.761109.

62. Martins, R.S.T.; Gomez, A.; Zanury, S.; Carrillo, M.; Canario, A.V.M. Photoperiodic Modulation of Circadian Clock and Reproductive Axis Gene Expression in the Pre-Pubertal European Sea Bass Brain. PLoS One 2015, 0144158, doi:/doi.org/10.1371/journal.pone.0144158.

