## Supplemental Figures and File for "Chromosome-Level Genome Assembly and Circadian Gene Repertoire of the Patagonia blennie *Eleginops maclovinus* - the closest ancestral proxy of Antarctic cryonotothenioids"

### Supplementary Figures

**Figure S1: Conserved genome-wide synteny between *E. maclovinus* and the platyfish, *X. maculatus*.** Genome-wide conserved synteny plot displaying orthology between the 24 chromosomes of *X. maculatus* (top) and *E. maclovinus* (bottom). Each ortholog gene pair between the two species is represented by a line color-coded according to their chromosome of origin. While synteny highlights multiple intra-chromosomal rearrangements, chromosomes largely display one-to-one correspondence between the two species. In turn, the *E. maclovinus* chromosomal scaffolds were named according to their orthologous *X. maculatus* chromosome.

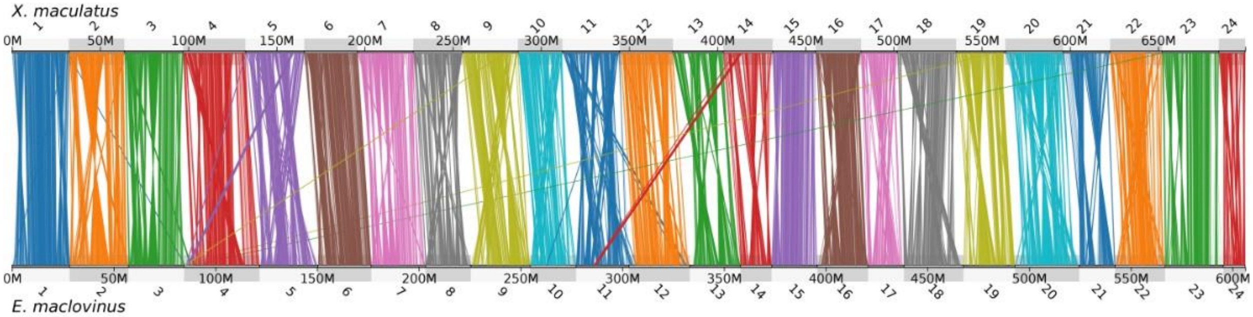


**Figure S2**. **Manually curated and annotated *cry3a* from *D. mawsoni* and *P. albipinna* genome assembly**. *E. maclovinus cry3a* gene structure is given as reference. "e" stands for exon. The two Thr residues encoded by the mini-e14 is indicated. Relative lengths of exons (rectangular boxes on grey bar) and between exons are to scale.

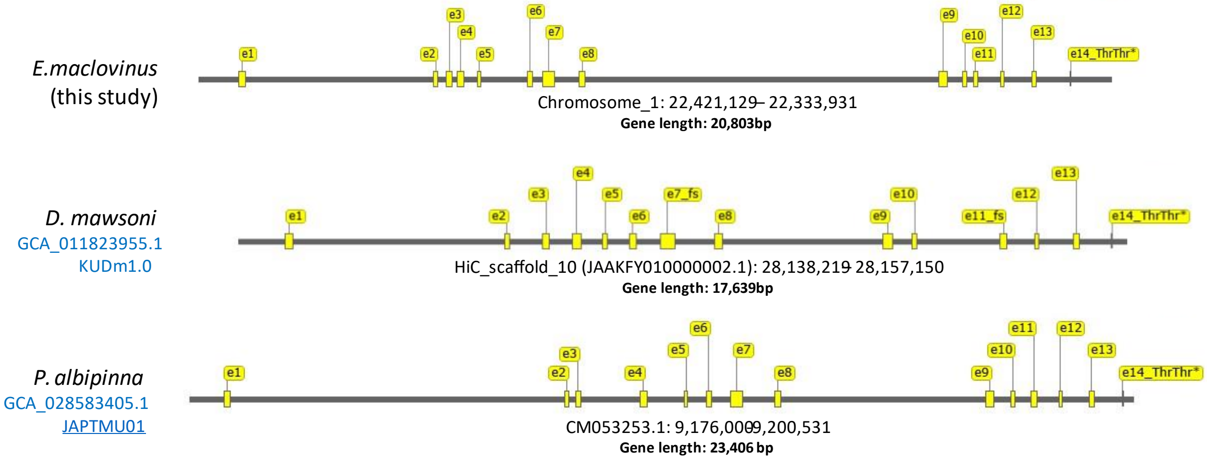

#### Supplemental File

#### File S1. Transcript sequences of circadian genes in *E. maclovinus* obtained from Iso-Seq and RNAseq transcriptomes.

```
>emac_fin_isoseq_HQ_transcript/28087_arntl1a mRNA_complete cds
GGAGAAACTGGGTCACCGATCGAAAATCGTTTATGTCACAGTTCTGTGGGTTGCTATTGC
ACATTACCAGCGCCTTTCAATGGATAGACCAATCTCATTTCATATGCTGGGTAGACGGGT
TGCACCAGCTCTTTGAGGATCAAATTAAATCGGGGCACAGCGACCACATCAATCCGGACA
AAACTAGGTCAGGCTGAATCAGAGCAGATCAGAGCAGACCCCGCTGATCCAGGCGATCCC
CGGCCCGTTCCAGTTTCATATCCAGTTGAGGGCCCAAGAGTGGATGGAAGGAGATGATTT
TAACACAGAGGCTGTGATGAACATCTGTGATGACCTGATGGCAGACCAAAGGATGGACAT
CTCGTCTACAATGACCGACTTCATGTCGCCCGGCTCCACCGACCTCATCTCCAGCTCCAT
CAGCACGCCCGGCATGGACTACACCCGCAAGAGGAAGGGCAGCACCACCGATTACCAAAT
TGATGGCTTCTCATTTGATGACATGGATCCAGACAAAGACAAACTGGGAGGTGACCAACA
GGGCCGGATTAAGAATGCCAGAGAGGCTCACAGTCAGATCGAGAAGCGGCGCAGGGACAA
GATGAACAGCTTCATCGACGAGCTGGCCTCTCTGGTGCCCACCTGCAACGCCATGTCCCG
CAAACTGGACAAGCTGACGGTGCTGCGCATGGCCGTCCAGCACATGAAGACACTAAGAGG
TGCAGCCAACCCGTACACAGAGGCCAACTACAAGCCCTCCTTCCTGTCGGACGAGGAGCT
GAAGCACTTGATATTGAGGGCGGCTGATGGCTTCCTGTTTGTGGTTGGGTGTGACCGCGG
GAAGATCCTCTTTGTCTCTGAGTCCGTCTTCAAGATTCTCAACTACAGCCAGAATGACCT
GATAGGTCAGAGTCTGTTTGACTACCTTCACCCCAAGGACATCGCCAAGGTCAAAGAGCA
GCTGTCCTCTTCAGACACAGCCCCTCGAGAGCGGCTCATCGATGCTAAAACCGGTCTCCC
AGTGAAGACGGACATCACCCCCGGCCCGTCCAGACTCTGCTCTGGAGCCAGACGCTCGTT
CTTCTGCAGGATGAAGTGCAACAGGCCTTCCGTCAAAGTGGAGGACAAAGACTTCCCCTC
CACCTGCTCCAAGAAGAAAGCCGATCGTAAGAGCTTCTGCACCATCCACAGCACGGGCTA
CCTGAAGAGCTGGCCCCCCACCAAGATGGGTCTGGACGACGACAACGAGCCCGACAACGA
GGGCTGCAACCTCAGCTGCCTGGTGGCCATCGGCCGGCTGCATCCCCACATTGTCCCCCA
GCCCAGCCTGGCAGACATCCGGGTGAAGCCCACCGAGTATGTCTCCAGACACGCCATCGA
CGGGAAGTTTGTCTTCGTCGACCAGAGAGCCACAGCCATCCTGGCCTATCTCCCCCAGGA
GCTGCTGGGAACCTCCTTCTATGAGTATTTCCACCAGGACGACATTGGACACCTGGCAGA
GTGTCACAGACAGGTGCTGCAGATGAGAGAGAAGATCAACACCAACTGCTACAAGTTCAA
GATCAAAGACGGCTCCTTCATCACGCTGAGGAGCCGCTGGTTCAGCTTCATGAACCCCTG
GACCAAGGAGGTGGAGTACATCGTCTCCACCAACACCGTCGTGTCGTGTCCTATGATGGA
GGGACCGGACTACCCTCAGTCTGCTGCCTCCCCTCAGAGCATGGACAGTGTTCTCACCTC
AGAGGGTGGAGGGAGGCGGGCGCTGCAGACGGTGCCTGGCATCCCCGGGGGGACAAGAGC
AGGAGCCGGGAAAATAGGACGCATGATTGCAGAGGAGGTGATGGAGATCCAGAGGATCCG
AGGCTCCTCCCCCTCCAGCTGTGGCTCCAGCCCTCTGAACATCACCAGCACCCCCCCACC
TGACACCTGCTCCCCAGGGGGCAAGAAGATTCAGAACGGTGGGACACCTGACCTGCCAAA
CACAGGGATCCTTCCTGGACCTGACTCGGTGGGCTACCCCTACTCCAACCAGTCCATCAT
GAGTGACAACTCCCACCTGAGTATAGACATCATGGACGAGCCGGGCTCCAGCAGCCCCAG
CAACGACGAGGCGGCCATGGCGGTCATCATGTCCCTGCTGGAGGCCGATGCCGGGCTGGG
GGGGCCCGTGGACTTCAGCGACCTGCCCTGGCCTTTGTGAGGAAGCTGGGTGGAGAGGAG
GAGGCGGGGACTCTGTCAGCAGAGCCGTGGTGAGAGAGTTCAGTCAGGGTGAAGTCTGAC
TGTTCCCCGTCCCCGACTCCCCCCCGCAGAGACATCATCTAACAGATATATCATCCAGTA
GAGAGGCTGCAGGACTACAGGGGGGTCCTTCAGTCCGGACAGATCACACCTACTCTTTTG
TTCATCATTCCCTGTAGGGGGGACTGTTATTATACAGAAATCCTCCCAAAGGTTAAGATG
ATTTCTACAGTGTTAGGGTCTGCACACTGTTTCAGCCGTGGGGGGGAGATCTCTAGGTAC
AGGACCCCGCTGCTAACATGAAGGACTGGACTCTGTCTCACTATGGCTCTGGGCCCCTTC
ACTGTCAGCCCCCTCCACACCAACACAGGACTCTGAGGCGTCGACTAACGCTGGCTGAAG
TTTGTTGTGGGGCGGGGGGGGGGGGGTTGATGCCGTACACACACGGCTAAGGTGCACCCC
CCCCCCCCCCCATCTGACAATAATGTATCAAATCTTTAAATATTATCATGTTAAGAGTAT
ACTTTATGCAAATATATATGAGTATGATGAGTGTATTGTATGTATCAGCAGTGAAATATT
TTTAAAATAAAATAATTTGTTTCTGTC
```

```
>emac_fin_isoseq_HQ_transcript/9158_arntl2 mRNA_complete cds
AGAATCGTTTCCCTCCGGTTCTCTACTCTGCCCTGATTGCACAGCATACAGCTACCTCCG
TGCGCCTTTTCCCTCTTTTTTCTTCATTTGGGGAATACAAGGCAGTTTTTCATCCTTGTG
CCACCGTCTCGGCTCCCCCACTCCCCGCCCTGTTATCACAGCAACACCCCGGAGCAGTCG
ACCGCCTCATTCATGCATTGGAGTCCCCCGCGCACCGGCTCCGGCTCCAGCCATGTCGGC
CAGGAATGCAGCGGCTGGCGGCGGTGACAGAGTGGGAGGCGAACCGGCAGATGTGCTGGT
TGAGGACAAACAGAGTGATTCTGTGTCATTGCCCAGCCCGATGACCCCGGCCTCAGCTGC
TGGCATGTCCTTGAGCATGGAGATGCCCCGGAAGCGTAAGGGCAGCGTGGACAACCAGGA
TACAAAATCTGCTTCAATGGATATGGACATTGAAGACTTGGAAGAGAGGCACAACGGGTC
CGATGGTGAGGACCAACATGTCAAAATGAAATGCTTCAGGGAACCACACAGCCAAATTGA
GAAGAGGAGACGGGACAAAATGAACAATCTCATTGACAAACTATCAGCCATGATCCCTAC
GTGTAACCCCATGTCCCGTAAGCTGGACAAACTCACTGTTCTCCGAATGGCCGTGCAGCA
```

```
CCTCAAATCTCTCAAAGGTTCAGCAAGTTCTTTTTCTGAAGCCAACTACAAGCCCTCATT
CCTTCCCAACGAGGATCTCAAACATCTTATCCTCAAGGCTGCGGACGGGTTCCTGTTTGT
AGTGGGCTGTGACCGTGGGAAAATTGTGTTTGTCTCAGAGTCCGTCACGAAGATATTAAA
TTATAGTCGGGCGGAGCTGATTGGACAGAGTCTGTTTGATTACATTCACCCAAAGGACAT
GGGAAAAGTGAAGGAGCAGCTGTCAGCCTCTGAATTATACCCTCGTGAACGGCTAATAGA
TGCTAAAACCGGTCTGCAGGTCCAGGCTGACCTCCCCGTCGGTGCAGCACGACTGTGTAC
AGGCGCACGGCGCTCATTCTTCTGTCGCATGAAGATAAACAAGATTTGTGTCAAAGTAGA
GGAGAAGGAATTCCAAGCCAGCACCTCCAAAAAGAAAGAGTCCCAGAAATACTGCACGGT
CCACTGTACAGGCTACATGCGCAGCTGGCCCACCAGTCAGTTGGGAGCTGAGGGAGAGGG
CGATGCAGACAAGCAGGACAGCTCACACTTCAGCTGCCTGGTGGCGGTGGGACGCGTCCA
CTCCCACTCCACTGCCCAGGTTAATGGAGAAGTCAGAGTAAAACCCACAGAGTTCATCAC
ACGCTACGCCATGGATGGCAAATTCACCTTTGTCGATCAAAGAGCGACAACCATTCTTGG
TTATCTTCCCCAAGAACTGCTAGGGACATCATGCTACGAGTACTTCCATCAAGACGACTT
ACCGCATTTAGCTGACAGACACCGAAAAGTGCTGCGGAGTAAAGAAAAAATTGAGACAAA
CTGCTATAAGTTCAAAACGAAAAGTGGCTCTTTCGTCACTCTGCAAAGTCAGTGGTTTAG
TTTTGTAAATCCTTGGACCAAAGAAGTGGAATACTTAGTGTCAACAAACACAGTTATATC
GTATGATAACAGTCGAACCTGTCGGTCAGGAAGCAAGTCTGAACAGTCAAGCAATTCCAA
GACCTCTGAAGATGGCAAGAAGTCCCTTCCAGTTATACCAGGCATCTCCACCACAACTGG
AGCTATGATTTATGCTGGAAGCATCGGGACCCAGATTGCCAATGAGCTGCTGGATTCCAA
CAGGATGAACTCTTCACCTTCCAGTGGCAGCGTCAGCCCTTTCAGTCTATCCCAGGACAA
ATGTCCCCACAATCAGATCAGCAACAATGTGCCAAACGGAGAGGCAACAGACATGGAGAT
TGCAGGAAAGTCCAGCTCAGAGGAGGATGCTCAAGGAGCTACATTCTCAGGAGCCGAGTC
TCTCATGGGGGAGAATCCCCAGATGGATTTGGACAGCGTGGTTGGACCGGGCCTTGGCAG
TCTCAGCAGTGACGAAGCAGCCATGGCAGTCATCATGAGCCTCCTGGAGACGGACACAAA
CCTGGGCGATGCCGTGGACTTTGAAGAGATGCACTGGTCCTTATAGAATCTGCTCTGAGA
GTGTGAGACACCGCAACACACTGAAAAACACTTGAACTCTCATTTTTGTGATGAGAACTG
CTTGAATTCTCCCTCTTAAGACTTGGCATTCCTTTTTGTTTATTTATTAGTTGAGTTTTG
CCACATCAGCTGTGTAGTCTTTGTGTGGCTTCTGTAGCAGGGAGCGGGATCGACCTCAAT
AAGGGACGCCTGTTGACCCTCGTCAGGATGAGGATGCGCCTCTAAAGCCTTACAGCGGTC
TGCCTGCCCTCACACGCTGTCCCTCACACCTACTGTCATCCTCTCACAAGGCTGAAGAAC
CCATGAACACTATTGAAAATGGCTTGGGATTTTCTCCTCCACCTACATTTTAACTAAGGA
TGGGCCTGACGATGCAAAGAAAACAAGATTGCACTGAGTTTTTCTTTTAAAGGTTGTCAT
TTAACTAGGTACTGTATGTTTAATGAAAATACAAGAACGACAACCTGTCCCCTCTCTGTG
ACCCCCTCAGGATCCTAGCCGTAGGTAGCCAGTGTAACATGGCAAACCACACTTCCATAA
CCTCAAACTTTGAGAGTCACCTAATGGTGAGCCACTCGGAGATTTTCTCTTGTCAGTATC
TGAACATTTTCCACGGAAACAGTCCAGCAACCTCCTCACTTGAACATGCTGTATACGTGT
GGATTTGGTATGAGATTTGCAAACATCATGTCTTCAAGCCATATCTGTCTTAAAGGGACA
GTTTGCATATGAAGAATATTTTTATTTTTTGGTATGTACACTTCTATGTACATCTAATAT
AGCCCTGGATTTTATTCCCTCAACTCTAAACATTTAACATGTATTGCCAGAAATTATCAG
TCTATTATTTGAGAGCATCCACTACTGATGGTTACCAAATCTTGAACTGATTTTTGAGCA
CTACAGAATCCCTTCAGGCTAAGTGAAAAGCCACACAAAGGAGTCTCTGTTGGCAACGGT
GAGCAGCAAACCGGTTTCAAACCTGTGACTTTTTATCAATACATGTTGGTTGACATTTGA
GTCTGACGGTAGAAGTTGAACCTCAGGATGTAATGGAAACAGAGCTGCCACGATTAAACA
TGGACTGCAGCAGTTGTCCTCCAATGTCTGAAATTCTATCCTACATACTCTTCTAGTAAT
TAGTCTGTAGTGGTTCACCTGTGTGTTTGTGTTCTGACCCTTATCATCAGCGTTATAATC
TCGCAGGTAAGCCGTAGCTTTAGCTGGTTTCTCCTGCACTTCCACTGTATGAGTCTTTCA
AAATACATGTTTATGTGCTTGTTAGAGGCACACAACTTTTTCATAGATACATTATGGCTT
CACGATATGAGGAGAATATAAGATATGTGATAAGATTGAATATCGCGCTAACTATATTAC
TTGAGTTAAATAAACAGATATTAAAGTCTGTTCTGCCTTTCTGCTGCTTTCAGTACATTG
CTAAAATACAACAGATTGCATGTTGACTTGACAGGAAATTATTCTTTTATTGAACAAATT
GAACTGAACACC

>emac_fin_Isoseq_HQ_transcript/7059_clockb mRNA_complete cds
GACCGACTCGATGCAACGTTCAAGTAAGCAAAACACGGAGGCTGCTGTCGATTACTCAGG
AAGGTGAGGTGTGGCTCACCCATGAAGAGTTGTCAGTGCCTGGAAGGATAGGATCTTAAT
TAGCTTTTCACATTCCAGAAAGTTTGGTATCGCCATGTCTTTCCTGTGGCCCGTTGTCCT
CTCGTGACCCCAATGACCTCAACAGCAGTTTTGACCATAGAGTCAGCTTTGACCCCCTAC
CCTTTCTCTGACCCCTGAGAAAAGACATTCAGTGACCACGTGCATAAGCTGTGTTGGGGT
GGTCTTCCAAATGCCTACCTCACGCCTAAAGAGCAAATCACTGCAGCCACAGGCCTGATC
AGCAGCTACCATGACATCAAGCATAGGGGATGATTGTAGCATCTTTGATGGGTTGATGGA
AGAAGATGAAAAGGACAAAGCAAAACGCGTGTCCCGTAACAAGTCTGAGAAGAAGCGGAG
AGACCAGTTCAATGTCCTCATCAAGGAACTCGGCACGATGTTGCCGGGCAACACCAGGAA
GATGGACAAGTCCACCATCCTGCAGAAAAGCATTGACTTCCTGTGTAAACACAAAGAGAT
TGCAGCCCAGTCGGAGTCGAGTGAGATCAGGCAGGACTGGAAGCCTCCATTTCTCAGCAA
TGAGGAGTTCACCCAGCTCATGCTGGAGGCTCTGGATGGATTCTTTATAGCAATAATGAC
TGATGGGAACATCCTCTACGTCTCTGAGAGTGTCACCTCACTACTTGAACATCTACCGAC
TGACTTGGTGGACCAGAACCTGTTAAACTTCCTGCCACTCGGGGAACACTCTGATGTGTA
CAAGGCGCTGTCCACACACCCCACCGACCACGACAACCTTAGCTCCGATTACCAAAAGAC
TAAGAACCACATGGAGTTCTGCTGCCATATGCTGCGAGGGGCCATTGACCCCAAAGAACC
```

GCCCGTCTACGAGTATGTCAAGTTCATTGGCAACTTCAAGTCCCTCAACAATGTTCCCAA
CGTGACGAGGAATGGTCTAGCAGGGGTGTTACAGCGCTCACTGCAGCCTGCGTTCGACGA
CCAGGTGTGCTTCGTAGCCACAGTTCGGCTGGCCAAACCTCAGTTCATCAAGGAAATGTG
TACAGTGGAAGAGCCTAATGAAGAATTCACCTCGAGGCACAGCCTAGAATGGAAGTTCCT
CTTCTTAGACCACAGAGCTCCACCAATCATAGGCTACCTGCCGTTCGAGGTTCTGGGAAC
ATCAGGGTACGACTATTACCATGTGGACGACCTGGAGACATTAGCCAAGTGTCACGAACA
CTTGATGCAGTACGGTAAAGGAAAGTCTTGCTACTATCGTTTCCTGACTAAAGGTCAACA
GTGGATCTGGCTTCAGACGCATTACTACATCACCTATCACCAGTGGAACTCTCGACCTGA
GTTTATCGTCTGTACGCACACAGTAGTCAGCTATGCAGAGGTCAGGGCCGAACAGCGCAG
AGAGCTGGGCATTGAAGAGTCCAACCCGGAAGGGACTGTGAATAAGAGTCAGGACTCGGA
GTCAGAGTCACAGCTGAATACCTCCAGCCTCAAGGAGGCTCTGGAGCGATTTGATCGCAG
CCGAACGCCCTCAACATCCTCACGAAGTTCTCGGAAGTCCTCGTCGCATGTCTCTGACAA
CACTTGCACCTCCTCTAAGCTACACATGGACACAGCCACACCTCCTCGGCAGTCCATGGC
CTCCACCATGGTGATGACGTCACAGCGGCGCTCATCAATCAGCAGCCAGTCGATGAGCTC
TCAGACCACGAGCCAGAGTGTAACTCCAGGGATGATCACTCAGCAGCAGCAGCAGCAGCA
GCAGCAGCAACAACAACAACAGCAACAACAGCAACAACAACAACAACAGCAACAGCCACA
ACAGCAACAGCCACAGCCACAACAGGCAAACGTACAGCCGGTGATGGAGTTCTCAGCGCA
GGTGAATGCCATGCAGCATCTGAAGGATCAACTGGAACAGAGGACAAGACTAATCCAAGC
CAACATCCAGCGGCAGCAGGACGAGCTGAGACACATCCAAGACCAGCTGCAGAGGGTGCA
GGGACAGGGCATACAGATGTTGTTGCAGCAGCAGGGAGGAGCGATGAACGTGCAGCTCCC
TCAGATGGGGTCGGTCCAACAAACAGCCAACTTGACCAACCAGGTCCAGCAAACAACAAT
TAACCCCGTCCATTCAGGAACACAGCAGCTCACCATCCAGCAGCAGGCCCCGCCCCCCCA
GCAGAACATACAACAGCAACAGCAGCAGACCAATGCACTCACACAGCCCCAGCGTCAGCC
CCAACAACCTCCGCAGGCTCAGCCCCAGACCCAGGGCTCCGTTTCTGCTCCACTCTACAA
CACCATGATGATTTCCCAGCCCGGGCAGCCCAACGTGCTGCAGATCAGCACCAGCTTGCC
GCAGACCAACACACAGCAAGGCACCGCGGTGGCAACCTTCACACAAGACCGACAGATCCG
CTTCCCAGCAGGGCAGCAGCTGGTGACCAAGCTGGTGACGGCGCCAATGGCCTGTGGAGC
AGTGATGGTGCCCACCTCCATGTTCATGGGGCAGGTGGTCACTGCCTACAACCCCTTTGG
TGGACAACAGCAGGGGGGGCAGACACAGACCCTGACCCTGCAACCAGCCCAGCCCCCCCA
AGGTCAGCCTGATGGCCAGAACCAGACCACTGTAGTGGCACAGGGCGGGCAGCAGGGGCA
GCAGCAACAACAGCAATTCCTACAGGGCACTCGTCTTCTCCACAGTAACCAGTCTACCCA
GCTGATTCTGCAGGCAGCTTTCCCACTCCAACAACAGGGCACTTTCACACAGGCAACTCA
TCAACAGCAACCACAACAACAGCAACAACAGCAACAACAACAGCAACAACAGCAGCAACA
ACAACAACAGCAACAACAGCAGCAACGGCAACAGCAGCAGCAGCAACAATCCCACCACCA
GAGGCACCAGCAGCAGCTGAAACCTCAGCCCCAGAAACCGCAGAAAGCCCCATCGTCGCA
CAGGACTGAGAGTGTCAGCAGCCAGCCGCAGTGAAGTCTGGAGGAGAAAACATTGGGATT
TAGAACTCGGAGGGGAGTTGCTGCGTATATTGGTTTTGTTCACCTGGCTGTCGCAAAGTC
CATTTTCCCCAAAAAATGTGTCGTCCCATCTTCCTCCATCTGAAAGAAATTAATCATAGG
AAAGAGGAGAGCCCGTGAGTCCTCAAGGAACTCACCATAGATTGTCACCATGGCAACAGG
ATTGATTCAAATCCACCAATTGTGAAGGAAGATCCAGTCTCCGTGGAAACCTGACAAAAA
TACAAAGAACACCTTATCGCCATGGTAACAACAGATTGTTTCGAAGGTGTGTCAAAGTGG
ACTGTGACCTTGTAGAGATTTACTGTATGATAAAAGAAAAACTTGTAAATATCATATATA
GCCTAACAGAAAAACAGAAATGTAGTATTTTATATGATTGCATGGAAAGAAAAACTGTGT
AAGCTTTTATTAGTCGCTTTTGAAAATTGAATTTTTACTTTGTTCAAGTGTTTTCTGGTC
AAAGATGGAGCAATGACATGCTCTGTCAATCTTCTCCCACCCCCTTCTCTTAACAAAAAA
ACACGGTTCGGAATAACACGGCTTTAAAGTTCTGCAGTTTGGTTGGGTTTGGGTTTTTTT
GTGGAAGTTGTGTTTTTAAACACTGTGTCTCTGAGACAAAAATGGCTCTTTTTAACTTTG
TAAGTACAAATTATCTTTGGC

>emac_fin_isoseq_HQ_transcript/7712_clockb mRNA_complete cds
GGCGCATGACTGTACGTCGTAATATGAAGTAGGCCTCCTGTTTTTCTCCCCAACCCGGAG
TCCTCCGAGCCCCCACCGACTCGATGCAACGTTCAAGTAAGCAAAACACGGAGGCTGCTG
TCGATTACTCAGGAAGGTTTGGTATCGCCATGTCTTTCCTGTGGCCCGTTGTCCTCTCGT
GACCCCAATGACCTCAACAGCAGTTTTGACCATAGAGTCAGCTTTGACCCCCTACCCTTT
CTCTGACCCCTGAGAAAAGACATTCAGTGACCACGTGCATAAGCTGTGTTGGGGTGGTCT
TCCAAATGCCTACCTCACGCCTAAAGAGCAAATCACTGCAGCCACAGGCCTGATCAGCAG
CTACCATGACATCAAGCATAGGGGATGATTGTAGCATCTTTGATGGGTTGATGGAAGAAG
ATGAAAAGGACAAAGCAAAACGCGTGTCCCGTAACAAGTCTGAGAAGAAGCGGAGAGACC
AGTTCAATGTCCTCATCAAGGAACTCGGCACGATGTTGCCGGGCAACACCAGGAAGATGG
ACAAGTCCACCATCCTGCAGAAAAGCATTGACTTCCTGTGTAAACACAAAGAGATTGCAG
CCCAGTCGGAGTCGAGTGAGATCAGGCAGGACTGGAAGCCTCCATTTCTCAGCAATGAGG
AGTTCACCCAGCTCATGCTGGAGGCTCTGGATGGATTCTTTATAGCAATAATGACTGATG
GGAACATCCTCTACGTCTCTGAGAGTGTCACCTCACTACTTGAACATCTACCGACTGACT
TGGTGGACCAGAACCTGTTAAACTTCCTGCCACTCGGGGAACACTCTGATGTGTACAAGG
CGCTGTCCACACACCCCACCGACCACGACAACCTTAGCTCCGATTACCAAAAGACTAAGA
ACCACATGGAGTTCTGCTGCCATATGCTGCGAGGGGCCATTGACCCCAAAGAACCGCCCG
TCTACGAGTATGTCAAGTTCATTGGCAACTTCAAGTCCCTCAACAATGTTCCCAACGTGA
CGAGGAATGGTCTAGCAGGGGTGTTACAGCGCTCACTGCAGCCTGCGTTCGACGACCAGG
TGTGCTTCGTAGCCACAGTTCGGCTGGCCAAACCTCAGTTCATCAAGGAAATGTGTACAG

TGGAAGAGCCTAATGAAGAATTCACCTCGAGGCACAGCCTAGAATGGAAGTTCCTCTTCT
TAGACCACAGAGCTCCACCAATCATAGGCTACCTGCCGTTCGAGGTTCTGGGAACATCAG
GGTACGACTATTACCATGTGGACGACCTGGAGACATTAGCCAAGTGTCACGAACACTTGA
TGCAGTACGGTAAAGGAAAGTCTTGCTACTATCGTTTCCTGACTAAAGGTCAACAGTGGA
TCTGGCTTCAGACGCATTACTACATCACCTATCACCAGTGGAACTCTCGACCTGAGTTTA
TCGTCTGTACGCACACAGTAGTCAGCTATGCAGAGGTCAGGGCCGAACAGCGCAGAGAGC
TGGGCATTGAAGAGTCCAACCCGGAAGGGACTGTGAATAAGAGTCAGGACTCGGAGTCAG
AGTCACAGCTGAATACCTCCAGCCTCAAGGAGGCTCTGGAGCGATTTGATCGCAGCCGAA
CGCCCTCAACATCCTCACGAAGTTCTCGGAAGTCCTCGTCGCATGTCTCTGACAACACTT
GCACCTCCTCTAAGCTACACATGGACACAGCCACACCTCCTCGGCAGTCCATGGCCTCCA
CCATGGTGATGACGTCACAGCGGCGCTCATCAATCAGCAGCCAGTCGATGAGCTCTCAGA
CCACGAGCCAGAGTGTAACTCCAGGGATGATCACTCAGCAGCAGCAGCAGCAGCAGCAGC
AGCAGCAACAACAACAACAACAACAACAACAACAACAACAACAGCAACAGCCACAACAGC
AACAGCCACAGCCACAACAGGCAAACGTACAGCCGGTGATGGAGTTCTCAGCGCAGGTGA
ATGCCATGCAGCATCTGAAGGATCAACTGGAACAGAGGACAAGACTAATCCAAGCCAACA
TCCAGCGGCAGCAGGACGAGCTGAGACACATCCAAGACCAGCTGCAGAGGGTGCAGGGAC
AGGGCATACAGATGTTGTTGCAGCAGCAGGGAGGAGCGATGAACGTGCAGCTCCCTCAGA
TGGGGTCGGTCCAACAAACAACCAACTTGACCAACCAGGTCCAGCAAACAACAATTAACC
CCGTCCATTCAGGAACACAGCAGCTCACCATCCAGCAGCAGGCCCCGCCCCCCCAGCAGA
ACATACAACAGCAACAGCAGCAGACCAACGCACTCACACAGCCCCAGCGTCAGCCCCAAC
AACCTCCGCAGGCTCAGCCCCAGACCCAGGGCTCCGTTTCTGCTCCACTCTACAACACCA
TGATGATTTCCCAGCCCGGGCAGCCCAACGTGCTGCAGATCAGCACCAGCTTGCCGCAGA
CCAACACACAGCAAGGCACCGCGGTGGCAACCTTCACACAAGACCGACAGATCCGCTTCC
CAGCAGGGCAGCAGCTGGTGACCAAGCTGGTGACGGCGCCAATGGCCTGTGGAGCAGTGA
TGGTGCCCACCTCCATGTTCATGGGGCAGGTGGTCACTGCCTACAACCCCTTTGGTGGAC
AACAGGGGGGGCAGACACAGACCCTGACCCTGCAACCAGCCCAGCCCCCCCAAGGTCAGC
CTGATGGCCAGAACCAGACCACTGTAGTGGCACAGGGCGGGCAGCAGGGGCAGCAGCAAC
AACAGCAATTCCTACAGGGCACTCGTCTTCTCCACAGTAACCAGTCTACCCAGCTGATTC
TGCAGGCAGCTTTCCCACTCCAACAACAGGGCACTTTCACACAGGCAACTCATCAACAGC
AACCACAACAACAGCAACAACAGCAACAACAACAGCAACAACAGCAGCAACAACAACAAC
AGCAACAACAGCAGCAACGGCAACAGCAGCAGCAGCAACAATCCCACCACCAGAGGCACC
AGCAGCAGCTGAAACCTCAGCCCCAGAAACCGCAGAAAGCCCCATCGTCGCACAGGACTG
AGAGTGTCAGCAGCCAGCCGCAGTGAAGTCTGGAGGAGAAAACATTGGGATTTAGAACTC
GGAGGGGAGTTGCTGCGTATATTGGTTTTGTTCACCTGGCTGTCGCAAAGTCCATTTTCC
CCAAAAAATGTGTCGTCCCATCTTCCTCCATCTGAAAGAAATTAATCATAGGAAAGAGGA
GAGCCCGTGAGTCCTCAAGGAACTCACCATAGATTGTCACCATGGCAACAGGATTGATTC
AAATCCACCAATTGTGAAGGAAGATCCAGTCTCCGTGGAAACCTGACAAAAATACAAAGA
ACACCTTATCGCCATGGTAACAACAGATTGTTTCGAAGGTGTGTCAAAGTGGACTGTGAC
CTTGTAGAGATTTACTGTATGATAAAAGAAAAACTTGTAAATATCATATATAGCCTAACA
GAAAAACAGAAATGTAGTATTTTATATGATTGCATGGAAAGAAAAACTGTGTAAGCTTTT
ATTAGTCGCTTTTGAAAATTGAATTTTTACTTTGTTCAAGTGTTTTCTGGTCAAAGATGG
AGCAATGACATGCTCTGTCAATCTTCTCCCACCCCCTTCTCTTAACAAAAAAACACGGTT
CGGAATAACACGGCTTTAAAGTTCTGCAGTTTGGTTGGGTTTGGGTTTTTTTTGTGGAAG
TTGTGTTTTTAAACACTGTGTCTCTGAGACAAAAATGGCTCTTTTTAACTTTGTAAGTAC
AAATTATCTTTGGC

>emac_fin_isoseq_HQ_transcript/24113_cry1a mRNA_complete cds
GAGTGTGGGCGGGGTCACGCTTGTAGGTGCGGCGAGAAGATTGAGCAGCAATTCGTGGAC
TTTGAGGAGAATTTACTCAAAACATCCGACTGACAGTGCAGGAGAAGACTCAAGCGGAAC
AGAGAAGGATTTATAAAAGCGACTTTTATGTTCAGGTTGTTTTAGGACGACTCTCTTGAG
AATATCGAAGGAACTGTGTTTACATAACCAGCTTCCCCTCTAACGCGGCGAGGAGCAGCC
TGTTTCCACGGTAACTTCTGACTAAGGAACAACTTCAATTTACAGGCTATTTAATTCCTC
TGCTACGTTATTGATATTCAATTTGAGTGCAGAAACGTTTTAGTACATGTATTGTTGTAC
AAGCTTTTTTCATCTTTCAGAAAAACAAGTTATACAACGCAAAACGAGGTTACTGCCTGT
CAGCGGACTGGGTCCTAAAGCCGATCGGGTTGCACATCGTGCCCAAATGGTTCAGCAGAC
TACTGTTCTTAAATTCTTTGACATTATGGAAAAAGTTCTGTTAGAACAAGGTTAATTGCA
AACTTGGACTGCGTCACTAAAAAGCAACGAGCCGTATGCTTCAATAGCATACGAGCTGAA
CAAGCCTACTTTTCTTTTGACATTTGGATTCGCTGTGAACTGATTTCCCCTTCTTGGATA
TACTGTTTTGTTTAACAGTGAGTCGGAGTAAGAAAGTAGACTTCTTAAATAACAAAAAAG
AAAAAGTAGGACTCGTCCACTTGAGTAATGGTCATTAATACGATCCACTGGTTCAGGAAG
GGCCTGCGGCTCCACGACAATCCGTCGCTCAAAGACTCCCTGCTGGGGGCGGATACTGTC
CGCTGTGTCTACATTCTCGACCCCTGGTTCGCAGGATCCTCCAACGTTGGGATCAACAGG
TGGAGGTTCTTACTGCAGAGTCTGGAGGACTTGGACTCCAGCCTCCGTAAGCTCAACTCT
CGACTGTTTGTGATCCGAGGCCAACCCACTGATGTCTTTCCCAGACTCTTCAAGGAATGG
AATATTTCTCGTTTGTCCTACGAGTACGACTCTGAGCCCTTTGGGAAAGAACGAGATGCA
GCGATTAAGAAACTGGCCTGTGAGGCTGGAGTGGAGGTGACCGTTCGCATCTCCCACACA
CTCTATGACCTGGACAAGATCATAGAGTTAAATGGGGGCCAGTCACCTCTGACCTACAAG
CGGTTCCAGACCCTCATCAGCCGGATGGATGCGGTGGAGGAGCCTGCAGACTCCATCACG
GCGGACATCATGGGGAAGTGCAGGACGCCGCTGTCCGAAGACCATGATGACAAGTTTGGC

GTCCCCTCATTGGAGGAGCTGGGTTTTGATACTGAAGGTCTTTCCTCCGCTGTGTGGCCG
GGGGGAGAGACGGAAGCCCTCACACGACTTGAGAGGCATCTGGAGAGGAAGGCGTGGGTG
GCCAACTTCGAGCGTCCCAGAATGAACGCCAACTCGCTGCTCGCCAGCCCGACCGGCCTC
AGCCCATACCTGCGCTTTGGCTGCCTCTCCTGCCGCCTCTTCTACTTCAAACTCACCGAC
CTCTACAAGAAGGTGAAGAAGAACAGCTCCCCTCCTCTCTCGCTGTATGGTCAGCTGCTC
TGGCGCGAGTTCTTCTACACAGCAGCCACCAACAACCCTTGCTTCGACACGATGGAGAGC
AACCCCATCTGTGTCCAGATCCCCTGGGACCGAAACCCAGAGGCGCTGGCCAAGTGGGCG
GAGGGCCGCACCGGCTTCCCCTGGATCGACGCCATCATGACGCAGCTGAGACAGGAGGGC
TGGATCCACCACCTCGCCCGACATGCTGTCGCCTGCTTCCTGACCCGGGGAGACCTGTGG
ATCAGCTGGGAGGAGGGCATGAAGGTGTTTGAGGAGCTGCTGCTGGATGCAGACTGGAGT
GTGAACGCGGGCAGCTGGATGTGGCTCTCCTGCAGCTCTTTCTTCCAGCAGTTCTTCCAC
TGCTACTGCCCCGTGGGCTTCGGCCGCCGCACAGACCCCAACGGAGATTACATACGGCGC
TACCTGCCCATTCTCAGAGGCTTTCCAGCCAAGTACATTTATGATCCGTGGAATGCTCCA
GAGAGTGTGCAGAAGGCAGCAAAGTGTATCATTGGAGTGCATTACCCCAAACCCATGGTG
AACCACGCAGAGTCCAGCCGGCTCAACATCGAGCGGATGAAGCAGATCTACCAGCAGCTG
TCCTGCTACAGAGGCCTCGGCCTTCTGGGTTCAGTTCCCTCCACTTCTAACGGTAACGGA
GAGACGTCCTCTGACGGGATGGGATTCTCTGCTGAGGCTACACACAACACTGCTGCAGCT
CCATCTGGCTATCAGATGAGCGTTCACTCGCAGGGAGATTGGCAGAGCGGCGTCATGACG
TTCATGCAGGGCGACACGCAAACCAGCTGCAGCTCACAGCAGCAGGGTTATGCAGGCACC
AGTAGTAGTATGCTGTGTTACACCCAAGGTGCACAGCAAATTCCAAAAGGACCTGAACAC
CACGTCACCCCTCTGTCCGGTGGGAAAAGACACTGCGAGGACCCTGGGAATGGCAGAGGC
TCTAAAATCCAGAGACAGAGTACACACTAAGCGAGTGTGTGCAGTCAGACCAAGGACACA
GCCAACACAAAATGGACATTTTTGTCTCAGCACTTCAAGATGAGCAACACAAACAAGACT
GAATATTATAATGTGGACTGCACGGCACTGCGACGTGCAGACGACTCAGAAGCAACTCTG
TTCTTTTCTTTGGGTTAACATAGCCATGACAACGGTCTGTGTGTATATAATGAATTTGCT
CATTTGACATACAACTTGTACATATTTGAATATTGTATGTGTGTTTTCTCTCACTGTGTA
GCCATAGACCCATTGATATATTTTTGATATATGAAATCATTGTGGATGGATCTTTGTTTT
TTTTTAATATTGATTAAAAACAACCTTTGTAAAATGTC

>emac_fin_isoseq_HQ_transcript/13334_cry1a mRNA_complete cds
AGGTGCGGCGAGAAGATTGAGCAGCAATTCGTGGACTTTGAGGAGAATTTACTCAAAACA
TCCGACTGACAGTGCAGGAGAAGACTCAAGCGGAACAGAGAAGGATTTATAAAAGCGACT
TTTATGTTCAGGTTGTTTTAGGACGACTCTCTTGAGAATATCGAAGGAACTGTGTTTACA
TAACCAGCTTCCCCTCTAACGCGGCGAGGAGCAGCCTGTTTCCACGGTAACTTCTGACTA
AGGAACAACTTCAATTTACAGGCTATTTAATTCCTCTGCTACGTTATTGATATTCAATTC
GAGTGCAGAAACGTTTTAGTACATGTATTGTTGTACAAGCTTTTTTCATCTTTCAGAAAA
ACAAGTTATACAACGCAAAACGAGGTTACTGCCTGTCAGCGGACTGGGTCCTAAAGCCGA
TCGGGTTGCACATCGTGCCCAAATGGTTCAGCAGACTACTGTTCTTAAATTCTTTGACAT
TATGGAAAAAGTTCTGTTAGAACAAGGTTAATTGCAAACTTGGACTGCGTCACTAAAAAG
CAACGAGCCGTATGCTTCAATAGCATACGAGCTGAACAAGCCTACTTTTCTTTTGACATT
TGGATTCGCTGTGAACTGATTTCCCCTTCTTGGATATACTGTTTTGTTTAACAGTGAGTC
GGAGTAAGAAAGTAGACTTCTTAAATAACAAAAAAGAAAAAGTAGGACTCGTCCACTTGA
GTAATGGTCATTAATACGATCCACTGGTTCAGGAAGGGCCTGCGGCTCCACGACAATCCG
TCGCTCAAAGACTCCCTGCTGGGGGCGGATACTGTCCGCTGTGTCTACATTCTCGACCCC
TGGTTCGCAGGATCCTCCAACGTTGGGATCAACAGGTGGAGGTTCTTACTGCAGAGTCTG
GAGGACTTGGACTCCAGCCTCCGTAAGCTCAACTCTCGACTGTTTGTGATCCGAGGCCAA
CCCACTGATGTCTTTCCCAGACTCTTCAAGGAATGGAATATTTCTCGTTTGTCCTACGAG
TACGACTCTGAGCCCTTTGGGAAAGAACGAGATGCAGCGATTAAGAAACTGGCCTGTGAG
GCTGGAGTGGAGGTGACCGTTCGCATCTCCCACACACTCTATGACCTGGACAAGATCATA
GAGTTAAATGGGGGCCAGTCACCTCTGACCTACAAGCGGTTCCAGACCCTCATCAGCCGG
ATGGATGCGGTGGAGGAGCCTGCAGACTCCATCACGGCGGACATCATGGGGAAGTGCAGG
ACGCCGCTGTCCGAAGACCATGATGACAAGTTTGGCGTCCCCTCATTGGAGGAGCTGGGT
TTTGATACTGAAGGTCTTTCCTCCGCTGTGTGGCCGGGGGGAGAGACGGAAGCCCTCACA
CGACTTGAGAGGCATCTGGAGAGGAAGGCGTGGGTGGCCAACTTCGAGCGTCCCAGAATG
AACGCCAACTCGCTGCTCGCCAGCCCGACCGGCCTCAGCCCATACCTGCGCTTTGGCTGC
CTCTCCTGCCGCCTCTTCTACTTCAAACTCACCGACCTCTACAAGAAGGTGAAGAAGAAC
AGCTCCCCTCCTCTCTCGCTGTATGGTCAGCTGCTCTGGCGCGAGTTCTTCTACACGGCA
GCCACCAACAACCCTTGCTTCGACACGATGGAGAGCAACCCCATCTGTGTCCAGATCCCC
TGGGACCGAAACCCAGAGGCGCTGGCCAAGTGGGCGGAGGGCCGCACCGGCTTCCCCTGG
ATCGACGCCATCATGACGCAGCTGAGACAGGAGGGCTGGATCCACCACCTCGCCCGACAT
GCTGTCGCCTGCTTCCTGACCCGGGGAGACCTGTGGATCAGCTGGGAGGAGGGCATGAAG
GTGTTTGAGGAGCTGCTGCTGGATGCAGACTGGAGTGTGAACGCGGGCAGCTGGATGTGG
CTCTCCTGCAGCTCTTTCTTCCAGCAGTTCTTCCACTGCTACTGCCCCGTGGGCTTCGGC
CGCCGCACAGACCCCAACGGAGATTACATACGGCGCTACCTGCCCATTCTCAGAGGCTTT
CCAGCCAAGTACATTTATGATCCGTGGAATGCTCCAGAGAGTGTGCAGAAGGCAGCAAAG
TGTATCATTGGAGTGCATTACCCCAAACCCATGGTGAACCACGCAGAGTCCAGCCGGCTC
AACATCGAGCGGATGAAGCAGATCTACCAGCAGCTGTCCTGCTACAGAGGCCTCGGCCTT
CTGGGTTCAGTTCCCTCCACTTCTAACGGTAACGGAGAGACGTCCTCTGACGGGATGGGA
TTCTCTGCTGAGGCTACACACAACACTGCTGCAGCTCCATCTGGCTATCAGATGAGCGTT

```
CACTCGCAGGGAGATTGGCAGAGCGGCGTCATGACGTTCATGCAGGGCGACACGCAAACC
AGCTGCAGCTCACAGCAGCAGGGTTATGCAGGCACCAGTAGTAGTATGCTGTGTTACACC
CAAGGTGCACAGCAAATTCCAAAAGGACCTGAACACCACGTCACCCCTCTGTCCGGTGGG
AAAAGACACTGCGAGGACCCTGGGAATGGCAGAGGCTCTAAAATCCAGAGACAGAGTACA
CACTAAGCGAGTGTGTGCAGTCAGACCAAGGACACAGCCAACACAAAATGGACATTTTTG
TCTCAGCACTTCAAGATGAGCAACACAAACAAGACTGAATATTATAATGTGGACTGCACG
GCACTGCGACGTGCAGACGACTCAGAAGCAACTCTGTTCTTTTCTTTGGGTTAACATAGC
CATGACAACGGTCTGTGTGTATATAATGAATTTGCTCATTTGACATACAACTTGTACATA
TTTGAATATTGTATGTGTGTTTTCTCTCACTGTGTAGCCATAGACCCATTGATATATTTT
TGATATATGAAATCATTGTGGATGGATCTTTGTTTTTTTTTAATATTGATTAAAAACAAC
CTTTGTAAAATGTCTATTCAAGAACAATTCAATTTGATAAATCTTGTATCTATCTTAATA
TATTTTTCTGTTCTCTTATTGTACTTTCTATTATACTCGAGATGTATATCAACAAGTGTT
TCAATCATTCCCTTTCTCTTTCTGCTTCTGACTTTAATCCTCACCACTCTAGAGGCAATA
TCAGTTTAACATCAAGTTGTTGCCAGATCTGCTGTTACTGAAAGAAGCCGACAGAAATAC
ATGTTCTTCATATCATGGTGCAGCCCCTGCCTCCTTGTATAAGCTTCAGTAAATGTAGTA
CATAAGCTTCTTTGGGCACCAGGAAGAGCGCTATATAAATACCATGTATTAGTATTATGA
CATGAGAGAACTGCTTTACTTGCGATTTGTTCCTTCTCAATGTGCTTCTTTAATAAACTG
GGTTTAATAT

>emac_fin_isoseq_HQ_transcript/18412_cry1a mRNA_complete cds
GCTTGTAGGTGCGGCGAGAAGATTGAGCAGCCTGTTTCCACGGTAACTTCTGACTAAGGA
ACAACTTCAATTTACAGGCTATTTAATTCCTCTGCTACGTTATTGATATTCAATTCGAGT
GCAGAAACGTTTTAGTACATGTATTGTTGTACAAGCTTTTTTCATCTTTCAGAAAAACAA
GTTATACAACGCAAAACGAGGTTACTGCCTGTCAGCGGACTGGGTCCTAAAGCCGATCGG
GTTGCACATCGTGCCCAAATGGTTCAGCAGACTACTGTTCTTAAATTCTTTGACATTATG
GAAAAAGTTCTGTTAGAACAAGGTTAATTGCAAACTTGGACTGCGTCACTAAAAAGCAAC
GAGCCGTATGCTTCAATAGCATACGAGCTGCACAAGCCTACTTTTCTTTTGACATTTGGA
TTCGCTGTGAACTGATTTCCCCTTCTTGGATATACTGTTTTGTTTAACAGTGAGTCGGAG
TAAGAAAGTACACTTCTTAAATAACAAAAAAGAAAAAGTAGGACTCGTCCACTTGAGTAA
TGGTCATTAATACGATCCACTGGTTCAGGAAGGGCCTGCGGCTCCACGACAATCCGTCGC
TCAAAGACTCCCTGCTGGGGGCGGATACTGTCCGCTGTGTCTACATTCTCGACCCCTGGT
TCGCAGGATCCTCCAACGTTGGGATCAACAGGTGGAGGTTCTTACTGCAGAGTCTGGAGG
ACTTGGACTCCAGCCTCCGTAAGCTCAACTCTCGACTGTTTGTGATCCGAGGCCAACCCA
CTGATGTCTTTCCCAGACTCTTCAAGGAATGGAATATTTCTCGTTTGTCCTACGAGTACG
ACTCTGAGCCCTTTGGGAAAGAACGAGATGCAGCGATTAAGAAACTGGCCTGTGAGGCTG
GAGTGGAGGTGACCGTTCGCATCTCCCACACACTCTATGACCTGGACAAGATCATAGAGT
TAAATGGGGGCCAGTCACCTCTGACCTACAAGCGGTTCCAGACCCTCATCAGCCGGATGG
ATGCGGTGGAGGAGCCTGCAGACTCCATCACGGCGGACATCATGGGGAAGTGCAGGACGC
CGCTGTCCGAAGACCATGATGACAAGTTTGGCGTCCCCTCATTGGAGGAGCTGGGTTTTG
ATACTGAAGGTCTTTCCTCCGCTGTGTGGCCGGGGGGAGAGACGGAAGCCCTCACACGAC
TTGAGAGGCATCTGGAGAGGAAGGCGTGGGTGGCCAACTTCGAGCGTCCCAGAATGAACG
CCAACTCGCTGCTCGCCAGCCCGACCGGCCTCAGCCCATACCTGCGCTTTGGCTGCCTCT
CCTGCCGCCTCTTCTACTTCAAACTCACCGACCTCTACAAGAAGGTGAAGAAGAACAGCT
CCCCTCCTCTCTCGCTGTATGGTCAGCTGCTCTGGCGCGAGTTCTTCTACACGGCAGCCA
CCAACAACCCTTGCTTCGACACGATGGAGAGCAACCCCATCTGTGTCCAGATCCCCTGGG
ACCGAAACCCAGAGGCGCTGGCCAAGTGGGCGGAGGGCCGCACCGGCTTCCCCTGGATCG
ACGCCATCATGACGCAGCTGAGACAGGAGGGCTGGATCCACCACCTCGCCCGACATGCTG
TCGCCTGCTTCCTGACCCGGGGAGACCTGTGGATCAGCTGGGAGGAGGGCATGAAGGTGT
TTGAGGAGCTGCTGCTGGATGCAGACTGGAGTGTGAACGCGGGCAGCTGGATGTGGCTCT
CCTGCAGCTCTTTCTTCCAGCAGTTCTTCCACTGCTACTGCCCCGTGGGCTTCGGCCGCC
GCACAGACCCCAACGGAGATTACATAAGGCGCTACCTGCCCATTCTCAGAGGCTTTCCAG
CCAAGTACATTTATGATCCGTGGAATGCTCCAGAGAGTGTGCAGAAGGCAGCAAAGTGTA
TCATTGGAGTGCATTACCCCAAACCCATGGTGAACCACGCAGAGTCCAGCCGGCTCAACA
TCGAGCGGATGAAGCAGATCTACCAGCAGCTGTCCTGCTACAGAGGCCTCGGCCTTCTGG
GTTCAGTTCCCTCCACTTCTAACGGTAACGGAGAGACGTCCTCTGACGGGATGGGATTCT
CTGCTGAGGCTACACACAACACTGCTGCAGCTCCATCTGGCTATCAGATGAGCGTTCACT
CGCAGGGAGATTGGCAGAGCGGCGTCATGACGTTCATGCAGGGCGACACGCAAACCAGCT
GCAGCTCACAGCAGCAGGGTTATGCAGGCACCAGTAGTAGTATGCTGTGTTACACCCAAG
GTGCACAGCAAATTCCAAAAGGACCTGAACACCACGTCACCCCTCTGTCCGGTGGGAAAA
GACACTGCGAGGACCCTGGGAATGGCAGAGGCTCTAAAATCCAGAGACAGAGTACACACT
AAGCGAGTGTGTGCAGTCAGACCAAGGACACAGCCAACACAAAATGGACATTTTTGTCTC
AGCACTTCAAGATGAGCAACACAAACAAGACTGAATATTATAATGTGGACTGCACGGCAC
TGCGACGTGCAGACGACTCAGAAGCAACTCTGTTCTTTTCTTTGGGTTAACATAGCCATG
ACAACGGTCTGTGTGTATATAATGAATTTGCTCATTTGACATACAACTTGTACATATTTG
AATATTGTATATGTTTTCTCTCACTGTGTAGCCATAGACCCATTGATATATTTTTGATAT
ATGAAATCATTGTGGATGGATCTTTGTTTTTTTTTAATATTGATTAAAAACAACCTTTGT
AAAATGTCTATTCAAGAACAATTCAATTTGATAAATCTTGTATCTATCTTAATATATTTT
TCTGTTCTCTTATTGTACTTTCTATTATACTCGAGATGTATATCAACAAGTGTTTCAATC
ATTCCCTTTCTCTTTCTGCTTCTGACTTTAATCCTCACCACTCTAGAGGCAATATCAGTT
```

TAACATCAAGTTGTTGCCAGATCTGCTGTTACTGAAAGAAGCCGACAGAAATACATGTTC
TTCATATCATGGTGCAGCCCCTGCCTCCTTGTATAAGCTTCAGTAAATGTAGTACATAAG
CTTCTTTGGGCACCAGGAAGAGCGCTATATAAATACCATGTATTAGTATTATGACATGAG
AGAACTGCTTTACTTGCGATTTGTTCCTTCTCAATGTGCTTCTTTAATAAACTGGGTTTA
ATATAC

>emac_fin_isoseq_HQ_transcript/13348_cry1b mRNA_complete cds
GGATACAGAAACTTGAAGGGCTTGCATGTGAGACGGATTTGCCAAGCGTAACAATTGATA
TATCATTTTCACGTGTAAGTGAGTAAGGGTGTGCGGTAACAAACCGGAATGATTTGCGTG
TCTGTGCTGTCTTTGAGCTAGTATTGTGTACATTATCCTCCCCTCGTTATAAACGTCGCT
GCAGACTGATTTTTTTTTCTTCTTCTTGGAACGCACTTTAGTCAAATGTCTTCACTTGAT
CGTTTCTATTAATCGAGACAAGCTTGTTTTGTTTTTCTCACGTCTGCCGTTGTACCTGCT
GTTCCACATGCAGCCTTCTGTCAGGAGACTCAACTCAGTGGATACATTCATGGAGCAAGG
TGGGACAAACCATTAACTCAGACTCTCTGAAGCTACTGGCCGAAGCGTATTTCACCCTCG
GATGTAAATTTACACTTTCGCTAAGAAAACAACGTAACAGGGTGTCACTTGAAAACACAT
CCCATCTTTCCAACCCCGCCCGGAAAAATAAGAGAAGGCAGGCAACATGGTGGTCAACAC
CATCCACTGGTTCAGGAAGGGGCTGCGGCTACACGACAACCCGTCTCTAAGGGACTCTAT
CCGGGATGCGGACACGCTGCGCTGTGTTTACATCCTGGACCCCTGGTTCGCAGGGTCCTC
CAATGTGGGCATCAACAGGTGGAGGTTTTTGTTGCAGTGCTTAGAGGACTTGGATGCAGG
CCTGCGAAAAATCAACTCCCGCTTGTTTGTCATCAGAGGCCAGCCCACAGATGTCTTCCC
AAGGATTTTTAAGGAATGGCAGATTAACCGTTTATCTTATGAATATGACTCGGAGCCGTT
TGGCAAGGAGCGTGATGCTGCCATCCAAAAACTAGCCAGTGAGGCTGGGGTAGAGGTCAT
GGTGCGGACCGCTCACACCTTGTACAATCTGGACAAGATCATAGAACTGAATGAGGGTCA
ATCTCCCCTCACCTACAAGCGCTTCCAGGCCCTCATCAACCGTATGGATGCTGTGGAGCT
GCCGGCAGAGACCATCACGTCAGAGGTTATTAAAGAATGTGCCACACCCATCAGCGAAGA
GCATAATGAGAAGTTTGGGGTGCCCTCCCTAGAAGAGCTTGGTTTTGAAACAGAGGGCTT
GACCACAGCAGTGTGGCCAGGTGGAGAAACAGAAGCCCTCATGAGACTGGAGCGCCACCT
GGAGAGGAAGGCATGGGTGGCAAACTTTGAGCGTCCGCGTATGAACGCCAACTCCCTGCT
GGCCAGTCCCACAGGACTAAGCCCCTACCTGCGCTTCGGATGTCTCTCCTGTCGACTCTT
CTACTTCAGACTTACCGACCTGTACAAGAAGGTCAAGAAGAACAGCACACCGCCACTTTC
CTTGTACGGCCAGCTGTTGTGGCGGGAGTTCTTTTATACATCTGCCACGAACAACCCCTG
CTTTGATAAGATGGAGGGAAACCCTGTGTGTGTCCAGATCCCCTGGGACCGAAACCCAGA
GGCGCTGGCTAAGTGGGCAGAGGGGCAGACAGGCTTCCCCTGGATAGACGCCATCATGAC
ACAGCTGAGACAGGAAGGCTGGATCCATCATTTGGCGAGGCACGCTGTAGCCTGCTTCCT
CTCCAGGGGAGACCTCTGGGTCAGCTGGGAGGAAGGCATGAAGGTGTTTGAAGAGTTGCT
TATAGATGCCGACTGGAGTGTGAATGCTGGCAGCTGGATGTGGCTCTCTTGCAGTTCATT
TTTCCAGCAGTTTTTCCACTGCTACTGCCCTGTCGGATTTGGACGACGCACAGACCCCAA
TGGAGACTACATTAGGCGTTATTTGCCCATCTTGAAGGGCTTTCCAGCCAAATACATTTA
TGATCCCTGGAATGCCCCAGAGGACGTGCAGAAGGCTGCAAAGTGTATCGTAGGAGTCCA
CTACCCCAAGCCCATGGTGAACCATGCTGAGGCCAGCCGCATCAACATAGAGCGGATGAA
GCAAATTTACCAGCAGCTTTCAAGCTACAAAGGATTGGGCCTGCTGGCAACCATACCTTC
CAATCCCCACAGTGGTGTCGGAGGAGTCAACACAGGAAGCAGTCACGGTGTAGGAGGCTT
CTCAACATACACAGTATCCCCTCAAACAGAAAGAGTACAGACCACAGAAAAGAGGCGCCA
TGAAGAGGGTCCACATAGAGGCTCCAAATCGTTAAAGCAGAGCAAATAGTGTGCAAAAGG
TTGAATACAAAGGGAGCCCCAGTCATTCTGGATCTTTTGGATATCATCTTTATATGCATT
GTTATCAGCTGACACAGCAGCAGCTGTAACTAACAACCATAGATGACCATCATGATCTTC
AAATAAGACATGTTATTAATCTAAGGAAAGGCGGAAGGCATTTAGACTATAATTGAAGAT
AAAGAACCCTGAAACAGCGGGATGTAAAATGCAAAACCATTTGATCTAAAGTATCCACAT
CCAGCTTCTGCTACAGTATGTTAAATGTTGATCGTTATCATCAATGCCCCGCCCCCCCCA
TGACTGATGTAGGGCGTTGAGCGGGAAATTGGGTCATGAGGTCTTCTGTATGAACTTTGT
AACTTGTGTGAATGTGATCCATGCCTAAGTTCATCAAGACAATGAGCTCAACTTTTAGCG
GCTTTGGCCTTTACTTTTTCTTTAAAAAAATGTCTAAATCTGATTCCCATTCAAATGATG
GGAACCTATAACACAATTTGGAGTTGCTTATCTGACCTGGGTAACATACAGTGGTGTCTT
TCCTAAATATTGGGAAATATTAGGAAAAGAATGAATAAATATGTCATAAATGTACATTTG
ACAATGTACAAGCTCTGGACACTAAAATGAGATGTACAGTAATGAAAATGACAACCTCTC
TGTCAGGTTTGACAATAAATAATGCCTTTTATAAATGCCAACAGACAGTTTTTAATATTT
ATTATTTTTATTTCCAACTTTATATATTTGGTTGTGTGGTTTTACAACAACTTTTGTAAA
TAATGTGTTACGAAAACAAAAATCCACTGGATATGCATTTCAATTTCTCCCATGTGATCG
TAAATATGTTATTTTTGTATAAAGAATGAAAAGATTCTAACTGTATTGGAATGGCAATAA
ATGACACATTGTCATATAATATTGATTCATCCTTTTATTTGGACTATAATTGTAAAGTTG
TTTTAACTGACACCATTCTTTCAACAAGAATTAAGGTATTGTTGTGCCTTTTTAAAGTCT
GATAATCCTGCATAAGGTTTGTGATGAACAAGTTTGTACTCATTGTCCTTTATGCAGTTA
GAAAAGATTTGTCACGCTGCACATCTATATAATTGTGTCTTCAGAGAAAAGCTTTAATGG
GGGAAACATCTACCCTTATTTTAATTAAAGGAATTCCTCTAAATCCCATT

>emac_fin_isoseq_HQ_transcript/18671_cry1b mRNA_complete cds
GATACAGAAACTTGAAGGGCTTGCATGTGAGACGGATTTGCCAAGCGTAACAATTGATAT
ATCATTTTCACGTGTAAGTGAGTAAGGGTGTGCGGTAACAAACCGGAATGATTTGCGTGT
CTGTGCTGTCTTTGAGCTAGTATTGTGTACATTATCCTCCCCTCGTTATAAACGTCGCTG

```
CAGACTGATTTTTTTTTCTTCTTCTTGGAACGCACTTTAGTCAAATGTCTTCACTTGATC
GTTTCTATTAATCGAGACAAGCTTGTTTTGTTTTTCTCACGTCTGCCGTTGTACCTGCTG
TTCCACATGCAGCCTTCTGTCAGGAGACTCAACTCAGTGGATACATTCATGGAGCAAGGT
GGGACAAACCATTAACTCAGACTCTCTGAAGCTACTGGCCGAAGCGTATTTCACCCTCGG
ATGTAAATTTACACTTTCGCTAAGAAAACAACGTAACAGGGTGTCACTTGTTTGTAATTG
GAAAACACATCCCATCTTTCCAACCCCGCCCGGAAAAATAAGAGAAGGCAGGCAACATGG
TGGTCAACACCATCCACTGGTTCAGGAAGGGGCTGCGGCTACACGACAACCCGTCTCTAA
GGGACTCTATCCGGGATGCGGACACGCTGCGCTGTGTTTACATCCTGGACCCCTGGTTCG
CAGGGTCCTCCAATGTGGGCATCAACAGGTGGAGGTTTTTGTTGCAGTGCTTAGAGGACT
TGGATGCAGGCCTGCGAAAAATCAACTCCCGCTTGTTTGTCATCAGAGGCCAGCCCACAG
ATGTCTTCCCAAGGATTTTTAAGGAATGGCAGATTAACCGTTTATCTTATGAATATGACT
CGGAGCCGTTTGGCAAGGAGCGTGATGCTGCCATCCAAAAACTAGCCAGTGAGGCTGGGG
TAGAGGTCATGGTGCGGACCGCTCACACCTTGTACAATCTGGACAAGATCATAGAACTGA
ATGAGGGTCAATCTCCCCTCACCTACAAGCGCTTCCAGGCCCTCATCAACCGTATGGATG
CTGTGGAGCTGCCGGCAGAGACCATCACGTCAGAGGTTATTAAAGAATGTGCCACACCCA
TCAGCGAAGAGCATAATGAGAAGTTTGGGGTGCCCTCCCTAGAAGAGCTTGGTTTTGAAA
CAGAGAGCTTGACCACAGCAGTGTGGCCAGGTGGAGAAACAGAAGCCCTCATGAGACTGG
AGCGCCACCTGGAGAGGAAGGCATGGGTGGCAAACTTTGAGCGTCCGCGTATGAACGCCA
ACTCCCTGCTGGCCAGTCCCACAGGACTAAGCCCCTACCTGCGCTTCGGATGTCTCTCCT
GTCGACTCTTCTACTTCAGACTTACCGACCTGTACAAGAAGGTCAAGAAGAACAGCACAC
CGCCACTTTCCTTGTACGGCCAGCTGTTGTGGCGGGAGTTCTTTTATACATCTGCCACGA
ACAACCCCTGCTTTGATAAGATGGAGGGAAACCCTGTGTGTGTCCAGATCCCCTGGGACC
GAAACCCAGAGGCGCTGGCTAAGTGGGCAGAGGGGCAGACAGGCTTCCCCTGGATAGACG
CCATCATGACACAGCTGAGACAGGAAGGCTGGATCCATCATTTGGCGAGGCACGCTGTAG
CCTGCTTCCTCTCCAGGGGAGACCTCTGGGTCAGCTGGGAGGAAGGCATGAAGGTGTTTG
AAGAGTTGCTTATAGATGCCGACTGGAGTGTGAATGCTGGCAGCTGGATGTGGCTCTCTT
GCAGTTCATTTTTCCAGCAGTTTTTCCACTGCTACTGCCCTGTCGGATTTGGACGACGCA
CAGACCCCAATGGAGACTACATTAGGCGTTATTTGCCCATCTTGAAGGGCTTTCCAGCCA
AATACATTTATGATCCCTGGAATGCCCCAGAGGACGTGCAGAAGGCTGCAAAGTGTATCG
TAGGAGTCCACTACCCCAAGCCCATGGTGAACCATGCTGAGGCCAGCCGCATCAACATAG
AGCGGATGAAGCAAATTTACCAGCAGCTTTCAAGCTACAAAGGATTGGGCCTGCTGGCAA
CCATACCTTCCAATCCCCACAGTGGTGTCGGAGGAGTCAACACAGGAAGCAGTCACGGTG
TAGGAGGCTTCTCAACATACACAGTATCCCCTCAAACAGAAAGAGTACAGACCACAGAAA
AGAGGCGCCATGAAGAGGGTCCACATAGAGGCTCCAAATCGTTAAAGCAGAGCAAATAGT
GTGCAAAAGGTTGAATACAAAGGGAGCCCCAGTCATTCTGGATCTTTTGGATATCATCTT
TATATGCATTGTTATCAGCTGACACAGCAGCAGCTGTAACTAACAACCATAGATGACCAT
CATGATCTTCAAATAAGACATGTTATTAATCTAAGGAAAGGCGGAAGGCATTTAGACTAT
AATTGAAGATAAAGAACCCTGAAACAGCGGGATGTAAAATGCAAAACCATTTGATCTACA
GTATCCACATCAAGCTTCTGCTACAGTATGTTAAATGTTGATCGTTATCATCAATGCCCC
GCCCCCCCCATGACTGATGTAGGGCGTTGAGCGGGAAATTGGGTCATGAGGTCTTCTGTA
TGAAACTTTGTAACTTGTGTGAATGTGATCCATGCCTAAGTTCATCAAGACAATGAGCTC
AACTTTTAGCGGCTTTGGCCTTTACTTTTTCTTTAAAAAAATGTCTAAATCTGATTCCCA
TTCAAATGATGGGAACCTATAACACAATTTGGAGTTGCTTATCTGACCTGGGTAACATAC
AGTGGTGTCTTTCCTAAATATTGGGAAATATTAGGAAAAGAATGAATAAATATGTCATAA
ATGTACATTTGACAATGTACAAGCTCTGGACACTAAAATGAGATGTACAGTAATGAAAAT
GACAACCTCTCTGTCAGGTTTGACAATAAATAATGCCTTTTATAAATGCCAACAGACAGT
TTTTAATATTTATTATTTTTATTTCCAACTTTATATATTTGGTTGTGTGGTTTTACAACA
ACTTTTGTAAATAATGTGTTACGAAAACAAAAATCCACTGGATATGCATTTCAATTTCTC
CCATGTGATCGTAAATATGTTATTTTTGTATAAAGAATGAAAAGATTCTAACTGTATTGG
AATGGCAATAAATGACACATTGTCATAT

>emac_fin_isoseq_HQ_transcript/97523_cry2 mRNA_partial cds
GAAGTGCTGGTTGGTTTGACAGCAGGTGAAGTTTCTCGGGGGTCTACAAACACTGCCTGT
TTACATTAGACGATCTCTGGATTCTTTCTTATGATGGAATAAGGAGTCATCATGGTGGTA
AACTCTGTGCACTGGTTTCGCAAAGGCCTGCGGCTGCACGATAACCCGGTGCTGCAGGAG
GCCCTGACCGGGGCGGACACGGTGCGCTGTGTGTATGTCCTGGACCCGTGGTTCGCCGGC
GCCGCTAACGTGGGAATCAACAGATGGAGGTTTCTGCTTGAGGCTCTGGAGGACCTGGAT
AACAGTCTGAAGAAGCTCCACTCCAGACTGTTTGTAGTCAGAGGGCAGCCCACCGATGTT
TTCCCAAGGCTCTTTAAGGAATGGAAAGTGACCAGGTTGACATTTGAGTATGACCCAGAG
CCTTATGGGAAGGAGAGGGACGGGGCTATCATCAAGATGGCCCAGGAGTTTGGAGTGGAG
ACTGTTGTCAGAAACTCACACACCCTCTACAACCTGGACCGGATAATAGAGGCGAACAAC
AACAGCCCCCCCCTGACCTTCAAGCGCTTTCAGGCCATTGTGAGCAGACTGGAGTTGCCC
CGGAGACCGCTGCCCCCCATCACCCAGCAGCAGATGGACAAATGTTGCACTAAAATAGCT
GACAACCATGACCAGCTTTACAGTATACCTTCACTGGAAGAACTAGGTTTCAGGACAGCA
GGTTTACCTCCAGCTGTGTGGAAGGGAGGAGAGTCAGAGGCTTTGGACCGACTCAACAAA
CATCTGGACAAGAAGGTGTGGGTGGCCAACTTTGAGCACCCCCGTGTCAACACGTGCTCA
CTGTATGCCAGCCCCACTGGCCTCAGCCCCTACCTGCGCTTCGGCTGTCTGTCCTGCAGG
GTTCTGTACTACAACCTCCGAGAGCTCTACATGAAG
```

```
>emac_fin_Isoseq_HQ_transcript/25284_cry2 mRNA_partial cds
GGCTGCTCTACAGCTCCTGATTCATCCACAGTGTGTGTGTCCTCCCTGTATCCAGGGCTG
GAGGGGGGGGGGCACAGTCACCCATCACAGACTGCCTCACACACACTGCCAGCCCCAGCT
TCAACACACACCCAGAGCAGTCGGTCAAGGTCCAAACCCTCCTCCCCCATGTCTCTTTCC
CCGAGTCCAGCCCAGACCATGACTCAGCACTCCTCTCAGGGCCAGAGGAAGAAAAGCCCG
GCCTGCAAGGTGCGTCGCAGCCAGAGGCAGCAGGGGAGCAACACTCCATCCAGGGCGGGA
GAGGAGGAGATTGAGGAAGGAGGGGTGGAGGAGATGATGGAGGAGGAGGACGTTGAGCAG
GGTGAGGAGAGAATGGAGGAGGAGACGGCAGAGCCTCAGCAGTGATGTGAGGTGGAGGTC
AAAGAACAATGGTCCAACATCCATCTCAGGCAACTGGACTCATAAACACCTTCAGGGAGA
CGTGTTCTGGATATGTGGACAACACTTCTAAGCATGTGCTCAGCAGGAAGCAGAACTAAA
TCATTTTACTTGAGCTTACTCCATCCAGAACAAAATGGTAAAACAGTCTGGATAATAATA
GCTCCATGAAACCAAAGGGCCCCTTTTTAACATTTTATAACAGAACCAGTCAAGAAGGAT
CCAGAAAGTGCTTAGAAATACAAAATATTATTATTATTATCAATTATCAATTACGGTAAA
TTATTTTAAAAAAAAATTTTTTATTCAAGGAAATGGGGGTGATAATCCTAAAAGTTTTTC
AAGATGGTTGACCTTCTCTTGTAAAGTTCCACCTCTGACACCCACACTTAATAAGACACA
TTGTACTGCAACTACTCACACACACTATCCCAGTGGAACACAGTGTGACATTCAGCCCCT
CTCCGGTGCCTAGCTGTCCTTACCTCTGGAAGCAAGACGCAGGAGCCTGTCCCTGCTTTG
CTCCCCTCCCCTCCCCTCGATAGTGATTCAGAAAGAGAAGTAACTGTTCCAGTCTGAAAC
ATGTCCTTGGTGGTGGGTGAGCACAGAGGGATCACTTCATTCTGGGTCCCGCATAAAACT
GACCATTAGTATTATTTAGCGGCAGTAAAATATATCAAATGTCCCTTTATGTTGCAGCTC
CCTTCAACGATCAACTGTGGATGCTTTGAGCCTGCTTTACGAGGCACTGGATTATATTCC
AGCGGGAAAACAGTCTCGTCAATGTCAGACTTTATTTAATGTAAAGCCATTTAGGCCTCA
GGAACAGAATGCTTTTCTGTTGAATTGGCCCTGATTATTTACTTTGCCACATTTTAAAAA
CACCTAAGGAGAAAATTGAAATATACATATTAATATTGAAAGCACAATGTAAGTATTTTG
GAAAAAAAGTCAGATATTTTTGCTAGGGGCAATGAAAATGTTTTATAGTGACGGGGAACA
TTTCTATAGACTACCTCCCTTAATCTGTCTGTTTTGGAGTCACATATACAGCCGCGGAAA
AAAATGAAGAGACCACTTCAGTATTATCATTTTCTCCTATTTTATTATTATTAGGTTTGT
CTTTGAGTTAAAAAAAAATGTTTTAATTTAAAAATAGTTCAATTATTTGTATTGTATTTTA
TTGTATTCTAAAAACTCCTGACGATTTTTCTCTGTGTTGGAATTCAACAGACACTGGAAT
GGCTGCCATACATGTAGAGATAAATATTTAAGAAAAATGTGGAGTAGTTTCCGCGGCTGT
ATATAAAATATATGAAACATTTAAGGTCAAATTAAAACCGTATGTATTACGTGTGTACCT
TATGCTTTTTGTCTCTGACAATTAAAAATGCTGCATGTGTAGAATGAATGTAGAACGAAA
AACGACGTGTTGTGCTCCCCATTTATTTATAAGAGTTTGTATTGAAATTATATGTAAAAT
CATAACCTGTGCTTCTGGTTTAAAGGGATTTTACCAATTTTCCTACCAGAAGCATCCACC
AGAGGGCAGCAGATGCAGTCATTTCCGCACATGTTTAGTTTTTGTACATTTACAAAAGCT
CCAACTCATTTCTTCACAATCCTGATTTGTTTCCTGTGATAAATGCACTAAATGCTATCA
GATACTGAAACCTGCCATTTCAATGTGTCTTTAGATGTCCTGTTTAGTGAACCCAGAGTC
CAAAACAAGATTCAGAGACCTTCACAGCCAAAAACAACAAATGACTTCATTACATTTTCG
TTCTGATAACTGACATTTAACTGATTAAAAAAATGTTTTTCTAAATGTAACTTCTTTTTG
TTTCAGCATTAACATACTGTACTGTCCTCCGCATTTCAAAGAGTGACAATCTGTTTGCTT
ATTCCTACCTCATACCTTTCTGTTTGACATTTTCCACTTCTAGCCTGAAAAACTGAAGCG
TTGAATCTTGTTAGGTGTGTTGTGTCAATGGATTTCATGTAACTGTGTTAGGTTAGTTGA
AAGCAATGATATTAAGTTGGACCTAATGTTTTTTATTTAAACTTGATATTATTGCTTTCG
ACTTAATAGAGAGATGTGAAATCGATTTACATAATACATTTGAGTTAAATTCAACGTTTC
AGTTCTTTCAGTGTAACTTCAGTGCAGAATAAGCCACCAAACCTTTGGCTGAGAGACCCT
CAGTGCTGCCTGGACGGTTCCAGGATGTTGATGTTCAGAAAAAAACTTCTCAGCACTGGC
AGTTCCTCAGTCGATTTAAAGGACGAGTGGGAACTATGACGTTTAAACGAAAAGCTCACA
GTTCCCACTGTTCCTAAACATCAACAGCTCCCTGACCAGGGGATTTGTATACGTTCCACC
TGCCACCATGATGCTCTGCAGCCGGGGTGTGTCTGACTGTGTGGATATTTATGTGTTTGG
TTTTTTTTGTACAGAAATAGTTTATTTTCATTAAAATTGCTGAAGGAAAGTGATTG

>emac_TRINITY_DN52132_c1_g6_i2_cry2 mRNA_partial cds
CTTCTGTCCACGCCCCCTAAGTGCTGGTTGGTTTGACAGCAGGTGAAGTTTCTCGGGGGT
CTACAAACACTGCCTGTTTACATTAACGATCTCTGGATTCTTTCTTATGATGGAATAAGG
AGTCATCATGGTGGTAAACTCTGTGCACTGGTTTCGCAAAGGCCTGCGGCTGCACGATAA
CCCGGTGCTGCAGGAGGCCCTGACCGGGGCGGACACGGTGCGCTGTGTGTATGTCCTGGA
CCCGTGGTTCGCCGGCGCCGCTAACGTGGGAATCAACAGATGGAGGTTTCTGCTTGAGGC
TCTGGAGGACCTGGATAACAGTCTGAAGAAGCTCCACTCCAGACTGTTTGTAGTCAGAGG
GCAGCCCACCGATGTTTTCCCAAGGCTCTTTAAGGAATGGAAAGTGACCAGGTTGACATT
TGAGTATGACCCAGAGCCTTATGGGAAGGAGAGGGACGGGGCTATCATCAAGATGGCCCA
GGAGTTTGGAGTGGAGACTGTTGTCAGAAACTCACACACCCTCTACAACCTGGACCGGAT
AATAGAGGCGAACAACAACAGCCCCCCCCTGACCTTCAAGCGCTTTCAGGCCATTGTGAG
CAGACTGGAGTTGCCCCGGAGACCGCTGCCCCCCATCACCCAGCAGCAGATGGACAAATG
TTGCACTAAAATAGCTGACAACCATGACCAGCTTTACAGTATACCTTCACTGGAAGAACT
AGGTTTCAGGACAGCAGGTTTACCTCCAGCTGTGTGGAAGGGAGGAGAGTCAGAGGCTTT
GGACCGACTCAACAAACATCTGGACAAGAAGGTGTGGGTGGCCAACTTTGAGCACCCCCG
TGTCAACACGTGCTCACTGTATGCCAGCCCCACTGGCCTCAGCCCCTACCTGCGCTTCGG
CTGTCTGTCCTGCAGGGTTCTGTACTACAACCTCCGAGAGCTCTACATGAAGCTGCGTAA
GCGCTGCAGTCCTCCTCTCTCTCTGTTTGGACAGCTGTTGTGGAGGGAGTTCTTCTACAC
```

AGCCGCCACCAACAACCCCAACTTTGACCGCATGGAGGGAAACCCCATCTGTGTGCAGAT
CCCGTGGGACCAGAACCCAGAGGCGCTGGCCAAGTGGGCTGAGGGTCGGACTGGTTTCCC
CTGGATCGATGCCATCATGACCCAGCTGAGACAGGAGGGCTGGATCCACCACCTGGCCCG
GCACGCCGTGGCCTGTTTCCTAACCAGGGGGGACCTTTGGATCAGCTGGGAGAGCGGCAT
GC

>emac_TRINITY_DN62964_c2_g6_i2_cry2 mRNA_partial cds
TTCTTCCAGCAGTTCTTCCACTGCTACTGCCCTGTGGGCTTTGGAAGGAGGACCGACCCG
TCCGGAGACTTCATCAGGCGTTACATCCCCATCCTGAAGCACTACCCCAACCGCTACATC
TATGAGCCGTGGAACGCTCCGGAGTCCGTCCAAAGGGCAGCCAACTGCATCGTGGGAGTG
GATTACCCCAAACCTATGATCAACCACGCAGAGGGCAGCAGGCTCAACATCGAGAGGATG
AAACAAGTGTACCAGCAGCTCTCCCACTACAGAGGCCTCAGTCTACTAGCATCAGTACCA
ACGATCCAAGAGGAGGCAGAGCCACCAATGAGCGATGAATCCCAGAGCAGCAGTGGCGCT
GACTCTCCTCCCAGAAGCCCTGCTGACAGCGAGGCAGCCGGCTGCTCTACAGCTCCTGAT
TCATCCACAGTGTGTGTGTCCTCCCTGTATCCAGGGCTGGAGGGGGGGGGGCACAGTCAC
CCATCACAGACTGCCTCACACACACTGCCAGCCCCAGCTTCAACACACACCCAGAGCAGT
CGGTCAAGGTCCAAACCCTCCTCCCCCATGTCTCTTTCCCCGAGTCCAGCCCAGACCATG
ACTCAGCACTCCTCTCAGGGCCAGAGGAAGAAAAGCCCGGCCTGCAAGGTGCGTCGCAGC
CAGAGGCAGCAGGGGAGCAACACTCCATCCAGGGCGGGAGAGGAGGAGATTGAGGAAGGA
GGGGTGGAGGAGATGATGGAGGAGGAGGACGTTGAGCAGGGTGAGGAGAGAATGGAGGAG
GAGACGGCAGAGCCTCAGCAGTGATGTGAGGTGGAGGTCAAAGAACAATGGTCCAACATC
CATCTCAGGCAACTGGACTCATAAACACCTTCAGGGAGACGTGTTCTGGATATGTGGACA
ACACTTCTAAGCATGTGCTCAGCAGGAAGCAGAACTAAATCATTTTACTTGAGCTTACTC
CATCCAGAACAAAATGGTAAAACAGTCTGGATAATAATAGCTCCATGAAACCAAAGGGCC
CCTTTTTAACATTTTATAACAGAACCAGTCAAGAAGGATCCAGAAAGTGCTTAGAAATAC
AAAATATTATTATTATTATCAATTATCAATTACGGTAAATTATTTAAAAAAACAATTTTT
TATTCAAGGAAATGGGGGTGATAATCCTAAAAGTTTTTCAAGATGGTTGACCTTCTCTTG
TAAAGTTCCACCTCTGACACCCACACTTAATAAGACACATTGTACTGCAACTACTCACAC
ACACTATCCCAGTGGAACACAGTGTGACATTCAGCCCCTCTCCGGTGCCTAGCTGTCCTT
ACCTCTGGAAGCAAGACGCAGGAGCCTGTCCCTGCTTTGCTCCCCTCCCCTCCCCTCGAT
AGTGATTCAGAAAGAGAAGTAACTGTTCCAGTCTGAAACATGTCCTTGGTGGTGGGTGAG
CACAGAGGGATCACTTCATTCTGGGTCCCGCATAAAACTGACCATTAGTATTATTTAGCG
GCAGTAAAATATATCAAATGTCCCTTTATGTTGCAGCTCCCTTCAACGATCAACTGTGGA
TGCTTTGAGCCTGCTTTACGAGGCACTGGATTATATTCCAGCGGGAAAACAGTCTCGTCA
ATGTCAGACTTTATTTAATGTAAAGCCATTTAGGCCTCAGGAACAGAATGCTTTTCTGTT
GAATTGGCCCTGATTATTTACTTTGCCACATTTTAAAAACACCTAAGGAGAAAATTGAAA
TATACATATTAATATTGAAAGCACAATGTAAGTATTTTGGAAAAAAAGTCAGATATTTTT
GCTAGGGGCAATGAAAATGTTTTATAGTGACGGGGAACATTTCTATAGACTACCTCCCTT
AATCTGTCTGTTTTGGAGTCACATATACAGCCGCGGAAAAAAAATGAAGAGACCACTTCAG
TATTATCATTTTCTCCTATTTTATTATTATTAGGTTTGTCTTTGAGTTAAAAAAATGTTT
TAATTTTAAAATAGTTCAATTATTTGTATTGTATTTTATTGTATTCTAAAAACTCCTGAC
GATTTTTCTCTGTGTTGGAATTCAACAGACACTGGAATGGCTGCCATACATGTAGAGATA
AATATTTAAGAAAAATGTGGAGTAGTTTCCGCGGCTGTATATAAAATATATGAAACATTT
AAGGTCAAATTAAAA

>emac_fin_isoseq_HQ_transcript/2988_cry3a
GGTGAGCCAAACTCTGATGAAGGTAGTGCAGCAGCAGCAGCAGAATTCAACACATTCATA
GAGTGCTTCTAGGCTCACAGCACTTTCCCGTACATCGTAACACTTGGAAGAAATGCGTCT
TTCTTTGGACTTGACATAAATATCAACTACAAGGAGCCAGCATCAGAACAAA ATGGCCCC
AAATTCCATCCACTGGTTCAGGAAAGGCCTGCGTCTCCATGACAACCCATCACTGCTGCA
GGCGGTCAAAGGAGCAGGCACCGTGCGCTGTGTTTACTTCCTGGATCCTTGGTTTGCAGG
CTCGTCCAACGTCGGTGTCAACAGGTGGAGGTTTCTCCTTCAGTGTTTGGAGGATCTTGA
CGCCAGCCTTCGGAAGCTTAACTCTCGCCTTTTTGTCATCAGAGGCCAACCAGCCAACGT
GTTCCCACGGCTCTTTAAGGAGTGGAAGATCTCTCGCCTGACCTTTGAGTATGATACGGA
GCCGTTTGGGAAGGAGAGAGACGCTGCCATCAAGAAGCTGGCCATGGAGGCAGGGGTGGA
GGTCGTCGTCAAGACGTCACACACCCTCTACGATCTGGACAAGATCATAGAGCTGAATGG
CGGACAGCCACCACTCACCTACAAGCGTTTCCAGACTCTGATCAGTCGAATGGATCCTCC
TGAGATGCCCGTGGAGACGCTGTCGGACACCCTGATGGGTCGTTGCGTCACCCCCGTCGC
AGAGGACCACGGAGAAAAGTACGGAGTCCCTTCCCTGGAGGAGCTAGGCTTTGACACCGA
GGGCCTGCCCACAGGCGTTTGGCCCGGAGGAGAGACAGAGGCTCTGACAAGGATAGAGCG
CCATCTGGAGAGAAAGGCGTGGGTAGCTAACTTCGAGCGTCCCAGAATGAATGCCAATTC
GTTGCTCGCCAGCCCGACTGGCCTCAGCCCATACCTGCGCTTCGGCTGCCTCTCCTGTCG
CCTTTTCTACTTCAAGCTCACCGACCTCTACCGCAAGGTGAAAAAGAACAGCTCCCCTCC
ACTCTCTCTGTACGGCCAGCTGCTGTGGCGAGAGTTCTTCTACACCACAGCAACCAACAA
CCCACGCTTCGACAAGATGGAGGGAAACCCGATCTGCGTGCGCATCCCCTGGGACAAAAA
TCCTGAAGCCCTTGCCAAGTGGGCGGAGGCTAAGACAGGATTTCCCTGGATAGACGCCAT
CATGACTCAGCTGCGGCAGGAGGGCTGGATCCATTACCTGGCCCGGCATGCTGTGGCTTG
CTTCCTCACAAGGGGAGACCTTTGGATCAGCTGGGAGGAGGGGATGAAGGTGTTTGAGGA

```
GCTGCTTCTTGATGCAGACTGGAGCGTGAACGCAGGCAGCTGGATGTGGTTGTCCTGCAG
TTCATTTTTCCAGCAGTTCTTCCACTGCTACTGCCCCGTGGGCTTCGGCCGCCGGACCGA
CCCCAACGGGGACTTCATTAGACGATACTTACCTGTCCTCCGAGGTTTCCCCGCTAAGTA
CATCTACGACCCGTGGAACGCTCCGGAGTCCGTGCAGGCAACCGCCAAGTGCATCATCGG
CGTCCATTACCCGAAGCCCATGGTGCATCACGCAGAGGCGAGCCGGCTCAACATCGAGAG
GATGAAGCAGATCTACCAGCAGCTGAGCCGATACAGGGGACTGAGCCTGCTGGCATCCGT
TCCGTCCTCCAATGGGAACGGGAACGGAGGAATGATGGCCTACCCCCTCGGAGAGCAGCA
GCCAGGGTCCAACAACAACAATTCACATTTGCCTGGAGTGTCGGGGAGTTCGGTTTCAAC
GAGTAACGGCAGCGGGAACATCCTCAACTTTCAAAATGAAGAAGACATGGGGCCTAGCAG
CAGACAACAACAGCAGCAGCAGCATGGATACCCCTCAGTGCAAGAAGCCAGCCAGACCAT
CAGCAGCAGCCGACTCTACCACGAGTTCGCTGTGCCTCAACACCCAGGATTTCTGCACAG
CAGAGGAAGCATTACAGGAAAGAGGGAGCGTGAATCGGAACGAGAAGGGGCGGGGGAGAA
AGACCCGGCGTCCTGCTCCGTGCACAAGATGCAGAGGCAGAGTGCAGAGACTACCTAGAG
CAGCTGTTGATGAGGATTACTTTGAAATCACCATCCAACCAGCGGAGAGAGGGAATCCAA
ACATGGTCACCTGTCCTCCAAACACGCGCTGACATCACTGCCTCGAACTTTATGCACCTA
CTCTCTGTCTTTAACGTCAGAGATTCCCAGGGGTGCGAACGCAGAGTTGAAAATGTGACT
TTTGACACTTTTTCTACAAGTCCATTGACCTTTGACTTCTTGAAGACAGGTGTGAAGACG
CAAGAGGAATGTGCAAAGTAACGAAACACCTCGAAACTTACACAGGAAAAATGATTCGTA
AAAACAAATATGTACATTATTGTCTTAAAACCGCTGCCACCAGTCAGTACATACTGTGTG
AACTCTGAAGCAGCTATACATGTCAGGTATAGAACAATTCAACTCTTTTTTTCAGAGACC
TTTTTGTACTTTCACGCAGTTTGTAATAATGCTCACCAGGATAACATATCACCTCCTCGC
GTTCCCTTTCACTACAGGAATATTTCTAGTCACTAGTGTTTAATCTTAATATTTGAAGAA
ATTATATTTATACTGATATTTATTGTGCGCTTGTAAATGTGTGAGAAAAAGAGGACAGTG
ATTTTAGACCTTGATTTCACTGGGCAAAATGACACAAAAAGTAATACCGTAATTAAGTAA
AGTTTCGATGTCAATGGGGATTAAGGCACTGGAGGTTCAGTTACTGAGTGAGCAGGAAAT
CCCTCTGAATATCAACGTGACTTTTCACAGTTGATATAATTAACACAAGTATTCATTTCA
TTTGAAACCATCCAATAAACAAAGAAAAGGTTTTCAGATTGTATGTCACCCGTATCAGGG
ATTTAAAAAAAAAAACTCAGACCGGTTTTTATTGTTGATCTGTAAAAACATGTAATTAGT
TAGGAGACATTCAGATGTGTCACTCTGGCTGTAAAACTGCCTTTTTGGAGTGTGGCATTG
TAGGTAAAGGAGTTCTTAATAGCTTCCCTGGTAGTCCTGTGTTCCATTGTGTGAGATTAT
CAGACACATAGATGAAACCTTCAATTAAAATATATATTTTTTTCCATTTCTATGAAAACC
ATGACGGTCTTGAGCCTTTGTCGCTTTTAACCGAACACACCGATCGATGAAACATGATCA
GGCTCTGTTCTGTGCTGTTACTGAGATTTCCTTTTCTATCACATGTTTTATTCTGAGTTT
CCAATGTTGCTTTTTCATTGTAAATTATTAATTATGTACAGATCCGCTCTATGGTTTCAC
GATGATTTGTTATAGAATTATTCAGATTTTCACAGCTGTTATATTTCTGTGATATGTTGA
TTTGACTCAACGCTTAGAAAGTCACTTGCATTCTCACCTCTTTGTTTCCCCCAGAAATCT
TCAGGGTTTCTAAAAGAGTAAAAGAAAATTGTATAACTTTGGAGTATCTCTAAGGCATTC
TGTGCTGAAATTTGAACATTTTTGCTGTGCAACACGTTCAACAACCAAAAACATGTGGTG
ATTCATTCGCGAGAGGACGTGGATGTTACATCTGTCAAATACCATGAAGACAGGGATCAG
TTCACTGTAAACAATGGGGTTTTTTTTCTAAGATTTTGAGGTACAAAAAGGAGGAAAAAC
GATTTAAAAAGAGGGAAACGACATTCCTCAAAAACCTTGTACCCTCCTGAAGACTGCTTC
ACAGGGAGTTATTTTATTAGTTACTTTATGTATTGCTTCTTGCTAAATTTTGCAGCAATT
AGTAAATACAAAATGTCAAATAACTGTTTCAGAGAGCTGTACAGCATTGGACGCTGTACA
GTCTATTGGAGCGCTGTACAGTCTATTGGAGGATTGTGTCTTATTTAGTTTCTACCCAAC
CCTCGATCATCACCTTTACTTCTGATGAGAACTAATCCAAGGAATGTATGATTGGCCACA
CCTTTCTGTTTCTCTCTCACACACACACACACACACACACACACACACAAACACACACTA
AACATTGATAACTAGTGCATACACAAACAGATACATCTACTGTAAATCCTGACATTAAAA
GCAGTCTAAATGTAACATTATTTAAAGTGATCCCTTACAATTGGTCCTGATTGATTATAT
ACATCTGTATCATGAATCTGTAAGTCCTTAAAAGGATTTTGGAACATTATTAGTTTATTT
TAAGTCGAACCAAATTCTTTTAATTTGTTTTTTTCTTTGTTTTGTTCTCCTCGTTGTGAC
ATGAGCAGCTGCCATCTATCATCTCTGTAATATGTATCTCTTTCTTCGATTCTCCGTGTG
TTGTGAAACACTTTTGTTATGCTTGATTAAGTTCCTGACAGGAAATAAAAACTGTAACAT
ACTG

>emac_fin_isoseq_HQ_transcript/49232_cry3a
GACATTCATAGAGTGCTTCTAGGCTCACAGCACTTTCCCGTACATCGTAACACTTGGAAG
AAATGCGTCTTTCTTTGGACTTGACATAAATATCAACTACAAGGAGCCAGCATCAGAACA
AAATGGCCCCAAATTCCATCCACTGGTTCAGGAAAGGCCTGCGTCTCCATGACAACCCAT
CACTGCTGCAGGCGGTCAAAGGAGCAGGCACCGTGCGCTGTGTTTACTTCCTGGATCCTT
GGTTTGCAGGCTCGTCCAACGTCGGTGTCAACAGGTGGAGGTTTCTCCTTCAGTGTTTGG
AGGATCTTGACGCCAGCCTTCGGAAGCTTAACTCTCGCCTCTTTGTCATCAGAGGCCAAC
CAGCCAACGTGTTCCCACGGCTCTTTAAGGAGTGGAAGATCTCTCGCCTGACCTTTGAGT
ATGATACGGAGCCGTTTGGGAAGGAGAGAGACGCTGCCATCAAGAAGCTGGCCATGGAGG
CAGGGGTGGAGGTCGTCGTCAAGACGTCACACACCCTCTACGATCTGGACAAGATCATAG
AGCTGAATGGCGGACAGCCACCACTCACCTACAAGCGTTTCCAGACTCTGATCAGTCGAA
TGGATCCTCCTGAGATGCCCGTGGAGACGCTGTCGGACACCCTGATGGGTCGTTGCGTCA
CCCCCGTCGCAGAGGACCACGGAGAAAAGTACGGAGTCCCTTCCCTGGAGGAGCTAGGCT
TTGACACCGAGGGCCTGCCCACAGGCGTTTGGCCCGGAGGAGAGACAGAGGCTCTGACAA
GGATAGAGCGCCATCTGGAGAGAAAGGCGTGGGTAGCTAACTTCGAGCGTCCCAGAATGA
```

ATGCCAATTCGTTGCTCGCCAGCCCGACTGGCCTCAGCCCATACCTGCGCTTCGGCTGCC
TCTCCTGTCGCCTTTTCTACTTCAAGCTCACCGACCTCTACCGCAAGGTGAAAAAGAACA
GCTCCCCTCCACTCTCTCTGTACGGCCAGCTTCTGTGGCGAGAGTTCTTCTACACCACAG
CAACCAACAACCCCCGCTTCGACAAGATGGAGGGAAACCCGATCTGCGTGCGCATCCCCT
GGGACAAAAATCCTGAAGCCCTTGCCAAGTGGGCTGAGGCTAAGACAGGATTTCCCTGGA
TAGACGCCATCATGACTCAGCTGCGGCAGGAGGGCTGGATCCATCACCTGGCCCGGCATG
CTGTGGCTTGCTTCCTCACAAGGGGAGACCTTTGGATCAGCTGGGAGGAGGGGATGAAGG
TGTTTGAGGAGCTGCTTCTTGATGCAGACTGGAGCGTGAACGCAGGCAGCTGGATGTGGT
TGTCCTGCAGTTCATTTTTCCAGCAGTTCTTCCACTGCTACTGCCCCGTGGGCTTCGGCC
GCCGGACCGACCCCAACGGGGACTTCATTAGACGATACTTACCTGTCCTCCGAGGTTTCC
CCGCTAAGTACATCTACGACCCGTGGAACGCTCCGGAGTCCGTGCAGGCGACCGCCAAGT
GCATCATCGGCGTCCATTACCCGAAGCCCATGGTGCATCACGCAGAGGCGAGCCGGCTCA
ACATCGAGAGGATGAAGCAGATCTACCAGCAGCTGAGCCGATACAGGGGACTGAGCCTGC
TGGCATCCGTTCCGTCCTCCAATGGGAACGGGAACGGAGGAATGATGGCCTACCCCCTCG
GAGAGCAGCAGCCAGGGTCCAACAACAACAATTCACATTTGCCTGGAGTGTCGGGGAGTT
CGGTTTCAACGGGTAACGGCAGCGGGAACATCCTCAACTTTCAAAATGAAGAAGACATGG
GGCCTAGCAGCAGACAACAACAGCAGCAGCAGCATGGATACCCCTCAGTGCCAGAAGCCA
GCCAGACCATCAGCAGCAGCCGACTCTACCACGAGTTCGCTGTGCCTCAACACCCAGGAT
TTCTGCACAGCAGAGGAAGCATTACAGGAAAGAGGGAGCGTGAATCGGAACGAGAAGGGG
CGGGGGAGAAAGACCCGGCGTCCTGCTCCGTGCACAAGATGCAGAGGCAGAGTGCAGAGA
CTACCTAGAGCAGCTGTTGATGAGGATTACTTTGAAATCACCATCCAACCAGCGGAGAGA
GGGAATCCAAACATGGTCACCTGTCCTCCAAACACGCGCTGACATCACTGCCTCGAACTT
TATGCACCTACTCTCTGTCTTTAACGTCAGAGATTCCCAGGGGTGCGAACGCAGAGTTGA
AAATGTGACTTTTGACACTTTTTCTACAAGTCCATTGACCTTTGACTTCTTGAAGACAGG
TGTGAAGACGCAAGAGGAAGGTGCAAAGTAACAAAACACCTCGAAACTTACACAGG

>emac_fin_isoseq_HQ_transcript/71427_cry5
GGTTTTATGAAACAAATCAACGTGTTGGTTACCAAACGCGCAACTCTCAGTAAACCAAGC
GGAATTCTAACTGTTGTGAGGCACAATAACGGCATGGCTCATTCTTGTATTCACTGGTTC
CGCAAGGGACTCAGGCTACATGACAACCCAGCCCTGATGGCTGCTCTGAGGGACTGTAAG
GAGCTGTACCCCGTGTTTATCCTGGACCCTTACCTATATAACAACACCCTTGTGAGTATC
AACCGCTGGAGGTTTCTCATTGGATCCCTCAAAGACCTGGACCGCAGCCTCAGGAAACTC
AACTCCAGGCTGTTTGTTGTGAGAGGGAAGCCAGAGGAAGTGTTCCCTAAGCTGTTCAAG
AAGTGGGACGTCACAAAGTTAACCTATGAGTACGACACAGAGCCCTACAGCCTGAGCCGG
GACAGAACAGTGACCACACTGGCCAAAGAACATGAAGTCGAAGTCATCTACAAAATCTCA
CACACTATTCATGATATGGACAGGATAATTGAGGAAAACAACGGGAAGGCTCCCCTTACT
TATAACCGGTTTCGAACAATATTGAAGACTATAGGTTCCCCTAAGAAACCCATTCCTGCT
CCAGCCATCGAAGATATTAAAGATGTGAAGACGCCTTGTTCAGAAAACCATGAGAAAGAT
TATGGGATACCTACTCTGGAGGAACTCGGTCATGACACCACAGCTCTTGGAGAAGAGAAG
TTCCCAGGAGGAGAACAAGAGGCTCTGAGGAGATTAGATGAACACATGGAAAGAAAGGAA
TGGGTGTGTAAGTTTGAGAAGCCTCAGACTTCTCCGAACTCTCTGAGCCCCAGCACCACT
GTCCTCAGTCCGTACGTCACTTTTGGCTGCCTGTCCGCACGCACCTTTTGGTGGAGGGTG
ACAGACGTCTATCAGGGGAAGAAGCACTCAGAGCCTCCTGTTTCCCTGCATGGCCAGCTT
CTGTGGAGAGAGTTTTACTACGCCGCTGGTGTGGGTATCCCTAACTTTGACAAAATGGAG
GGCAACCCTGTTTGTACCCAGGTGGATTGGGACACAAACCCTGAATATCTTGCTGCATGG
AGAGAGGCTCGGACTGGTTTCCCTTTCATCGACGCCATCATGACTCAGCTGAGGCAGGAG
GGCTGGATCCACCACCTGGCCAGACATGCTGTCGCCTGTTTTCTCACCAGGGGAGACCTG
TGGATCAACTGGGAAGAAGGGCAGAAGGTGTTCGAGTATCTTTTATTGGATGGCGATTGG
GCCCTGAATGCTGGAAACTGGCAGTGGCTCTCAGCGAGTGCCTTCTTCCATCAGTACTTC
AGAGTTTACTCGCCTGTTGCCTTTGGCAAGAAGACAGACAAAAATGGAGATTACATCAAA
AAGTACCTTCCTCTTCTGAAAAAGTTCCCAGCCCAGTATATCTATGAGCCTTGGAAAGCT
CCGTGCAGTGTCCAGCAGGCAGCAGGTTGCATTGTGGGTAAAGACTACCCGCAACCTATT
GTTAAGCATGAAATGATCAGCAAGCAAAACATCCAGAGGATGAAGTTGGCTTATGCTAAG
AGATCCAAAGATTCTGCTGAATCACCAAACGAATCACCCAGCAAAATACAAGGTGTCAAG
CGAAAGGCTAAATCCGTCATTGACATGCTGCAAAAGAAAAAGAAAAACTAATAATGGTAC
ACAAACAAACGGGAAGCGATGTCTGGGCTGAAAGGTTGGGACACTTTTTTGTGTATGCAG
ACAGTGTACTATGTCTACCTCATCTCCTTTAACTAGTTACTGTTTTATATTGTATTTATG
TCAATGTTTTTGTAAAAAGCGAGTGAAAGGGTATACTTAAAAAAAATATATTCTTGCATG
GGGATTGAGTTAAAATGTGTAAAGATGAATGGTCTCGTATTGTATGAAGACGAGAGATTT
TGAGGCATCATATTGTTGCACCTTAAAGATTGAATGTTTATTTTTTTACGGTTTGTTTTT
AATATCCAATATAAATAAATTCATACATATT

>emac_fin_isoseq_HQ_transcript/72338_cry-dash mRNA_complete cds
GTCACACGGAGCAGCTACACACCGGGAGGAGCTGCTCTCTCACACCGAGCACTTCTGTGC
CTAATATAACTATTAGTTACACTAGTATATAATATATAAGTATGTCTACCTCGCGTACAA
TCATATGTTTACTGAGAAACGACTTGCGTCTGCACGACAACGAGCTTTTCCACTGGGCTC
AAAGAAATGCAGAACACATTGTCCCTCTGTACTGCTTCGACCCAAGACATTATGTGGGGA
CGTATAACTACAACCTGCCAAGGACCGGGCCTTTCCGCCTGCGTTTCCTACTGGACAGCA
TCAGGGACCTCAGACACACGTTGCTCAACAAGGGCAGCAACCTGGTAGTGAGACGGGGTA

```
AACCAGAGGAGGTCGTTGCGGACCTCATCAAGCAGCTGGGCTCCGTCAGTACCGTGGCTT
TCCATGAAGAGGTGACTTCGGAGGAACTGAATGTGGAGAAACGAGTGAAGGATGTCTGCG
CACGGATGAAGGTCAAAGTTCACACCTGCTGGGGGTCCACACTGTACCACAGAGACGATC
TTCCATTCAACCACATATCCAGGCTGCCTGATGTGTACACTCAGTTCAGGAAGGCGGTGG
AGACCCAGAGCCGGGTGAGACCTCTGATACCGACACCTGAGCAGCTGAAGACTCTGCCCC
AAGGGCTGGAGGAAGGAGCCATTCCCACAGCAGAGGACCTGGAACAAACAGAGCCTGTGT
CTGATCCCCGCTCAGCCTTCCCCTGCAGTGGTGGGGAGAGTCAAGCTTTGGCCAGACTCA
AACACTATTTCTGGGACACTAATGCAGTTGCAACCTACAAGGAGACTCGCAACGGCTTGA
TCGGTGTGGATTATTCCACTAAATTTGCACCTTGGCTGGCGCTGGGTTGCATCTCACCCA
GGTACATCTATCATCAGATCCAGCAGTATGAGAAGGAGCGGACAGCCAATCAGAGCACAT
ACTGGGTCATTTTTGAACTTTTGTGGAGGGATTACTTCAAGCTTGTAGGGGTCAAGTACG
GGAACAGACTGTTTCAGATCAAAGGACTTCAAGACAAATCTGTTCCATGGAAGAAGGACA
TGAAGCTTTTCGACGCATGGAAAGAGGGACGCACAGGAGTGCCCTTTGTGGACGCAAACA
TGAGAGAGTTGGCCATGACGGGCTTCATGTCCAACAGGGGGCGGCAAAATGTTGCCAGCT
TCCTCACAAAGGACCTGTGCCTGGACTGGAGGATGGGAGCGGAGTGGTTTGAATACCTGC
TGATTGACCATGATGTCTGCAGCAACTATGGGAACTGGCTGTACAGCGCGGGCATTGGAA
ACGACCCCAGAGAGAACAGGAAGTTCAACATGATCAAGCAGGGCCTGGACTACGACAACA
ATGGTGACTATGTGAGGCAGTGGGTGCCTGAGCTGCAGGGGATCAGGGGAAGCGATGTAC
ACACACCCTGGACCCTCAGCACTGCGTCGCTGTCACACGCCAACGTCTCCCTCGGTGAAA
CCTACCCAACCCCCATCGTCATGGCGCCTGAATGGAGCCGACACGCTAACAAGAAACCGA
GTGGCGCTGGGCCTTCATCACGAGGAAAGAAAGGCCCGTCTCACACTCCCAAACAACACC
GTGACAGAGGAATAGATTTCTATTTCTCCAAAAGCAAAAACCTTTGACCCTCACAGTGAA
CCCAGGAGGGAAACTCCAGTGATTCTAACTTGGACTGCTGCTGGACAAAATCGGTGTTGC
CTGATGAGGATTTTATCAAATCACAAAGCGCCTTTTATGGATGTGCTTCCTGTTCGACTG
CAGTAACGATGAAGCAAAACTGTGGTTTTCATGCAGTCAGATGATGCTTTGATGCAAGTT
TTAATGTGGTTTTCTTTTGAAGTTAGCTTTTTTATGTGTGTTACCCAATAAAATGGAAAT
ATTAAGTTGTATAAGCGGTCGTGATTTTGTTATGCGATGCAAATAAAAATGTTTTTGAC

>emac_fin_isoseq_HQ_transcript/76532_cry-dash mRNA_complete cds
GTCACACGGAGCAGCTACACACCGGGAGGAGCTGCTCTCTCACACCGAGCACTTCTGTGC
CTAATATAACTATTAGTTACACTAGTATATAATATATAAGTATGTCTACCTCGCGTACAA
TCATATGTTTACTGAGAAACGACTTGCGTCTGCACGACAACGAGCTTTTCCACTGGGCTC
AAAGAAATGCAGAACACATTGTCCCTCTGTACTGCTTCGACCCAAGACATTATGTGGGGA
CGTATAACTACAACCTGCCAAGGACCGGGCCTTTCCGCCTGCGTTTCCTACTGGACAGCA
TCAGGGACCTCAGACACACGTTGCTCAACAAGGGCAGCAACCTGGTAGTGAGACGGGGTA
AACCAGAGGAGGTCGTTGCGGACCTCATCAAGCAGCTGGGCTCCGTCAGTACCGTGGCTT
TCCATGAAGAGGTGACTTCGGAGGAACTGAATGTGGAGAAACGAGTGAAGGATGTCTGCG
CACGGATGAAGGTCAAAGTTCACACCTGCTGGGGGTCCACACTGTACCACAGAGACGATC
TTCCATTCAACCACATATCCAGGCTGCCTGATGTGTACACTCAGTTCAGGAAGGCGGTGG
AGACCCAGAGCCGGGTGAGACCTCTGATACCGACACCTGAGCAGCTGAAGACTCTGCCCC
AAGGGCTGGAGGAAGGAGCCATTCCCACAGCAGAGGACCTGGAACAAACAGAGCCTGTGT
CTGATCCCCGCTCAGCCTTCCCCTGCAGTGGTGGGGAGAGTCAAGCTTTGGCCAGACTCA
AACACTATTTCTGGGACACTAATGCAGTTGCAACCTACAAGGAGACTCGCAACGGCTTGA
TCGGTGTGGATTATTCCACTAAATTTGCACCTTGGCTGGCACTGGGTTGCATCTCACCCA
GGTACATCTATCATCAGATCCAGCAGTATGAGAAGGAGCGGACAGCCAATCAGAGCACAT
ACTGGGTCATTTTTGAACTTTTGTGGAGGGATTACTTCAAGCTTGTAGGGGTCAAGTACG
GGAACAGACTGTTTCAGATCAAAGGACTTCAAGACAAATCTGTTCCATGGAAGAAGGACA
TGAAGCTTTTCGACGCATGGAAAGAGGGACGCACAGGAGTGCCCTTTGTGGACGCAAACA
TGAGAGAGTTGGCCATGACGGGCTTCATGTCCAACAGGGGGCGGCAAAATGTTGCCAGCT
TCCTCACAAAGGACCTGTGCCTGGACTGGAGGATGGGAGCGGAGTGGTTTGAATACCTGC
TGATTGACCATGATGTCTGCAGCAACTATGGGAACTGGCTGTACAGCGCGGGCATTGGAA
ACGACCCCAGAGAGAACAGGAAGTTCAACATGATCAAGCAGGGCCTGGACTACGACAACA
ATGGTGACTATGTGAGGCAGTGGGTGCCTGAGCTGCAGGGGATCAGGGGAAGTGATGTAC
ACACACCCTGGACCCTCAGCACTGCGTCGCTGTCACACGCCAACGTCTCCCTCGGTGAAA
CCTACCCAACCCCCATCGTCATGGCGCCTGAATGGAGCCGACACGCTAACAAGAAACCGA
GTGGCGCTGGGCCTTCATCACGAGGAAAGAAAGGCCCGTCTCACACTCCCAAACAACACC
GTGACAGAGGAATAGATTTCTATTTCTCCAAAAGCAAAAACCTTTGACCCTCACAGTGAA
CCCAGGAGGGAAACTCCAGTGATTCTAACTTGGACTGCTGCTGGACAAAATCGGTGTTGC
CTGATGAGGATTTTATCAAATCACAAAGCGCCTTTTATGGATGTGCTTCCTGTTCGACTG
CAGTAACGATGAAGCAAAACTGTGGTTTTCATGCAGTCAGATGATGCTTTGATGCAAGTT
TTAATGTGGTTTTCTTTTGAAGTTAGCTTTTTTATGTGTGTTACCCAATAAAATGGAAAT
ATTAAGTTG

>emac_fin_isoseq_HQ_transcript/2370_csnk1da mRNA_complete cds
GAGCTACAACAGATCACTGACAGAGACATACAGGATCCTGCGGAACCCTCTCAAAATCCA
TCTTCATCCGTACTGAATTAAAGCGTAGCCATCTCAAAAACAGCAGGAGCATAAAGTACA
CACTTCTACCTTTCTAACGAGTGCTTTATTAGCACGGCATGACGTTATTAGGATTTTAAA
ACAAGAGGCGAAAGATTGTGAGAGCTGACTTGACTATCCGGAGAGGACATACTACAGAAC
AGAGGACACTGTTCATCTTATCTGAAAGGAGGTTTCATGTTTCGGATGCTGAACAGGCTG
```

GGGCTGTGAACCTGAACGTTATTTTGGTGAAATTTGGGAGCTAGCGAGCTGCTTGCTAGC
CGAGACCATGGAGCTGAGAGTGGGGAACAGATACAGACTGGGCAGGAAGATTGGAAGTGG
ATCATTTGGGGACATCTATCTGGGAACTGATATCTCCGTGGGAGAGGAGGTCGCCATCAA
GCTGGAATGTGTGAAGACCAAACACCCCCAGCTCCACATAGAGAGCAAAATCTACAAGAT
GATGCAGGGTGGAGTGGGCATCCCGACGATAAAGTGGTGCGGGGCGGAGGGGGACTATAA
TGTGATGGTGATGGAGCTGCTGGGGCCCAGCCTGGAGGATCTGTTCAACTTCTGCTCCCG
GAAGTTCAGCCTCAAGACAGTCCTGCTGCTGGCTGACCAGATGATCAGCCGCATCGAATA
CATCCACTCCAAGAACTTCATCCACCGAGACGTGAAGCCGGACAACTTCCTGATGGGGCT
GGGCAAGAAGGGCAACCTGGTCTACATCATCGACTTCGGCCTGGCCAAGAAGTACCGCGA
CGCGCGCACACACCAGCACATCCCCTACCGCGAGAACAAGAACCTGACCGGCACCGCCCG
CTACGCCTCCATCAACACTCACCTGGGCATTGAACAGTCCAGGCGGGACGACCTGGAGTC
CCTGGGCTATGTCCTCATGTACTTCAACCTGGGCTCTCTGCCCTGGCAGGGCCTCAAGGC
TGCCACCAAGAGGCAGAAGTATGAACGCATCAGTGAGAAGAAGATGTCCACGCCCATCGA
GGTCCTCTGCAAAGGATACCCATCGGAGTTCGCCACCTACCTGAATTTCTGCCGCTCGTT
GCGCTTCGACGACAAGCCCGACTACTCGTACCTCCGCCAGCTCTTCAGGAACCTGTTCCA
TCGACAGGGCTTCTCCTACGACTACGTCTTCGACTGGAACATGCTCAAATTTGGTGCCAA
CCGAGCCGTGGAGGATGCAGAGAGGGAGCGGCGGGAGCGTGAGGAGCGTCTGAGGCACAG
CAGGAACCCCGGGGCCCGGGGCATTCCCTCGGCCTCAGGCAGAGCCAGGGCAACACAGGA
AGTCGCAGCCCCCTCCCCACTCAACCCCGCCTCACACACAGGTTTAGAGAAGGAGAGGAA
GGTGAGCATGCGTCTTCATCGCGGAGCGCCAGTCAACATCTCCTCTTCAGACCTGACGGG
ACGCCAGGACACGTCCCGCATGTCCTCACAGGCTCTGTCCTCCCGAGTCACACCCAGCGG
CCTCCAGTCTGCAGCTCCACGGTGAAAGCTGCCAAAAACATCCACCTGTGCACCATGGAG
GCCTCAACATGCACCGCCTGCCTGCAGCACACACACACAAACACACAAACACACACATAC
ACTTTCATGTATACACACAGGACTGTTAAATGACAGAGGGGGACTACCATCCAATGCACC
AGAAGAGTCTGGCTGCTCTCTTGTTCACTTTACAGAGGAAGAGCTGCACAACCAGGAGTG
TCTGGTGTGTGTCTGTATGTGAGCACCTCCCCCCCTCCCTGAAGCCTTGGATCTTCTTTT
TTTAACCTTTTTGGTTTTACTTTATTTTTTGTTCTTTAGCATTAAAACCTTTCTTCCACC
ACTGATCAGACTTCACCAGGGTCTGTTGTCGTAGGACTTTTTGGAAAACTGGAAGAGACA
CGATCCATCTTTGCATTGTTAAAGGACAGTGCCTAAACTCTAATGTTTGTCCTGAACGAC
TCCTTGAATCCCACAAGTCTGTTTTTGTTTAGATGTTTGAGGTTTAAATGTTAAGGTAGG
CATTGTTTTAAAAATGGGACAAGAGGTCATTTTAATGTTCTAGTTTCAAAGGTGTCTTTT
CTTTTTTTTTTGTCCTCCTCAGGTGCGTGTTGTATTTGATCCGAAAACACGCAGCACACT
CTCTGTCCGTGTACAGGCTGTGACTGCAGTGTGCTGCTTGGGAAATCTGTGAATGGATGA
CATGTTGTTTTTAATCTGTGCGTAGTGATGTGCTTTGTGTGGTTCTGTCTTAGGGTTATA
CAAGAACATGTCAGGGTGTATTTTCTTTTAGGGTTCCTTACTCCTCAGCCCCTTAACTGC
TTTTTACACCACATAACATCTGCCTCCATTTTTTTCAAGTCCACCTCTGGCCATTTTCTA
CCGAGACCTCGACAAACTCTGTATTATCGGTGGACTCAGCATTTCAGGAAGTTACAACTT
CCTCCTTCTTTGCAAAAAAAAAACAACCTCACAGATGCCAACAAAACTGTCTTTTCCAAT
GGAATGACTGTTCTTCAGCGCAGATAAATCCACGGCCGTTCTCACCGCGGTCACATAGTG
GTGAATATTATTTGCTTGTTTGCTGGATTGTATTTCAGAAATGCATTCACATAAAGGAAG
CATTTCTTCTTTTAAAGCTGCCTTCATATGATGCGCGATGCTTCTATGGTTAGCGGCCCT
GCTAGCTACTGTAACTGGTATTTGATTAGTTGCGCTTCGAAAGGTCAGCAGCGAGTTTGG
TCAGAATAGCATTTATTCTCTTAAAGTTGCCTTCACATTATGCGCAATTTAAACCTCACT
TACTGAAGTTAGGCATGTAGCATTGCAGCGAAAACAGCTAGCTAGTTTTGCTAGTATGGT
AACTTTTTCTGCTAGCTAGCACCGCTAAGTACAGCGGTTATCTGATTGGTTGCGCAACAA
TCAAGCTAAACAACATGTTAAGAATTTGAAGTTTGGGCACAGCAAAACATCACTTCTGCG
AGTTTAGTTGGAATAGCATTTCTCCTCGCCTGTAGCCGCCTTCACATATTCAGCAGTTTT
TGTTTCTATAGTTAGCTCCCCTGCTAGCTTGCGCCGCTACATGCAGCTGGTATGTGATTG
CTCTGCGAGTTTGGTCGATATCAAAGCATTTCTTCTCGCGTAAAGGTGCCTCCTCCTGTT
ACGCGATTAAAGGGTAAGCCTTGCAGGAAAACAGCTAGCTAGCTTCTACGGTAACCTCCC
CTGCTGGCTTGCACCGCTATTAAAAGCTGGTTGAAAGGTAATCGACAATCGAGATCGAAA
GAAATGTCAACTCTGCAATTTCAAGCAGTGGGCGAAGCCAAGGACCGCTCTCTTCATAGA
AACAGCAGAATGTCAACACAGCTGTTTTTTGAGTTTGTGAAGGCAGCTTCATCCAAATAA
TTCCTGTTAGGACTTCGCTACGTGATTGTACAGTACACCACTGGCGTAAAGTATACCCCC
GCAGCCCCCGCGAGGCGGGGGGGCCCCGGCCCCCCTCATATGAGAACGGGCCCCCTACCT
CTATACCCCAAAGCGGCCGCCGACAGCATGTTAGGTTTCTACTAGAGCGGCCGCGGCCGG
CCGCAATTTTGAAAGACACTACATGAAATAATACTGCGCCAGTGGATACATGAACAACTT
ACTTTTATTTTGAAACATTATTTTGAGTAGGCTAGTAGTAAGGAGGCCGTGGCAGGTGGC
GGTGGGGGGCCCCAGTCTAAGTACCTGCAGGGGGGCCCCAAGTGCTTGTGTTACGCCAGT
GCAGTACACAACAGAAGTGAAGCCATGCAGCTCACACGATTCATTTTTGACATGAAACAA
AAATAAAAGGCACTCGTTTGGATCAGTAAATGTGAGAGTCCAGGTAGAAAATGGACCGCC
ATGCTTTTACGAGGCTTTACGGACGTAAGGAGGTCATGGAGAGCTGTCTTAAGAAAGCGT
GTACGAGCCATGTGGGACTTTTTCGGTCACCTGTGAATGACCAATGAATCCTGCAGACTT
TAAAGTAGAAGAATCCTTGATGTGATAGACTGGTTGTAGGCTGTAATCGTGGGAGGGTGG
GATGGGGGGGTTGTACATGCTTGTTCTGTTGTTGTGTATAAGGTGTTACTGTCTGCCACG
TCCGGATGATGGAGGCGGAATACGGAGGGATTGGAAAATGCCTCATTACACACCTCACAT
GGGGGCTTGAAAAACAATCTCAGAAAGTGGCCAATGAGCTTGTCTCACTGACGCTGGCGA
TGATAGATACAGGGACACTTAATAAGGGATAATAGTTTTTTCCATGTCTGCTTCTCACTT
CTTTTCTTAGTTTGTCACCTCCGGTCCAGTTTTGCTTTGAACTGCGCAGCGGATTTATCT

CTTACTGTGGATTCAAGGGAAATCTTTCTTTCCACTCCCACCTTAAGACTTATGCTCCTC
ATTTGTAATGCCACTGTATTTGTGTATTTTAAAAGTGTAAATAAATAAGTACTTGC

>emac_fin_isoseq_HQ_transcript/50994_csnk1da mRNA_complete cds
ATCCGGAGAGGACATACTACAGAACAGAGGACACTGTTCATCTTATCTGAAAGGAGGTTT
CATGTTTCGGATGCTGAACAGGCTGGGGCTGTGAACCTGAACGTTATTTTGGTGAAATTT
GGGAGCTAGCGAGCTGCTTGCTAGCCGAGACCATGGAGCTGAGAGTGGGGAACAGATACA
GACTGGGCAGGAAGATTGGAAGTGGATCATTTGGGGACATCTATCTGGGAACTGATATCT
CCGTGGGAGAGGAGGTCGCCATCAAGCTGGAATGTGTGAAGACCAAACACCCCCAGCTCC
ACATAGAGAGCAAAATCTACAAGATGATGCAGGGTGGAGTGGGCATCCCGACGATAAAGT
GGTGCGGGGCGGAGGGGGACTATAATGTGATGGTGATGGAGCTGCTGGGGCCCAGCCTGG
AGGATCTGTTCAACTTCTGCTCCCGGAAGTTCAGCCTCAAGACAGTCCTGCTGCTGGCTG
ACCAGATGATCAGCCGCATCGAATACATCCACTCCAAGAACTTCATCCACCGAGACGTGA
AGCCGGACAACTTCCTGATGGGGCTGGGCAAGAAGGGCAACCTGGTCTACATCATCGACT
TCGGCCTGGCCAAGAAGTACCGCGACGCGCGCACACACCAGCACATCCCCTACCGCGAGA
ACAAGAACCTGACCGGCACCGCCCGCTACGCCTCCATCAACACTCACCTGGGCATTGAAC
AGTCCAGGCGGGACGACCTGGAGTCCCTGGGCTATGTCCTCATGTACTTCAACCTGGGCT
CTCTGCCCTGGCAGGGCCTCAAGGCTGCCACCAAGAGGCAGAAGTACGAACGCATCAGTG
AGAAGAAGATGTCCACGCCCATCGAGGTCCTCTGCAAAGGATACCCATCGGAGTTCGCCA
CCTACCTGAATTTCTGCCGCTCGTTGCGCTTCGACGACAAGCCCGACTACTCGTACCTCC
GCCAGCTCTTCAGGAACCTGTTCCATCGACAGGGCTTCTCCTACGACTACGTCTTCGACT
GGAACATGCTCAAATTTGGTGCCAACCGAGCCGTGGAGGATGCAGAGAGGGAGCGGCGGG
AGCGTGAGGAGCGTCTGAGGCACAGCAGGAACCCCGGGGCCCGGGGCATTCCCTCGGCCT
CAGGCAGAGCCAGGGCAACACAGGAAGTCGCAGCCCCCTCCCCACTCAACCCCGCCTCAC
ACACAGGTTTAGAGAAGGAGAGGAAGGTGAGCATGCGTCTTCATCGCGGAGCGCCAGTCA
ACATCTCCTCTTCAGACCTGACGGGACGCCAGGACACGTCCCGCATGTCCTCACAGGCTC
TGTCCTCCCGAGTCACACCCAGCGGCCTCCAGTCTGCAGCTCCACGGTGAAAGCTGCCAA
AAACATCCACCTGTGCACCATGGAGGCCTCAACATGCACCGCCTGCCTGCAGCACACACA
CACAAACACACAAACACACACATACACTTTCATGTATACACACAGGACTGTTAAATGACA
GAGGGGGACTACCATCCAATGCACCAGAAGAGTCTGGCTGCTCTCTTGTTCACTTTACAG
AGGAAGAGCTGCACAACCAGGAGTGTCTGGTGTGTGTCTGTATGTGAGCACCTCCCCCCC
TCCCTGAAGCCTTGGATCTTCTTTTTTTTAACCTTTTTGGTTTTACTTTATTTTTTGTTC
TTTAGCATTAAAACCTTTCTTCCACCACTGATCAGACTTCACCAGGGTCTGTTGTCGTAG
GACTTTTTGGAAAACTGGAAGAGACACGATCCATCTTTGCATTGTTAAAGGACAGTGCCT
AAACTCTAATGTTTGTCCTGAACGACTCCTTGAATCCCACAAGTCTGTTTTTGTTTAGAT
GTTTGAGGTTTAAATGTTAAGGTAGGCATTGTTTTAAAAATGGGACAAGAGGTCATTTTA
ATGTTCTAGTTTCAAAGGTGTCTTTTCTTTTTTTTTTGTCCTCCTCAGGTGCGTGTTGTA
TTTGATCCGAAAACACGCAGCACACTCTCTGTCCGTGTACAGGCTGTGACTGCAGTGTGC
TGCTTGGGAAATCTGTGAATGGATGACATGTTGTTTTTAATCTGTGCGTAGTGATGTGCT
TTGTGTGGTTCTGTCTTAGGGTTATACAAGAACATGTCAGGGTGTATTTTCTTTTAGGGT
TCCTTACTCCTCAGCCCCTTAACTGCTTTTTACACCACATAACATCTGCCTCCATTTTTT
TCAAGTCCACCTCTGGCCATTTTCTACCGAGACCTCGACAAACTCTGTATTATCGGTGGA
CTCAGCATTTCAGGAAGTTACAACTTCCTCCTTCTTTGC

>emac_fin_isoseq_HQ_transcript/99369_csnk1db mRNA_complete cds
ACAACGAGATGGAGCTGAGAGTAGGGAACCGATACAGACTGGGCAGGAAAATCGGAAGTG
GATCTTTCGGGGACATCTATTTAGGCACAGATATTTCCGTGGGTGAAGAGGTGGCCATTA
AGTTGGAATGCGTGAAGACCAAACACCCCCAGCTCCACATTGAGAGCAAGATCTACAAGA
TGATGCAAGGAGGAGTGGGCATTCCAACCATCAAGTGGTGTGGAGCAGAAGGCGACTACA
ACGTGATGGTGATGGAGCTGCTGGGGCCCAGTCTGGAGGATCTCTTCAACTTCTGCTCTC
GCAAGTTCAGCCTGAAGACGGTCCTGCTGCTCGCTGATCAGATGATCAGTCGTATTGAGT
ACATTCACTCCAAGAACTTCATCCACAGAGATGTGAAGCCCGATAACTTCCTGATGGGTC
TGGGCAAAAAGGGCAACCTGGTCTACATCATCGACTTTGGCCTGGCTAAAAAGTACCGCG
ACGCCAGAACACACCAGCACATCCCCTACCGAGAGAACAAGAACCTGACTGGCACCGCCC
GCTACGCCTCCATCAACACACATCTGGGGATCGAACAGTCGAGACGTGACGACCTGGAGT
CTTTGGGCTATGTTCTCATGTATTTTAACCTCGGCTCTTTGCCCTGGCAAGGCCTCAAGG
CTGCTACCAAGAGGCAGAAGTACGAGCGGATCAGTGAGAAGAAAATGTCCACCCCCATTG
AGGTTCTTTGCAAGGGATACCCCTCTGAGTTTGCGACCTACCTGAACTTCTGTCGCTCCC
TGCGCTTTGACGACAAGCCGGACTACTCGTACCTGCGGCAGCTCTTCAGGAACCTGTTCC
ACAGACAGGGCTTCTCTTACGACTATGTGTTCGACTGGAACATGCTCAAGTTTGGAGCTA
ACCGTGCAGCTGAGGAAGCAGAGAGGGAGAGGCGGGACCGGGAGGACAGGCTGAGGCACG
GCAGGAACCCAGGGGCCAGAGGAGTCCCTGCTGCATCAGGACGGCCCCGAGCAGCCCAAG
AGGGAGCCCCACCCACCCCGTTAACACCCACCTCACACACAGCCAACACGTCCCCTCGAC
AAGTATCCGGTATGGAGCGTGAGCGAAAGGTCAGCATGCGACTGCACCGCGGCGCCCCTG
TCAACGTGTCCTCCTCAGACCTGACAGGACGGCAGGAGACGTCCCGCATGTCCACCTCAC
AGATGATGTCCGGTGTACCGAACGCTGGTCTCCATCTTCTAGCTCCTCGATGAGCCCTCC
CCACAATGCACCCTGGCGGACCAGGCCCTTTCCACTAAGAAGCAATCTTAACATTCTTAC
TAACTACTATTCCTAACTACAGCCGCCTGCCGGCTCACTGCTCCCTGACTGCTCTCAGTA
CAGCTCTCATTCAGACTCTGGTCCTCTCCCAGGACTAAACAGTGGCCGCACCTCGGGACC

ACTCTTGTTTTTCAGGAGACGTAAAAACAAAACACCACGAGCAGAGTCTGAATGGGCAGC
GCTTTGTCTGAGGCAAACATGGCAGCCGGACTCACTGAAGTGAAGACTCCTTCCTCTGCA
GAAACATCTGTCATTGAGGACCCCACATTTCCACCGTCTGCTACTTCGAACCTGATCAAT
AAAACATCTCTTTTGAGCAGTCAAATGT

>emac_fin_isoseq_HQ_transcript/5959_csnk1db mRNA_complete cds
GTTAAACTGCCAAAGTACTAACGAAGCAACAGCCTTAGCTAACAACGAGATGGAGCTGAG
AGTAGGGAACCGATACAGACTGGGCAGGAAAATCGGAAGTGGATCTTTCGGGGACATCTA
TTTAGGCACAGATATTTCCGTGGGTGAAGAGGTGGCCATTAAGTTGGAATGCGTGAAGAC
CAAACACCCCCAGCTCCACATTGAGAGCAAGATCTACAAGATGATGCAAGGAGGAGTGGG
CATTCCAACCATCAAGTGGTGTGGAGCAGAAGGCGACTACAACGTGATGGTGATGGAGCT
GCTGGGGCCCAGTCTGGAGGATCTCTTCAACTTCTGCTCTCGCAAGTTCAGCCTGAAGAC
GGTCCTGCTGCTCGCTGATCAGATGATCAGTCGTATTGAGTACATTCACTCCAAGAACTT
CATCCACAGAGATGTGAAGCCCGATAACTTCCTGATGGGTCTGGGCAAAAAGGGCAACCT
GGTCTACATCATCGACTTTGGCCTGGCTAAAAAGTACCGCGACGCCAGAACACACCAGCA
CATCCCCTACCGAGAGAACAAGAACCTGACTGGCACCGCCCGTTACGCCTCCATCAACAC
ACATCTGGGGATCGAACAGTCGAGACGTGACGACCTGGAGTCTTTGGGCTATGTTCTCAT
GTATTTTAACCTCGGCTCTTTGCCCTGGCAAGGCCTCAAGGCTGCTACCAAGAGGCAGAA
GTACGAGCGGATCAGTGAGAAGAAAATGTCCACCCCCATTGAGGTTCTTTGCAAGGGATA
CCCCTCTGAGTTTGCGACCTACCTGAACTTCTGTCGCTCCCTGCGCTTTGACGACAAGCC
GGACTACTCGTACCTGCGGCAGCTCTTCAGGAACCTGTTCCACAGACAGGGCTTCTCTTA
CGACTATGTGTTCGACTGGAACATGCTCAAGTTTGGAGCTAACCGTGCAGCTGAGGAAGC
AGAGAGGGAGAGGCGGGACCGGGAGGACAGGCTGAGGCACGGCAGGAACCCAGGGGCCAG
AGGAGTCCCTGCTGCATCAGGACGGCCCCGAGCAGCCCAAGAGGGAGCCCCACCCACCCC
GTTAACACCCACCTCACACACAGCCAACACGTCCCCTCGACAAGTATCCGGTATGGAGCG
TGAGCGAAAGGTCAGCATGCGACTGCACCGCGGCGCCCCTGTCAACGTGTCCTCCTCAGA
CCTGACAGGACGGCAGGAGACGTCCCGCATGTCCACCTCACAGATGATGTCCGGTGTACC
GAACGCTGGTCTCCATCTTCTAGCTCCTCGATGAGCCCTCCCCGCAATGCACCCTGGCGG
ACCAGGCCCTTTCCACTAAGAAGCAATCTTAACATTCTTACTAACTACTATTCCTAACTA
CAGCCGCCTGCCGGCTCACTGCTCCCTGACTGCTCTCAGTACAGCTCTCATTCAGACTCT
GGTCCTCTCCCAGGACTAAACAGTGGCCGCACCTCGGGACCACTCTTGTTTTTCAGGAGA
CGTAAAAACAAAACACCACGAGCAGAGTCTGAATGGGCAGCGCTTTGTCTGAGGCAAACA
TGGCAGCCGGACTCACTGAAGTGAAGACTCCTTCCTCTGCAGAAACATCTGTCATTGAGG
ACCCCACATTTCCACCGTCTGCTACTTCGAACCTGATCAATAGAACATCTCTTTTGAGCA
GTCAAATGTATAAAGATAAAGCAGCAGCTTTTCATTTTCTAAATATATGTATGGGTAGTT
TTTGGGGTGTGATGCTTTTCACTTTTCATTCCCTCCTCCCACTGACCACTTTTTCTGTTT
TTAACTAGGGCCCCCACCACTGATGGGGCTCTCTTCATTAACTATATAGCCGTTTGTATT
AAATAAAAACAAAGTACTTCTTTGCATGTTTTAGAGGTGAGTCTTCAGCGGCAGGCGTGG
ACTCAACTGCTAGTTGAACCCCTGCACCTACCAACCATGTGCTTTGGTGTGCTGGGTGAT
GGAGGTTGTGTATTCACACGTTAGTTGGCAAAAAAAAAAGTTCTGTCTTTTCATCTCCAG
GCTCTGGGGCCCACTGAGCCCGCTTCCCCTTTCTAGAGTTCCATACTAAGGAAGTGCACT
TTATGTAGGGGGCAAGGACATCAGCTAGCATATTTTACCAGGTGTCCCTTCACTGTGTAC
AGAGAAACATCTTTCTTATTTGTATCATTTTCAAGCTGTGAATTACTTGTAATATTGTTG
TAAAAATGTTTGCTCATCTGTTGATGGGTGTGGCTTGCTCGAGGAACCTGTGAGTTTGAG
ATGCCATAAATGTAGTAGTGTCCCATCAGCCCCCCCGCGGAGAAATGGAGGTATCACAGA
GCCAGCTGCTGGCAGGGAGGGTGGATGTGAAAGACTTGGTGCGACCTGTTTGCTGTCACT
GCTCTTCGGTGATCTGGCATGTGACAAACTGCAGTACTGCTACAGGCACTATAGTTTTGT
GAAACAATTCTTTGTTGAGAGGAAACCTCAACTTGTACAGCAGCAGTGGGGTTTTTTGTT
TTCAATCTGGACACTTCTCTGCCATGTCTATCAGAATGAAATGGCCTGTTCATTCTTCCT
GCTGCTGCTCCTTCTAATATCGGATTTTGACCCTTGACCTCTTCGGTCAGCCGCTTTACA
TCGAGTAAATTATGTAAGCAAACGGATCAGCGAGATTTACTTAATCACATTTATATTTGC
CACTGTATTTGTGTACTTAAACTGTAAATAAATGAATTACTTGTACATAGTTCCTGCGCA
TGAATAAGTTCATCTGTGACCAGAGGGTAGACGCATGTGGTTCATCCACAGAAAATATAT
ATTTATTTTGGGAAAATCAGGCATCACAGGCCTAGATTGTGTTGGACAACACTCAGAATA
GCAATGTGAGGTGCACTGCATTTCTTCAACACGTACACAGTAAAGCACATGGATCACAGC
ATCAGACATCAAACCAACACAACCCAGTGACCCTTGCAGCATCAACAGTAGTAAAGGCAT
ACAGGGACGTACATTCATCAGCTGAGTATACAACACCGACTGACCATTCAAGTTTCCAGC
ACACTGTATCAGGCAGCAACTGAAGAAACCTCAGTAGTAGCTCACAGCTGTCACGTTACA
ACAAAACAAAGAAGACTAACATGGCAGCAACTAGTGAACAGCTACGGTATGACATTTAGC
TGCCCCTTGTTTAAATAGTTTTACAAGCAACTTTGAAATGAGAGTTGCTGTCCTGTTGTA
ACATAAAACCACAAATACATTCAGACATTGTTGATAGTAGCTTGACAATATATTTGTAAC
ACGACGTCCACAAAGTCTTTGGAAGCTTTAAACCGTGTTTCAGTTGTTGTTCTTGAATGA
CATGTTCTAAGTTATCTTGTGAGATAAGCGGCAACTGGTGTTTGTGTATAGGCAGTTTTT
GTATATGTAATGATGATGCAGAACGGAGAGGAAACGTAATTAGTGCTGTCGTTCAATTGA
AACATTTAAATCGAGTTAATCACAATGATAATCTGATTAATCATGATTCAGGCGTAGGAG
AAGGCCTTTTCCCTGCCGGGATTGTAAACGTTTGAGCTGGTATTAGCGACTTTGGGTCAA
ACCTTCTCATGTATTAATTTGCATTAAATGTTTTGTTTTTAAATGTTCAATCCCGAGCGT
AAACTTTGACTGGCCTTTTTTTCTTTCACATGTATGCATTGTAAATGTTACGGGACGTTT
GTACATACATCCGCGGAAACATTTAAGAGACCACTTCAGCATTATCATTGTTCTTCTTTT

ATTAACATTTTTTATTGTATTCTATAAACTACTGACAACATTTCTCTGTGTTGAAATTCA
ACAGACACTGGACTGGCTGCTATACATGTAGAGATAAAGATTT

>emac_fin_isoseq_HQ_transcript/23750_csnk1db mRNA_complete cds
ACGGAGAACATAATCTACGTTTTTCACATTATCGAATGCAGTCGATATTTTAGGTGTTCG
TTCCTCAGAAAGGTAAAAAAGGGATGGTGACATTTGTATTGTTACGGAAGGAGCTGGGAT
TCTGGGGGGCGGGGGAATGTGAAATAAGGGGGTTGTATATCGCTTAGTGAAGCTGTTATC
TGCCTTGGTAAGGTAAACAAAAGGTTAAAAGCTAAACATCAAGGATACGCAACGTTAAAC
TGCCAAAGTACTAACGAAGCAACAGCCTTAGCTAACAACGAGATGGAGCTGAGAGTAGGG
AACCGATACAGACTGGGCAGGAAAATCGGAAGTGGATCTTTCGGGGACATCTATTTAGGC
ACAGATATTTCCGTGGGTGAAGAGGTGGCCATTAAGTTGGAATGCGTGAAGACCAAACAC
CCCCAGCTCCACATTGAGAGCAAGATCTACAAGATGATGCAAGGAGGAGTGGGCATTCCA
ACCATCAAGTGGTGTGGAGCAGAAGGCGACTACAACGTGATGGTGATGGAGCTGCTGGGG
CCCAGTCTGGAGGATCTCTTCAACTTCTGCTCTCGCAAGTTCAGCCTGAAGACGGTCCTG
CTGCTCGCTGATCAGATGATCAGTCGTATTGAGTACATTCACTCCAAGAACTTCATCCAC
AGAGATGTGAAGCCCGATAACTTCCTGATGGGTCTGGGCAAAAAGGGCAACCTGGTCTAC
ATCATCGACTTTGGCCTGGCTAAAAAGTACCGCGACGCCAGAACACACCAGCACATCCCC
TACCGAGAGAACAAGAACCTGACTGGCACCGCCCGCTACGCCTCCATCAACACACATCTG
GGGATCGAACAGTCGAGACGTGACGACCTGGAGTCTTTGGGCTATGTTCTCATGTATTTT
AACCTCGGCTCTTTGCCCTGGCAAGGCCTCAAGGCTGCTACCAAGAGGCAGAAGTACGAG
CGGATCAGTGAGAAGAAAATGTCCACCCCCATTGAGGTTCTTTGCAAGGGATACCCCTCT
GAGTTTGCGACCTACCTGAACTTCTGTCGCTCCCTGCGCTTTGACGACAAGCCGGACTAC
TCGTACCTGCGGCAGCTCTTCAGGAACCTGTTCCACAGACAGGGCTTCTCTTACGACTAT
GTGTTCGACTGGAACATGCTCAAGTTTGGAGCTAACCGTGCAGCTGAGGAAGCAGAGAGG
GAGAGGCGGGACCGGGAGGACAGGCTGAGGCACGGCAGGAACCCAGGGGCCAGAGGAGTC
CCTGCTGCATCAGGACGGCCCCGAGCAGCCCAAGAGGGAGCCCCACCCACCCCGTTAACA
CCCACCTCACACACAGCCAACACGTCCCCTCGACAAGTATCCGGTATGGAGCGTGAGCGA
AAGGTCAGCATGCGACTGCACCGCGGCGCCCCTGTCAACGTGTCCTCCTCAGACCTGACA
GGACGGCAGGAGACGTCCCGCATGTCCACCTCACAGATGATGTCCGGTGTACCGAACGCT
GGTCTCCATCTTCTAGCTCCTCGATGAGCCCTCCCCACAATGCACCCTGGCGGACCAGGC
CCTTTCCACTAAGAAGCAATCTTAACATTCTTACTAACTACTATTCCTAACTACAGCCGC
CTGCCGGCTCACTGCTCCCTGACTGCTCTCAGTACAGCTCTCATTCAGACTCTGGTCCTC
TCCCAGGACTAAACAGTGGCCGCACCTCGGGACCACTCTTGTTTTTCAGGAGACGTAAAA
ACAAAACACCACGAGCAGAGTCTGAATGGGCAGCGCTTTGTCTGAGGCAAACATGGCAGC
CGGACTCACTGAAGTGAAGACTCCTTCCTCTGCAGAAACATCTGTCATTGAGGACCCCAC
ATTTCCACCGTCTGCTACTTCGAACCTGATCAATAAAACATCTCTTTTGAGCAGTCAAAT
GTATAAAGATAAAGCAGCAGCTTTTCATTTTCTAAATATATGTATGGGTAGTTTTTGGGG
TGTGATGCTTTTCACTTTTCATTCCCTCCTCCCACTGACCACTTTTTCTGTTTTTAACTA
GGGCCCCCACCACTGATGGGGCTCTCTTCATTAACTATATAGCCGTTTGTATTAAATAAA
AACAAAGTACTTCTTTGCATGTTTTAGAGGTGAGTCTTCAGCGGCAGGCGTGGACTCGAC
TGCTAGTTGAACCCCTGCACCTACCAACCATGTGCTTTGGTGTGCTGGGTGATGGAGGTT
GTGTATTCACACGTTAGTTGGCAAAAAAAAAAGTTCTGTCTTTTCATCTCCAGGCTCTGG
GGCCCACTGAGCCCGCTTCCCCTTTCTAGAGTTCCATACTAAGGAAGTGCACTTTACGTA
GGGGGCAAGGACATCAGCTAGCATATTTTACCAGGTGTCCCTTCACTGTGTACAGAGAAA
CATCTTTCTTATTTGTATCATTTTCAAGCTGTGAATTACTTGTAATATTGTTGTAAAAAT
GTTTGCTCATCTGTTGATGGGTGTGGCTTGCTCGAGGAACCTGTGAGTTTGAGATGCCAT
AAATGTAGTAGTGTCCCATCAGCCCCCCCGCGGAGAAATGGAGGTATCACAGAGCCAGCT
GCTGGCAGGGAGGGTGGATGTGAAAGACTTGGTGCGACCTGTTTGCTGTCACTGCTCTTC
GGTGATCTGGCATGTGACAAACTGCAGTACTGCTACAGGCACTATAGTTTTGTGAAACAA
TTCTTTGTTGAGAGGAAACCTCAACTTGTACAGCAGCAGTGGGGTTTTTTGTTTTCAATC
TGGACACTTCTCTGCCATGTCTATCAGAATGAAATGGCCTGTTCATTCTTCCTGCTGCTG
CTCCTTCTAATATCGGATTTTGACCCTTGACCTCTTCGGTCAGCCGCTTTACATCGAGTG
AATTATGTAAGCAAACGGATCAGCGAGATTTACTTAATCACATTTATATTTGCCACTGTA
TTTGTGTACTTAAACTGTAAATAAATGAATTACTTGTACAT

>emac_fin_isoseq_HQ_transcript/15321_csnk1e mRNA_complete cds
GGACTATCAACAGGGAGAGCAACCGACCACAGAGCTCAGAGCTGGGGGTATGAGAGCCGC
CAGAATCCGCTCCCATCCTCCTTTACTCGGGCTTCTGTCCGACTGACAACCATCGGATCA
GAACCTCCACTGAACCAGCTCTCGGACGAACCCAATATCGGTGACAACTCGCAAACATGG
AGCTGAGAGTGGGGAACAAGTACCGGCTCGGGCGAAAGATAGGGAGTGGCTCCTTTGGCG
ACATTTACCTTGGTGCCAACATTGCCACAGGGGAGGAGGTAGCCATCAAGCTGGAATGTG
TGAAGACCAAACACCCACAGCTGCACATCGAGAGCAAGTTCTACAAGATGATGCAAGGAG
GAGTGGGCATCCCGTCGATAAAGTGGTGTGGTGCAGAGGGAGACTACAACGTGATGGTCA
TGGAGCTGCTCGGCCCCAGCCTGGAGGACCTTTTCAACTTCTGCTCCCGGAAGTTCAGCC
TGAAGACCGTCCTGCTTCTGGCTGATCAGATGATCAGTCGCATCGAGTACATCCACTCCA
AGAACTTCATCCACCGGGATGTAAAGCCTGATAACTTCCTGATGGGGCTCGGCAAGAAGG
GTAACCTGGTGTACATCATCGACTTCGGCCTGGCCAAGAAGTACCGCGACGCCCGCACTC
ACCAGCACATCCCCTACAGGGAGAACAAGAACCTGACGGGCACGGCGCGCTACGCCTCTA
TCAACACGCACCTGGGAATCGAGCAGTCCAGACGTGACGACCTGGAGTCTCTGGGCTACG

TCCTCATGTACTTCAACCTGGGTTCCCTCCCCTGGCAGGGCCTCAAGGCCGCCACCAAGA
GACAGAAGTACGAGCGAATCAGTGAGAAGAAAATGTCCACGCCCATCGAGGTTCTCTGCA
AAGGATACCCCTCTGAGTTCTCCACATACCTGAACTTCTGCCGCTCGCTCCGTTTTGACG
ACAAGCCCGACTACTCTTACCTACGACAGCTCTTCAGGAACCTCTTCCACCGCCAGGGCT
TCTCCTACGACTACGTCTTTGACTGGAACATGCTCAAATTTGGTGCCAGCCGAACAGCTG
AGGACGGAGAACGGGAGAGGAGGACGGGAGATGAGAGAGATGAGAGGGTCGCAGGGGTCC
CCAGAGGCTCGGCATCACGGGGCCTCCCACCGGGTCCCCTCCCCGCAGCTGCCAACAGAG
TGAGGAACGGGCCGGAGCAGGCCATCTCTAACCCCGCCTCACGGGTGCAGCAGTCCGGGA
ACACGTCGCCTCGTGCCATTTCTCGCGCAGAGAGGGAAAGGAAGGTGAGCATGCGGCTGC
ACCGCGGAGCGCCCGCCAACGTGTCGTCCTCTGACCTCACAGCCCGCCACGACGCATCCA
GAATCTCCACGTCACAGGTCAGCGTGCCATTCGAGCACTTGGGGAAGTAGTATTCTTTGG
CTCTGCATGCACATGAAACTGACAGGGCTGCAAGGTATATTCATCGCTCCCCTTTATTTT
TTTCCTTTTATTTTTACCTTGTCTTATGAAAAGATGATTGCCAATGGACACATGGATCTT
TGGAAAATGGACAGTAATTTTTTTTTAAAGATCCAGGGAAGTTTGTGACAGTATAACAAT
TTGTTTCTGACGCGTGGACCGCCACCACCACCACCACCACCACCACGATTCAGAAAG
CAAACACTCCTGCACCATTTTTACTCCATGGTTGATCATACCCTGCTCCACTCCTCAAAA
CATTTCCCTCAGTACCTTCTGTCTGACAAATGTGAAGAGCTGTTGGCATCAAAGAAGCAA
AAAGTAATCGGGGACGCGCATGTTTCTCGTCAAAGGTCATCCCAGTGATTGATATAGGTT
TAGTAACGATTGCTTTTTCAGCTCGACACCGCAAAAACGATTCTGTGTATGTCCACATTC
TTATCTTTCGTCAGGATATCTTAAATTGGAGCTCCTGTCTTTGCTTTGTTTAGAGAGGAA
TAACAGCTAAGAGATTATGACATGAGGAGAGGCAGGTGAGAGGTTAGTGGGACACAATGA
GAGGTGACTTTTAAAGAGGATCAGTAAGTGATGTTTTAGCTGTCGTAGAGTAATGTTTCC
GGTGAAGTAGCATATGGTGCTTGCTTGCCCTTCATCCACGTCGTGACAACTCGGCCACCG
ACACCTCCACCACCCCTTGATTTTCCAGTTTTCTTTGTTTTAAATGTGAGCGGGGGTAAG
GTGAAGAATGACCATAGATATTGCAAAGACACACTCCACGTCTTGAGGGAAGCGCCTCAG
GAAATTCTCAGCGTATTTGACCACACAGCTGCCGGTTTTGGCTAACAAACAGTGGCTGCA
ATTTTTATGATACTTCTAAATGTTACCTTTTTTAAAATTATTTTTCTTATTGGAGGTCAC
AGGGGAGTGGGCAGGCTCCGTTAGGGATTACTCTTAAGGGTGGGGGGGGGGGGTTGGACA
TCCAATTATGAATTTTTTTTTTTAATGTTTACTTCACTGTAAATATATTTTTTTATTTTGAT
GTAAAGCCAAACTGTGGTTATCTGTTTTTAAAGTAGAAATGCATACTAATGACATACCTT
GGGGCTGGGGGCAGAATTTTAGATATTTAAAGTACTAAACCGGGGGGGAGGGGCTTGTGT
TGCTTTGCCAGCTGGGTCATTTAATATTTTGTGTGTGTGTGAGACCGAGGGAATCCCTGC
TAAGAATTTTCATTCTGTCTCTGAATGCAGACGTTATCCCACTTCTCTGCTATTTATGTT
AAACTATTTTGTACCCGTCTGTCCTCTTATTTGTGTGTCCTTTCAACTTCCTGTAACTTA
TTAGAAATAGTGTGTGTTCATGTTTTTTTTTGCCTGTCCTTTTCTTTTTTCTGCTTCAAA
GGTTGTCCTCTTATAATACTTTCAATATTCCTTCCTGCAAAAAGTACATACTAATAGTTT
TGATGTCACCTCTTGGATTTGATCCTCCAGCAACCTTAATGATGTCATTCAACAGAAAAC
ATAACTGATGCAATCGCAGGGATGTGTAAATAGCAGTGTGTACTGTGTGTAATTACCTTT
GTACAGGCCACCGTGACCCAGTGGCATCGTAACTGTGGGGTTGGGAATGGCCTAGAAGAG
GATGAGGCCAGGGGGGTTTCACTTTATGTATTGTGTACTGATATCTTGTACAGTTCTTTA
ACACACTTTATTTACAACAGTATTTGTGTATATTTTATGAAGTAATAAAAATGAATAACG
TTGCTCAC

>emac_fin_isoseq_HQ_transcript/21893_nr1d2a mRNA_complete cds
ACACACAACAAGGGGGAGCTCCTCGGACTCAGACGCTTGTTCTCTCTTACAGACTCGGAC
CTGCAGATTATTTCCACAAATAGACTCCTGAAATGAGATTTATCTTTAAACTACTCACGC
ACAAACAATTGTAATATCTTCTTGGGTGTTTTGGGAAAGTGGCAGCAGTCGTGACAGACA
TTTTGCATTATATCTGTTTTCCTATTCCTCGAAACGAGATGCCTGAGGACATGGGGACAG
CCAAGCCTGGAGGAGTCATAGCCTATATTTCTTCCGGATCTGCCTCCAGCCCAGAGTCCT
GTATGAGCACCAGCCCGAGCAGAAGCTATTTGTCCACATCCCAGGCCGTGTCCCGCCCTT
CCCTCCCCAGCAGGGCAGTGGGCATGGTGGTGGATATCTCTCCACCCACAAAAAACCAAC
GTAGCCACGGCATGGAGAAGGCTGGGCGCTCCTCCACATCTGCCAAGAGCAACATTACCA
AAATTAACGGCATGGTGCTGTTGTGCAAGGTGTGCGGTGATGTGGCATCCGGCTTTCACT
ATGGAGTCCATGCCTGCGAGGGCTGCAAGGGTTTCTTCAGGAGAAGCATCCAGCAAAACA
TCCAGTACAAGAAATGTCTTAAGATGGAGAACTGCACCATCATGAGAATCAACAGGAACC
GCTGCCAGCAGTGCCGCTTCAAGAAGTGTCTGTCTGTCGGCATGTCCAGAGATGCTGTCC
GATTTGGCCGCATCCCAAAGAGGGAAAAGCAAAGGATGCTGCTGGAGATGCAGAACGCCA
TGAACAACATGATGAGCAACAACAGCCAGCTGCACAGCATGCTGCACGGCCACCAGAGCA
CGTCTCCCATGGAGGCCATGACCAGCGACGCCGGCTCCTCGCCCTCTTCCTCCTCTACTT
CCGATTCCCTGTCAAGCTCCAACCCCTCCTCCCCGCATTGCCCCCAGGACTCAGAGTCTG
TGGTTTCAATGGACACCAACTCCAGCTCCGCCTCCTCCTGCGCGTCAGACAGCGGCGAGG
ACGAGGCCGTAAATGCTTTGAGCAGGCAGCACCAAGATGCCTTTGCATACAGCCAGCAGA
AGGTGCAGAGACAGGGCCAGGCCCCAACCACGTCGAGCACTGTCGCCCTGGAGGCGGGGT
GTGACGGCCGGGCGGAGGAGCAGCAGCAGCAAGATAGCTGGAGCTGCTGGAACAACAATG
CGGTTGTGGGGGGGTACCAGCAAGGATCTTACAGTAACATCCAACACGTCAGCAACACTG
CTTACAGGGAGGAGAAGGTTTACCAGCAGCAGCCTCAGCAGCTCAACAGCACCCCTAAGT
GTGAGCCATCAGATGACAGCGAAACAGAACATGGGTTTAACACACAAACTGCCTCAAGTG
GTTACAATGGACCAACCAGCTGCGTGAATAATAGAGCATACTTGGTCTGTCCAATGAACA
CATCACCCTACGTGGACCCCCACAAGCCGAGCCAGGAGATCTGGGAGGAGTTCTCCATGG

GTTTCACCCCCGCCGTGCGGGAGGTGGTCGACTTCGCCAAGAGGATCCCGGGCTTCCGCG
ACCTCCCGGAGCACGACCAGGTCGGCCTGTTAAAAGCTGGAACTTTTGAGGTGCTGATGG
TCAGATTTGCCTCCCTGTTCAATGTCGTTGAGCGGACAGTGACCTTTCTGAGCGGAAAGC
GTTACAGCCTTGACACGCTGCGGGCGCTGGGCGCTGGCGAGTTGCTTAACTCCATGTGCG
AGTTCAGCGAGAAGCTGGCTGCGCTGAGGCTTGACTCTGACGAAATGAGCCTCTTCACAG
CCGTGGTGCTCGTCTCAGCCGATCGCTCAGGGATCCAGGACCTGAACTCGGTCGAGGCCC
TGCAGGACAAACTGATCCGGGGTCTGAGGAACTTGGTGATGCAGAACCACGGCGAAGAGT
CGGCCACCACATTCACCAAGCTGCTGCTCAAACTTCCCGAGCTGCGTTCCCTCAACAACA
TGCACTCGGAGGAACTGCTGTCCTTCAAAGTGCACCCGTGAAAGCTTCCTGAGGGGTCAC
ACCTATGCCCTCCACCTAATCCACACACCTCCACCACCACCTCCACCAGGACCTGCCTCT
TACTGCCCCCTCACCCTCCCTGTTCGACCAACCCAGTTGTTTGTTTCTAGAATGTATTGA
AAGTTATTTTTTATTCTATTTTTGTTAAGTCGACCACGTGGGGTCTGAAGAGAGAAACAT
ATAAAAGATGTACTGTATTTATAGAAGAGATAGCTTGTTGCACACAAACACACGCAAGTG
CATGTGTGTCAGGTTCTTAGTTTTGCAAACTTGGAGGCGCTTTTCTATTCTCCAGAGAAC
CAAGAGTACTTAACTGGCTTTCTGAAAAGATATGATAAAAAACCAATTAGTCGATTCCAA
AAGCCATTTAATTGTGTGTGTTCTCTGATGAGCGGGTGAATGTGAGAGACAATATGAACC
TCACATACCGCGTGTTTGATTGGTCCAATATTTGACTGATACTGGCAAGGCAAGCTATCA
GAAAATGTATTGAAGAATGTGTGACGACCGCTGCTAATGCATGGCTGTACCTAATGCAAA
CTATTATTACTTGGATCTTCTCCCCTTGCCTCAGAAAGCATCAGTTCAAAGTTTCAGTGT
ATATAGTGTTTTCTTTTTATTTAATTTTTTACTTGTGTTCGTCAACTGCCTGTACACTGC
CTGGTGATTTCTGCGGGTTGTCCATGTTCTGGAGCAGAAAAACCAATAACATGTAAAGGC
TACATGTAAACGTAGTCACACGCATTTTTGCTGTTGTCCCACTTTCCGGGTTTTTTTTGT
GTGTTGTTTTTAATCAAACTTTGTGTGCATTCAGCATAGCACTGAATTTGATGTGGTTGT
CATATATTTTGTTGAAATATCTGTTTCTTCAACTGGCAACACTGCCATATGTAAATATCT
ATATAAATCAATATATATATATAAATATCTGATGAACATAGAGTATAAATATGATATAGT
ATGAATAAATACAAGTG

>emac_fin_isoseq_HQ_transcript/33623_nr1d2b mRNA_complete cds
GAGACCCGCGGTTTCATTTCCTGCAAGAGTAAAGCAAAAACCAGCTTCCAGGACGACAAA
CTGGCCATTTTGCTCTCGTTTCTAACTCAGATTTCTAATATCGAGTCTGTCACTGCCGCT
GTATTTATTTCTAGTGTTTTATTTGATTGCCTGGTTTAGGTGTGGTTGATTTAAATGTTA
TAGCGAATTTCTCCACCGGGATACATCAGCGACTTCTCAGTTGCTGAACACAAACACAGA
AAACACGGCTTCAGCACTGAGGTAGCACAGGAGGAGGAAAACCCCCCGGAAATCACACAA
AATAAAGGATATGGAGTCGACCAAAGCTGGGGGCGTAATAGCCTACATCAGCTCCACCAC
CTCCAGCCCTGAGTCCTGCCACAGTGACTCCTCCAACGGCAGCTTACCCTCCTCCTCCTC
ACCCTCCTCTTCTTCACCCAGCCGCCTGCAGCTGGCACCCCCCACCCCCCAGATCCTGCC
CGGGACACCGAAGACCGGACGCTCCTCCTCATCCTCCACCTCCACCGCAAAATGTGGCAT
CACAAAAATCAACGGCCTGGTGCTGCTGTGTAAAGTCTGCGGGGACGTGGCCTCTGGGTT
CCACTACGGGGTTCACGCCTGCGAGGGCTGCAAGGGCTTCTTCCGGAGGAGCATCCAGCA
GAACATCCAGTACAAGAAGTGCCTTAAGAACGAGAGCTGCCCCATCATGAGGATCAACAG
GAACCGCTGCCAGCAGTGCCGCTTCAAGAAGTGCCTGCTGGTGGGGATGTCCCGGGACTC
GGTGCGTTTCGGCCGGATCCCAAAGCGGGAGAAGCAGCGCATGCTCCTGGAGATGCAGAG
CGCCATGAACAACATGATGAACAACAGCCAGCTGCAGGGCCAGCTGCACGGGGGCCAGAG
CCAGCTGCTCGGGGGCCCCATGCTGCCTTGCCCTCTCCCGGCCACTGACGCTACCTCAGA
GAACGCAGGCCCTGCCCCCTCCCCGCGCTCCAGCCAATCAGACGTCGGCTCTGACCCCGG
CGTCGCCATGGACACCAGCTCGGCCTCTCCCGCGTCTGACAGCGCGTCGTCGGAGGAAGC
GATTGGATCCGTGACGAGGGCCCACCAGGAGACCTTCATGTACAACCAGGAGCCCAGCAG
CCTGGCCCCCGAGGCCCCGCCCCCCGCCGCCACCCCCCCTCACCTCGGGGAGCGGCAGGA
CGCCTGGAACCACCACAACAACCTGGTGACCATCCAGGACCCCCCCTCCAGCCTGGGCAC
CCAGGGTCAGGAGGAGACGCCCCCCCCCACCCACTGTCCCTTCAGGATGTCCAGAGGCTC
CGCCCCCAGCCACTGCCCCTCCTACGCACACGGGGCCTTCGGAGCTCAGCCCCCTGAGGG
CCCTGTCAACACCGCCTGCAGCGCCCCCCTGTGGAGAGGAGGCAACAGGATGCACCTGGT
GTGTCCGATGAACACCTCCCCTCACGTGGACCCTCACAAGTCGGGCCACGAGGTTTGGGA
GGATTTCTCCCACAGCTTCACCCCTGCCGTCAGGGAGGTGGTGGAGTTCGCCAAGAAGAT
CCCGGGCTTCAGAGACCTGTCGCAGCACGACCAGGTCAGCCTGCTGAAGGCCGGCACCTT
CGAGGTGCTGGTGGTCCGCTTTGCTTCCCTGTTTGACGTGAAGGACCACACCGTCACCTT
CCTGGGGGGGAAAAAGTACAGCGTGGAGACTCTGAGGGCCATGGGGGCCGGAGACCTCCT
GAACTCCATGTTCGACTTCAGCGAGAAGCTGGTCAACCTGGGCCTCAGCGAGGAGGAGAT
GAGCCTCTTCACCGCCGTGGTGCTGGTCTCTGCAGACCGCTCGGGCATCGAGAACGTGAA
CTCGGTGGAGGCGCTGCAGGAGACGCTGATCCGGACGCTGCGCAGCCTCATCACCAGGAA
CCACCCCAACGAGTCGGCCGTGTTCACCAGGCTGCTGCTGAAGCTGCCCGACCTGCGCTC
GCTCAACAACATGCACTCCGAGGAGCTGCTGGCCTTCAAGGTCCATTCCTGAACCTTCCT
CTGATACTTGACGAACTGTGAGCGGACCACCTGCTCCTGCCTACAACCCCCACCCCCCCC
ACCCCCACCTCTCAAAGCTGAAATTTGACATTCAAAGGGATGGGGGGGGGGGGTGGACA
CTGAAATGCTGACGAGTTTAAAAAAGCAGACGACAAGGAGACGTAAAGCTGCTAAACAAC
CCTGAATGTTTTAATCTGAGGGAATTAGGGACGTATCATGAGTCTTATCTGCCATGCGAG
TTGTCGTCCATTGCGTGTCATTATTTGTAAATGATTTGGGCATTGGGATGATTTAAAGTG
CCACGATTGCTTGCTGATTTGTGCTCGTGCGTCACTGAAAGAAAGGACGTATTTAAGACT
GTAAGGTTTATTTCGCTCCCACTTGCTTGAATAAACACGTTCTGTTCTACAAAACTGTAC

AGACTTTAGCTGAGTAGTTGAACTGAATAGGTCGGACTATTCCAGATAAAGTATGATTGT
AACGTCACTGAAGTGATTGTAATTTAAAGGCTGTTCTTGTTTCCAGAGCTTCTGTCGTGG
TGGTTCTGTTCTCCTCTGTGAAAGGACAGCAGTATGTGTAGATTTCTATATGAATATAAA
CAAACTATCAAACATTACTGCATAGATGAATATATCTATATAAATACATTTTATAAATGC

>emac_fin_isoseq_HQ_transcript/42009_nr1d2b mRNA_complete cds
GAAACATAGAAAACACGGCTTCAGCACTGAGGTAGCACAGGAGGAGGAAAACCCCCCGGA
AATCACACAAAATAAAGGATATGGAGTCGACCAAAGCTGGGGGCGTAATAGCCTACATCA
GCTCCACCACCTCCAGCCCTGAGTCCTGCCACAGTGACTCCTCCAACGGCAGCTTACCCT
CCTCCTCCTCACCCTCCTCTTCTTCACCCAGCCGCCTGCAGCTGGCACCCCCCACCCCCC
AGATCCTGCCCGGGACACCGAAGACCGGACGCTCCTCCTCATCCTCCACCTCCACCGCAA
AATGTGGCATCACAAAAATCAACGGCCTGGTGCTGCTGTGTAAAGTCTGCGGGGACGTGG
CCTCTGGGTTCCACTACGGGGTTCACGCCTGCGAGGGCTGCAAGGGCTTCTTCCGGAGGA
GCATCCAGCAGAACATCCAGTACAAGAAGTGCCTTAAGAACGAGAGCTGCCCCATCATGA
GGATCAACAGGAACCGCTGCCAGCAGTGCCGCTTCAAGAAGTGCCTGCTGGTGGGGATGT
CCCGGGACTCGGTGCGTTTCGGCCGGATCCCAAAGCGGGAGAAGCAGCGCATGCTCCTGG
AGATGCAGAGCGCCATGAACAACATGATGAACAACAGCCAGCTGCAGGGCCAGCTGCACG
GGGGCCAGAGCCAGCTGCTCGGGGGCCCCATGCTGCCTTGCCCTCTCCCGGCCACTGACG
CTACCTCAGAGAACGCAGGCCCTGCCCCCTCCCCGCGCTCCAGCCAATCAGACGTCGGCT
CTGACCCCGGCGTCGCCATGGACACCAGCTCGGCCTCTCCCGCGTCTGACAGCGCGTCGT
CGGAGGAAGCGATTGGATCCGTGACGAGGGCCCACCAGGAGACCTTCATGTACAACCAGG
AGCCCAGCAGCCTGGCCCCCGAGGCCCCGCCCCCCGCCGCCACCCCCCCTCACCTCGGGG
AGCGGCAGGACGCCTGGAACCACCACAACAACCTGGTGACCATCCAGGACCCCCCCTCCA
GCCTGGGCACCCAGGGTCAGGAGGAGACGCCCCCCCCCACCCACTGTCCCTTCAGGATGT
CCAGAGGCTCCGCCCCCAGCCACTGCCCCTCCTACGCACACGGGGCCTTCGGAGCTCAGC
CCCCTGAGGGCCCTGTCAACACCGCCTGCAGCGCCCCCCTGTGGAGAGGAGGCAACAGGA
TGCACCTGGTGTGTCCGATGAACACCTCCCCTCACGTGGACCCTCACAAGTCGGGCCACG
AGGTTTGGGAGGATTTCTCCCACAGCTTCACCCCTGCCGTCAGGGAGGTGGTGGAGTTCG
CCAAGAAGATCCCGGGCTTCAGAGACCTGTCGCAGCATGACCAGGTCAGCCTGCTGAAGG
CCGGCACCTTCGAGGTGCTGGTGGTCCGCTTTGCTTCCCTGTTTGACGTGAAGGACCACA
CCGTCACCTTCCTGGGGGGGAAAAAGTACAGCGTGGAGACTCTGAGGGCCATGGGGGCCG
GAGACCTCCTGAACTCCATGTTCGACTTCAGCGAGAAGCTGGTCAACCTGGGCCTCAGCG
AGGAGGAGATGAGCCTCTTCACCGCCGTGGTGCTGGTCTCTGCAGACCGCTCGGGCATCG
AGAACGTGAACTCGGTGGAGGCGCTGCAGGAGACGCTGATCCGGACGCTGCGCAGCCTCA
TCACCAGGAACCACCCCAACGAGTCGGCCGTGTTCACCAGGCTGCTGCTGAAGCTGCCCG
ACCTGCGCTCGCTCAACAACATGCACTCCGAGGAGCTGCTGGCCTTCAAGGTCCATTCCT
GAACCTTCCTCTGATACTTGACGAACTGTGAGCGGACCACCTGCTCCTGCCTACAACCCC
CACCCCCCCCACCCCCACCTCTCAAAGCTGAAATTTGACATTCAAAGGGATGGGGGGGGG
GGGGGGGGGCATACAATGTTCTCGTACTGTACTGTAGGGCGCCACTGCCTTTGTGATGGT
GATGCATACAGACACACACTGATGGACGTGTGTGTTTCCTGTGTTTGTGTGTAGCATCCT
GTGTGTGGGGACAAACATTAGCAGCACTTTCTGTATCGTGTAAGCTGGTATTTCTGCATT
CATAGTGCAATATGTGACTTTTGATTTATAGCGAGGAGGAGAAACTTGTGTATACCTTCG
ACGGGAGATTTAAATGCTGTTTGATAAGCGGGAAAAAAAAACCTGGACACTGAAATGCTG
ACGAGTTTAAAAAAGCAGACGACAAGGAGACGTAAAGCTGCTAAACAACCCTGAATGTTT
TAATCTGAGGGAATTAGGGACGTATCATGAGTCTTATCTGCCATGCGAGTTGTCGTCCAT
TGCGTGTCATTATTTGTAAATGATTTGGGCATTGGGATGATTTAAAGTGCCACGATTGCT
TGCTGATTTGTGCTCGTGCGTCACTGAAAGAAAGGACGTATTTAAGACTGTAAGGTTTAT
TTCGCTCCCACTTGCTTGAATAAACACGTTCTGTTCTACAAAACTG

>emac_fin_isoseq_HQ_transcript/29306_nr1d2b
GATACATCAGCGACTTCTCAGTTGCTGAACACAAACATAGAAAACACGGCTTCAGCACTG
AGGTAGCACAGGAGGAGGAAAACCCCCCGGAAATCACACAAAATAAAGGATATGGAGTCG
ACCAAAGCTGGGGGCGTAATAGCCTACATCAGCTCCACCACCTCCAGCCCTGAGTCCTGC
CACAGTGACTCCTCCAATGGCAGCTTCCCCTCCTCCTCCTCACCCTCCTCTTCTTCACCC
AGCCGCCTGCAGCTGGCACCCCCCACCCCCCAGATCCTGCCCGGGACACCGAAGACCGGA
CGCTCCTCCTCATCCTCCACCTCCACCGCAAAATGTGGCATCACAAAAATCAACGGCCTG
GTGCTGCTGTGTAAAGTCTGCGGGGACGTGGCCTCTGGGTTCCACTACGGGGTTCACGCC
TGCGAGGGCTGCAAGGGCTTCTTCCGGAGGAGCATCCAGCAGAACATCCAGTACAAGAAG
TGCCTTAAGAACGAGAGCTGCCCCATCATGAGGATCAACAGGAACCGCTGCCAGCAGTGC
CGCTTCAAGAAGTGCCTGCTGGTGGGGATGTCCCGGGACTCGGTGCGTTTCGGCCGGATC
CCGAAGCGGGAGAAGCAGCGCATGCTCCTGGAGATGCAGAGCGCCATGAACAACATGATG
AACAACAGCCAGCTGCAGGGCCAGCTGCACGGGGGCCAGAGCCAGCTGCTCGGGGGCCCC
ATGCTGCCTTGCCCTCTCCCGGCCACTGACGCTACCTCAGAGAACGCAGGCCCTGCCCCC
TCCCCGCGCTCCAGCCAATCAGACGTTGGCTCTGACCCCGGCGTCGCCATGGACACCAGC
TCGGCCTCTCCCGCGTCTGACAGCGCGTCGTCGGAGGAAGCGATTGGATCCGTGACGAGG
GCCCACCAGGAGACCTTCATGTACAACCAGGAGCCCAGCAGCCTGGCCCCCGAGGCCCCG
CCCCCCGCCGCCACCCCCCCTCACCTCGGGGAGCGGCAGGACGCCTGGAACCACCACAAC
AACCTGGTGACCATCCAGGACCCCCCCTCCAGCCTGGGCACCCAGGGTCAGGAGGAGACG
CCCCCCCCCACCCACTGTCCCTTCAGGATGTCCAGAGGCTCCGCCCCCAGCCACTGCCCC

```
TCCTACGCACACGGGGCCTTCGGAGCTCAGCCCCCTGAGGGCCCTGTCAACACCGCCTGC
AGCGCCCCCCTGTGGAGAGGAGGCAACAGGATGCACCTGGTGTGTCCGATGAACACCTCC
CCTCACGTGGACCCTCACAAGTCGGGCCACGAGGTTTGGGAGGATTTCTCCCACAGCTTC
ACCCCTGCCGTCAGGGAGGTGGTGGAGTTCGCCAAGAAGATCCCGGGCTTCAGAGACCTG
TCGCAGCACGACCAGGTCAGCCTGCTGAAGGCCGGCACCTTCGAGGTGCTGGTGGTCCGC
TTTGCTTCCCTGTTTGACGTGAAGGACCACACCGTCACCTTCCTGGGGGGGAAAAAGTAC
AGCGTGGAGACTCTGAGGGCCATGGGGGCCGGAGACCTCCTGAACTCCATGTTCGACTTC
AGCGAGAAGCTGGTCAACCTGGGCCTCAGCGAGGAGGAGATGAGCCTCTTCACCGCCGTG
GTGCTGGTCTCTGCAGACCGCTCGGGCATCGAGAACGTGAACTCGGTGGAGGCGCTGCAG
GAGACGCTGATCCGGACGCTGCGCAGCCTCATCACCAGGAACCACCCCAACGAGTCGGCC
GTGTTCACCAGGCTGCTGCTGAAGCTGCCCGACCTGCGCTCGCTCAACAACATGCACTCC
GAGGAGCTGCTGGCCTTCAAGGTCCATTCCTGAACCTTCCTCTGATACTTGACGAACTGT
GAGCGGACCACCTGCTCCTGCCTACAACCCCCACCCCCCCCACCCCCACCTCTCAAAGCT
GAAATTTGACATTCAAAGGGATGGGTGGGGGGGGGGGGGGGCATACAATGTTCTCGTACT
GTACTGTAGGGCGCCACTGCCTTTGTGATGGTGATGCATACAGACACACACTGATGGACG
TGTGTGTTTCCTGTGTTTGTGTGTAGCATCCTGTGTGTGGGGACAAACATTAGCAGCACT
TTCTGTATCGTGTAAGCTGGTATTTCTGCATTCATAGTGCAATATGTGACTTTTGATTTA
TAGCGAGGAGGAGAAACTTGTGTATACCTTCGACGGGAGATTTAAATGCTGTTTGATAAG
CGGGAAAAAAAAACCTGGACACTGAAATGCTGACGAGTTTAAAAAAGCAGACGACAAGGA
GACGTAAAGCTGCTAAACAACCCTGAATGTTTTAATCTGAGGGAATTAGGGACGTATCAT
GAGTCTTATCTGCCATGCGAGTTGTCGTCCATTGCGTGTCATTATTTGTAAATGATTTGG
GCATTGGGATGATTTAAAGTGCCACGATTGCTTGCTGATTTGTGCTCGTGCGTCACTGAA
AGAAAGGACGTATTTAAGACTGTAAGGTTTATTTCGCTCCCACTTGCTTGAATAAACACG
TTCTGTTCTACAAAACTGTACAGACTTTAGCTGAGTAGTTGAACTGAATAGGTCGGACTA
TTCCAGATAAAGTATGATTGTAACGTCACTGAAGTGATTGTAATTTAAAGGCTGTTCTTG
TTTCCAGAGCTTCTGTCGTGGTGGTTCTGTTCTCCTCTGTGAAAGGACAGCAGTATGTGT
AGATTTCTATATGAATATAAACCAACTATCAAACATTACTGCATAGATGAATATATCTAT
ATAAATACATTTTATAAATGC

>emac_fin_isoseq_HQ_transcript/19625_nr1d2b mRNA_complete cds
ACAGTCCCTCTGCACAACCAGGAGCCGAGCTGTTTCTCTGCATTCAGCCTGTTTCTGTCA
ATGGGAAGCGGAATATCTGCATTTAAATCCTGAGAACACAGAGAGAACCAGACCCGCGGT
TTCATTTCCTGCAAGAGTAAAGCAAAAACCAGCTTCCAGGACGACAAACTGGCCATTTTG
CTCTCGTTTCAAACTCAGATTTCTAATATCGAGTCTGTCACTGCCGCTGTATTTATTTCT
AGTGTTTTATTTGATTGCCTGGTTTAGGTGTGGTTGATTTAAATGTTATAGCGAATTTCT
CCACCGGGATACATCAGCGACTTCTCAGTTGCTGAACACAAACACAGAAAACACGGCTTC
AGCACTGAGGTAGCACAGGAGGAGGAAAACCCCCCGGAAATCACACAAAATAAAGGATAT
GGAGTCGACCAAAGCTGGGGGCGTAATAGCCTACATCAGCTCCACCACCTCCAGCCCTGA
GTCCTGCCACAGTGACTCCTCCAATGGCAGCTTCCCCTCCTCCTCCTCACCCTCCTCTTC
TTCACCCAGCCGCCTGCAGCTGGCACCCCCCACCCCCCAGATCCTGCCCGGGACACCGAA
GACCGGACGCTCCTCCTCATCCTCCACCTCCACCGCAAAATGTGGCATCACAAAAATCAA
CGGCCTGGTGCTGCTGTGTAAAGTCTGCGGGGACGTGGCCTCTGGGTTCCACTACGGGGT
TCACGCCTGCGAGGGCTGCAAGGGCTTCTTCCGGAGGAGCATCCAGCAGAACATCCAGTA
CAAGAAGTGCCTTAAGAACGAGAGCTGCCCCATCATGAGGATCAACAGGAACCGCTGCCA
GCAGTGCCGCTTCAAGAAGTGCCTGCTGGTGGGGATGTCCCGGGACTCGGTGCGTTTCGG
CCGGATCCCAAAGCGGGAGAAGCAGCGCATGCTCCTGGAGATGCAGAGCGCCATGAACAA
CATGATGAACAACAGCCAGCTGCAGGGCCAGCTGCACGGGGGCCAGAGCCAGCTGCTCGG
GGGCCCCATGCTGCCTTGCCCTCTCCCGGCCACTGACGCTACCTCAGAGAACGCAGGCCC
TGCCCCCTCCCCGCGCTCCAGCCAATCAGACGTCGGCTCTGACCCCGGCGTCGCCATGGA
CACCAGCTCGGCCTCTCCCGCGTCTGACAGCGCGTCGTCGGAGGAAGCGATTGGATCCGT
GACGAGGGCCCACCAGGAGACCTTCATGTACAACCAGGAGCCCAGCAGCCTGGCCCCCGA
GGCCCCGCCCCCCGCCGCCACCCCCCCTCACCTCGGGGAGCGGCAGGACGCCTGGAACCA
CCACAACAACCTGGTGACCATCCAGGACCCCCCCTCCAGCCTGGGCACCCAGGGTCAGGA
GGAGACGCCCCCCCCCACCCACTGTCCCTTCAGGATGTCCAGAGGCTCCGCCCCCAGCCA
CTGCCCCTCCTACGCACACGGGGCCTTCGGAGCTCAGCCCCCTGAGGGCCCTGTCAACAC
CGCCTGCAGCGCCCCCCTGTGGAGAGGAGGCAACAGGATGCACCTGGTGTGTCCGATGAA
CACCTCCCCTCACGTGGACCCTCACAAGTCGGGCCACGAGGTTTGGGAGGATTTCTCCCA
CAGCTTCACCCCTGCCGTCAGGGAGGTGGTGGAGTTCGCCAAGAAGATCCCGGGCTTCAG
AGACCTGTCGCAGCACGACCAGGTCAGCCTGCTGAAGGCCGGCACCTTCGAGGTGCTGGT
GGTCCGCTTTGCTTCCCTGTTTGACGTGAAGGACCACACCGTCACCTTCCTGGGGGGGAA
AAAGTACAGCGTGGAGACTCTGAGGGCCATGGGGGCCGGAGACCTCCTGAACTCCATGTT
CGACTTCAGCGAGAAGCTGGTCAACCTGGGCCTCAGCGAGGAGGAGATGAGCCTCTTCAC
CGCCGTGGTGCTGGTCTCTGCAGACCGCTCGGGCATCGAGAACGTGAACTCGGTGGAGGC
GCTGCAGGAGACGCTGATCCGGACGCTGCGCAGCCTCATCACCAGGAACCACCCCAACGA
GTCGGCCGTGTTCACCAGGCTGCTGCTGAAGCTGCCCGACCTGCGCTCGCTCAACAACAT
GCACTCCGAGGAGCTGCTGGCCTTCAAGGTCCATTCCTGAACCTTCCTCTGATACTTGAC
GAACTGTGAGCGGACCACCTGCTCCTGCCTACAACCCCCACCCCCCCCACCCCCACCTCT
CAAAGCTGAAATTTGACATTCAAAGGGATGGGTGGGGGGGGGGGGGGGGCATACAATGTT
CTCGTACTGTACTGTAGGGCGCCACTGCCTTTGTGATGGTGATGCATACAGACACACACT
```

GATGGACGTGTGTGTTTCCTGTGTTTGTGTGTAGCATCCTGTGTGTGGGGACAAACATTA
GCAGCACTTTCTGTATCGTGTAAGCTGGTATTTCTGCATTCATAGTGCAATATGTGACTT
TTGATTTATAGCGAGGAGGAGAAACTTGTGTATACCTTCGACGGGAGATTTAAATGCTGT
TTGATAAGCGGGAAAAAAAAAACCTGGACACTGAAATGCTGACGAGTTTAAAAAAGCAGAC
GACAAGGAGACGTAAAGCTGCTAAACAACCCTGAATGTTTTAATCTGAGGGAATTAGGGA
CGTATCATGAGTCTTATCTGCCATGCGAGTTGTCGTCCATTGCGTGTCATTATTTGTAAA
TGATTTGGGCATTGGGATGATTTAAAGTGCCACGATTGCTTGCTGATTTGTGCTCGTGCG
TCACTGAAAGAAAGGACGTATTTAAGACTGTAAGGTTTATTTCGCTCCCACTTGCTTGAA
TAAACACGTTCTGTTCTACAAAACTGTACAGACTTTAGCTGAGTAGTTGAACTGAATAGG
TCGGACTATTCCAGATAAAGTATGATTGTAACGTCACTGAAGTGATTGTAATTTAAAGGC
TGTTCTTGTTTCCAGAGCTTCTGTCGTGGTGGTTCTGTTCTCCTCTGTGAAAGGACAGCA
GTATGTGTAGATTTCTATATGAATATAAACCAACTATCAAACATTACTGCATAGATGAAT
ATATCTATATAAATACATTTTATAAATGC

>emac_fin_isoseq_HQ_transcript/29306_csnk1db mRNA_complete cds
GATACATCAGCGACTTCTCAGTTGCTGAACACAAACATAGAAAACACGGCTTCAGCACTG
AGGTAGCACAGGAGGAGGAAAACCCCCCGGAAATCACACAAAATAAAGGATATGGAGTCG
ACCAAAGCTGGGGGCGTAATAGCCTACATCAGCTCCACCACCTCCAGCCCTGAGTCCTGC
CACAGTGACTCCTCCAATGGCAGCTTCCCCTCCTCCTCCTCACCCTCCTCTTCTTCACCC
AGCCGCCTGCAGCTGGCACCCCCCACCCCCCAGATCCTGCCCGGGACACCGAAGACCGGA
CGCTCCTCCTCATCCTCCACCTCCACCGCAAAATGTGGCATCACAAAAATCAACGGCCTG
GTGCTGCTGTGTAAAGTCTGCGGGGACGTGGCCTCTGGGTTCCACTACGGGGTTCACGCC
TGCGAGGGCTGCAAGGGCTTCTTCCGGAGGAGCATCCAGCAGAACATCCAGTACAAGAAG
TGCCTTAAGAACGAGAGCTGCCCCATCATGAGGATCAACAGGAACCGCTGCCAGCAGTGC
CGCTTCAAGAAGTGCCTGCTGGTGGGGATGTCCCGGGACTCGGTGCGTTTCGGCCGGATC
CCGAAGCGGGAGAAGCAGCGCATGCTCCTGGAGATGCAGAGCGCCATGAACAACATGATG
AACAACAGCCAGCTGCAGGGCCAGCTGCACGGGGGCCAGAGCCAGCTGCTCGGGGGCCCC
ATGCTGCCTTGCCCTCTCCCGGCCACTGACGCTACCTCAGAGAACGCAGGCCCTGCCCCC
TCCCCGCGCTCCAGCCAATCAGACGTTGGCTCTGACCCCGGCGTCGCCATGGACACCAGC
TCGGCCTCTCCCGCGTCTGACAGCGCGTCGTCGGAGGAAGCGATTGGATCCGTGACGAGG
GCCCACCAGGAGACCTTCATGTACAACCAGGAGCCCAGCAGCCTGGCCCCCGAGGCCCCG
CCCCCCGCCGCCACCCCCCCTCACCTCGGGGAGCGGCAGGACGCCTGGAACCACCACAAC
AACCTGGTGACCATCCAGGACCCCCCCTCCAGCCTGGGCACCCAGGGTCAGGAGGAGACG
CCCCCCCCCACCCACTGTCCCTTCAGGATGTCCAGAGGCTCCGCCCCCAGCCACTGCCCC
TCCTACGCACACGGGGCCTTCGGAGCTCAGCCCCCTGAGGGCCCTGTCAACACCGCCTGC
AGCGCCCCCCTGTGGAGAGGAGGCAACAGGATGCACCTGGTGTGTCCGATGAACACCTCC
CCTCACGTGGACCCTCACAAGTCGGGCCACGAGGTTTGGGAGGATTTCTCCCACAGCTTC
ACCCCTGCCGTCAGGGAGGTGGTGGAGTTCGCCAAGAAGATCCCGGGCTTCAGAGACCTG
TCGCAGCACGACCAGGTCAGCCTGCTGAAGGCCGGCACCTTCGAGGTGCTGGTGGTCCGC
TTTGCTTCCCTGTTTGACGTGAAGGACCACACCGTCACCTTCCTGGGGGGGAAAAAGTAC
AGCGTGGAGACTCTGAGGGCCATGGGGGCCGGAGACCTCCTGAACTCCATGTTCGACTTC
AGCGAGAAGCTGGTCAACCTGGGCCTCAGCGAGGAGGAGATGAGCCTCTTCACCGCCGTG
GTGCTGGTCTCTGCAGACCGCTCGGGCATCGAGAACGTGAACTCGGTGGAGGCGCTGCAG
GAGACGCTGATCCGGACGCTGCGCAGCCTCATCACCAGGAACCACCCCAACGAGTCGGCC
GTGTTCACCAGGCTGCTGCTGAAGCTGCCCGACCTGCGCTCGCTCAACAACATGCACTCC
GAGGAGCTGCTGGCCTTCAAGGTCCATTCCTGAACCTTCCTCTGATACTTGACGAACTGT
GAGCGGACCACCTGCTCCTGCCTACAACCCCCACCCCCCCCACCCCCACCTCTCAAAGCT
GAAATTTGACATTCAAAGGGATGGGTGGGGGGGGGGGGGGGCATACAATGTTCTCGTACT
GTACTGTAGGGCGCCACTGCCTTTGTGATGGTGATGCATACAGACACACACTGATGGACG
TGTGTGTTTCCTGTGTTTGTGTGTAGCATCCTGTGTGTGGGGACAAACATTAGCAGCACT
TTCTGTATCGTGTAAGCTGGTATTTCTGCATTCATAGTGCAATATGTGACTTTTGATTTA
TAGCGAGGAGGAGAAACTTGTGTATACCTTCGACGGGAGATTTAAATGCTGTTTGATAAG
CGGGAAAAAAAAAACCTGGACACTGAAATGCTGACGAGTTTAAAAAAGCAGACGACAAGGA
GACGTAAAGCTGCTAAACAACCCTGAATGTTTTAATCTGAGGGAATTAGGGACGTATCAT
GAGTCTTATCTGCCATGCGAGTTGTCGTCCATTGCGTGTCATTATTTGTAAATGATTTGG
GCATTGGGATGATTTAAAGTGCCACGATTGCTTGCTGATTTGTGCTCGTGCGTCACTGAA
AGAAAGGACGTATTTAAGACTGTAAGGTTTATTTCGCTCCCACTTGCTTGAATAAACACG
TTCTGTTCTACAAAACTGTACAGACTTTAGCTGAGTAGTTGAACTGAATAGGTCGGACTA
TTCCAGATAAAGTATGATTGTAACGTCACTGAAGTGATTGTAATTTAAAGGCTGTTCTTG
TTTCCAGAGCTTCTGTCGTGGTGGTTCTGTTCTCCTCTGTGAAAGGACAGCAGTATGTGT
AGATTTCTATATGAATATAAACCAACTATCAAACATTACTGCATAGATGAATATATCTAT
ATAAATACATTTTATAAATGC

>emac_fin_isoseq_HQ_transcript/178_per1b mRNA_complete cds
GGTGTCCCTGCGTGTAAACTACTGTAGGGGGGTCGAGCAATATTTTGAGAGCATATGCAG
ACAGCTGAGCAAAGGCAGTAGTTGCTACAAGGATTTATGCTGTTCCTTATTGACTTGCAA
ACACCTTACCGTGGAATATCAAAGCTGTTGTCGTTTTGAAACATATCTAAGGTGACCTTC
ATCTGAAGCATCACATCTCTGAAGAGGCAAGAAGACGAGATACAGAGTGTTCTCTGCTCT

CACCAAGAGCTGTTTTTCTCACTCTCCCTCACAATCTCTCATCCCATCTGTTGGAGCCCT
TGTGCCAAGGCTGCTCCGTAAACAGAAATTCATCTCTTGTCGGCATCACTTGAGTGACCT
TTCCAGATAATTGCACAGTTTTTGGAAGCCTTCGCTTTGGTCATTTGCAAAATGTTTACA
ACCATTTGAATTTTCTCTGGTTTTTGCACCAGAAACAATGAATTTCTCTGTAAACGACAA
GCCAATAACCATGATTCACCTGTCTATACATAGTAGACCCACTTAAATCAGAGTTATTGC
TGTTTTCACCTTGGCTGACACCTTTAAAAAAGCTGCTCCAGCCTCATCCTACTCTAGTAT
GAGTTATGACAACTCTAATTCATTGCCCAGCAGCAGCACTCGAGGGCGAGTGGCAAGGGC
CGACGAGAAAGAAAATGAACAGGAAGCAGAATCAGAGGAGTTAAACTCTCTGAAAACGAG
CAGTGTTCAGCCCAGCGCTAATGGCCCCGCTCAGGAGCAGGGGAGTGGAAATGGAGGCGA
CTCCACTGGCGGCTCATCTCCCAGTGTGGGGTCGGCGTCCGGAGGCTCGCCCCGAGACCG
CAGGGGGCCCAACTCAGATGATATGGACGGACTCTCCAGCGGAAACGACTCCGGGGAGAG
GGAGAGCGAAGGCGGGATGGAAAGAGACGGCGGGTCACGTGGGCGCCAGTCCGTACGCAG
CTCCCACAGCTCGTCAAACGGAAAGGACTCTGGCATGATGCTGGAAAACACGGAGAGCAA
CAAGAGCTCCAACTCCCAGAGTCTCTCTCCCCCCAGCAGCTCCCTGGCCTACAGCCTGCT
GTCCACCAGCTCCGAGCATGACCCCCCCTCCACCTCGGGCTGCAGCAGTAACCAGTCGGC
GAGGGTGCAGACCCAGAAGGATCTGATGAAGGCCATCAAGGAGCTGAAGCTCCGCCTCCC
AACGGAGCGCAAGGCCAAGGGCCACACCAGCACCCTAAATGCACTCAAATACGCTCTTCA
GTGTGTCAAACAAGTCCGAGCCAACAAGGAGTACTATCACCAGTGGAGCGTAGAGGAGTG
CCAGGGCTGCAGTCTGGACCTGTCTGCCTTCACTATCGACGAGCTTGACAACATCACGTC
AGAATACACACTCAAAAACACAGACACTTTCACCATGGGCGTGTCCTTCCTGTCGGGGAA
GGTCGTGTACGTATCACCTCAGGCCTCATCCCTGTTGCGCACCAAGCCAGAGCGTCTCCA
GGGGACTATGTTCTCTGAGCTTTTGGCCCCGCAGGATGTCAGCACTTTCTACAGCGGTAC
AGCACCCTGCCGCCTGCCAAACTGGGCCTCCTGCATCGGGTCAGCTTCTGCTCCAGTGGA
CTGCACTCAGGAGAAGTCCATGTTCTGCCGGATAAGTGCCGACCGGGCGCAGGGCTGCGA
GATGCGCTACTACCCGTTCCGCCTCACGCCCTACCAGCTCACCATCAGAGACTCGGACGA
CGCTGAGCCACAGCCCTGCTGCCTGCTCATCGCTGAGAGGGTCCACTCCGGATACGAGGC
TCCCCGTATCCCTCCAGACAAGAGGATCTTCACTACAAGTCACACCCCGAGCTGCATCTT
CCAGGAAATTGACGAGAGGGCAGTGCCACTGTTGGGATACCTGCCTCAGGACTTAGTCGG
AACCCCCGTCCTGCTCTACATTCACCCGGAAGATAGGCACATAATGGTGGCTATACATGA
GAAGATCTTTCAGTTTGCAGGGCAGCCGTTTGACTATTCGCCCCTGAGGATGTGTACCCG
AAACGGGGAATATTTGACCATCGACACCAGCTGGTCGTCCTTTGTCAACCCCTGGAGCCG
GAAGGTGGCTTTCATCGTAGGACGCCACAAAGTCCGAACGAGCCCTCTGAATGAAGACAT
GTTCACGGCGCCACAGGGCTGCGAGACTTGGACCACCACGGCCGACGTGGTGCAGCTGAG
CGAGCAGATCCACCGACTCCTGGTGCAGCCGGTGCACAGCGGCAGCTCTCAGGGCTACAG
CTCGCTGGGCTCCAGCGGCTCCCGAGGGTCCCGGCGGTCTCACCAGCAGCACCTCAGCGC
CGCCTCGTCCAGCGACAGCAACGGCCCCGCCACGGACAAAGCGGTCGCTCTGCGCAAACC
TATGACTTTCCAGCAGATCTGTAAAGACGTCCACATGGTCAAGACTAACGGACAGCAGGT
TTTCATCGAGTCTCGTAACAGGCCAACGCCCAGGAAAACCTGCAGCATAGGCACAACAAG
CGTCAGAGCAATCAGCAGCGACCCAATCAGAGGCTTGATAGCCGACATGACGATACCACC
CAAAGCTTTGCTCCCTGCTCCACTCGTGCCAAAGGAGCCTCCTACCGGCTACTCCTACCA
GCAGATCAACTGTCTGGACAGCATCATAAGGTACTTGGACAGCTGTAACATTCCTCACAC
GGTCAAAAGGAAGTGTGGCTCTTCCTCCTACACGTCTGACGAGGACAAGCAGCAAGATGG
CGGCAACAACACAGGTGGTTCAGTTAACCTTGTAAGTGAACCACCTCCTCTGCCTCCCCT
GACCATGGCCACAAAGGCAGAGAGTGTAGCCTCAGTCACGTCGCAGTGTAGCTTCAGCAG
CACCATCGTGCATGTGGGAGACAAGAAGCCTCCTGAGTCAGACATCATCATGGAGGAGGC
TCCCACTACTCCCAATCCTGCTCCTTCTACGTCCGCACCGACTGCTCTGTCGAGCGGTCT
GACGCCTCCCGCTAACACAGCCGTCGCCTTTCCTCCTCTCCCTCCACCTCAAGCCTCTCA
GCTAGAGAGGGACAGCCGGAGGAGCAGCAGTGTAGGAGGGGGCCGTATGGGCCTCACAAA
GGAGGTGCTATCTGCCCACACCCAGCAGGAGGAGCAGGCGTTCCTTGACCGCTTCAAGGA
CCTCAGCAAGCTGCGCGTCTTCGATCAGACGGCGGCATCAACTCTGCACTGTCAGAACCC
GGCTGCCAATCCTCTGTCGCGAGGAGTGCGCTGTTCCCGGGATTACCCCGCCGCGGGCAG
CAGCACCAGCCACCGGCGAGGTCGAGGCGGGAAGAGACTGAAACACCAGGAGTCGGCCGA
GCAGCACAGCTCTCTGCACCTGAGCGGGAGCCGGCATGACCCCCGAACCAGCACAGCCCC
CATGCCTGTCAACATGCCCATCGGACCCCCGACAAACTCCTCCTCCTGGCCTTCTGTAGG
CTCCCAGGCCAGCATTCCCGCCGCACCATTCGCTCCGGGAATGCTTCCCATCTACCCCGT
TTACCCACCGATGGCACAGCCCATACCCGTCCCAGATCCATCCCGTTTCCCCCCTGCCCA
GATGGTTCCTCCCATGATGGCCCTCGTTCTGCCCAACTACATGTTCCCCCAGATGGCAGC
CCCCATCGCTCAGCCGGCCCCCACCACCGGACACTTCTACAACCCTAACTTCACATACCC
CACCCCAGTCGTCCCAGTTATCTCCAACCCAGTGCCGATCCCAGCCACCTGTGCCCCGTC
CCGCAGCAGCACCCCCCAGTCCTACAGCCAGCCCCCGCCGACCGAGAGGGCGCAGAGTC
CCCCCTCTTCCAGTCCAGATGTTCCTCACCGCTCAACCTGTTGCAGCTGGAGGAAACACC
GAGCAACCGGCTGGAGGTCGCCACGGCGCTGGCAGCATCACAGCAAGCCCCGCCCTCTGT
TCAGGGGGGAGCGGCTGGGGGACAGAGCTCAGCCAATCAGAGGAGCTCTGATGAAATATC
CAAGGAGAATGAGAACGGCGAGGCTAATGAGTCCAACAACGATGCCATGTCCACCTCCAG
CGACCTGCTGGATATGCTGTTGCAGGAAGATGCCCGCTCGGGCACCGGCTCAGCCGCCTC
TGGTTCAGGGTCCTCAGGCACGAGGTCCTCCGGCTCAGGCTCTGGATCCGGCTCCAACGG
TTGCAGCTCTTTTGGCACCAGTGGCATGAGCAGCAGCCAGGGCAGCCACACCAGCAAGTA
CTTCGGCAGCATCGACTCATCAGAGAACGATCATTCCCGCAAACAGCCAGCAGGGGGCAG
CAGCAGCGCAGGCGGAGATGGTGCAGAGGAGCAGTTTATCAAGTGTGTCCTGCAGGACCC

CATCTGGCTGCTGATGGCCAACACCGACGACAAGGTCATGATGACCTACGAGCTGCCTGT
CAGAGACATGGAGACTGTGCTACGAGAAGACCGCGAGGCGTTGAGGAACATGCAGAAACA
GCAGCCACGCTTCAGTGAGGACCAGAAGAGGGAGCTGAGCCAGGTTCACCCCTGGATCCG
CACAGGACGCCTGCCCCGAGCCATCAACATCTCTGGCTGTACAGGCTGCAAGTCCCCCCC
CTTCGAGCCTCCCACGGCTCCATTCGATGTTGTGATACACGAGATGGAGATGTGCAGCGC
GCTAAAAGCACGAGAAAATATTGCTGGAAAGGGCAATACAATTCCATCAGAGACAGCCAT
GGAGGAAGCTCACCAAGACGACGAAGAGGAAGCAACCAAAACGCAAGACAGCAACCAAGA
AATGACAACCGAGGCCTCAGAAGCTGTGAAGGAAAGTCTCACATGACTCACTAAGGAGGA
GGCCAAAGACGAATAACGCTATGGAGACATTTCCGCGCTGATGTGCAGAACACGTGCACC
TTCCCAGACTTCTTGTGCCCAAAAATAAATCACTGCACAGAGTTAAGTCGCTGAAGCGGG
AGGAAAAAAAGGTGCATATTGATCGAGACAACTTCCGTTATTGCTGGACAGCACAACACA
ACACTGCATATTGAAATGCCTTGGAGGCAGGACACACGCACTTTCCTCGTTCTTGGCATG
TTTATTGAAGCTGTAAGGATAGACGGGAAGATGAATGTGCATCAGAACGACGCCTTTATG
TAAAAACGGACATGCAATGTCCTCGAAGTTTCGCTCACGAGGGAAAGAAGGGAGGTTTCA
GAGACTTTTCTGTTTTACAGGTCATGGAATCCTAATCCAATCCTGTTTCTCTCAAAGACT
TAATTTTGTCAATTTTTGTATGTATAAAATGTGCCCTGATAGAAGAGAAACCCACGTTCG
CTCTATTGAAAGATATAGAGCGACAGCCAGTTTATTTTGTAAACTGTGTATATTTTTATA
CTTTCTGCCACTGCGAATTGTGATAGGATAAATGTCTTAAAGAGTGTGTACTGTCCCGTA
TCAGTTCAGTCTTTACACAGACATGGTGTTTGTTCTGTTGCTTCACTTGAGGGATGTACA
GTGAATGTGTGGGACCGTGTGGCAATCGCCAGGAGACTGACCAGGACTTTAAGGTCTCTC
CTCGGCCGGGCATTAGTAGGAGCATTGGGGAGGAATCCCCACAGTGAGAAACGGACAATA
TCTCTCTGAGAAGTGATGACAGTATTCGCTGAAAACGGAGCTCTGAATCCTGGAAAGGGA
GGGTTCCTGTGTAGCAAGCGCTTGCACACGGTTGCGAGAAAGAGGTCGGCGACGTTTCTC
TCTGAGAGATTGGAATGTTAAGTATATTTCCTCCTGTCTGTCCCTATCGCTCTACCTGTG
ACACTGTTCTGGTCATGTGATATCCTTTTGCTTCAGTCAGTGGTGTCCGCAACAGTTTCT
AAATATTTTTGTGAAATTCTTTGTACAAAAAAGACAATAAAAACGACTGGAAAATCCTTT

>emac_TRINITY_DN55851_c0_g1_i4_per2 mRNA_partial cds
GAGATCATAGGGGATGTCCAAGGAACAGGAGAAACAGTAGAATCCAGCCAGAGCCCCGGCGCTCCTCTCTGTGTTGTTTCACCTCCCAGTCAAGAGAGGGAGTCCT
ATAAGAAACTCGGGTTGACCAAGCAGGTTTTGGCAGCCCATACGCAGAAGGAGGAGCAGACCTTTCTGTATCGCTTCAAGGAGCATCAAGGACTTACAGCGCTTAA
AGAAAACTGCTCTCAGTACCTGGAGCGTCAGAGGGAACAGATCGCCAGCTATGCTATACCCGCTGCTCAGTCCTCCAAGCAGGATGGACCAATTGCAGAGCCCACC
GCTCGGCGAGGTACACGGAACAAGAGGACCAAGTCAAAGCGGGCGAAGCAGATGGAGTCCTCCGACAGCACAGTGTCCCATCACAGACAGCAACACCTGCGGCCTC
CTCTTCTAAACCACGGCCTTAACCTGACCTCTTGGTCGAGCTCTGACACCTCGCAGTCAACGTTCCCCATGGCTTACCCCTCCGTGATGCCAGGCTACCCCCTCCC
GGTGTACCCCAGAGGCAGTTCTATAGCCCCTCGCACAGACGCCACTCTTCAAGGCTTTGTGGACAATCAGGGTACCCAGCCCCCTCCCTGCCCCCCACCCATCCAC
ACTTCTCCCTACACCGCCCCTATGGTCACTCCTATTGTGGCTCTCATGCTGCCCATGCCCTTCTCTCCATTGGCCCCCTCACTGCCACCTCCACAGCCAATGTACC
ATGCAGCCACTGCTGGCTTCCCCACCCAAATGCAGCCTTTTGGTCAGGCTGCCTTTCCAATCCCAGGCCCCTTCGCAGCGTCTCCATCCTTCACTGTCCAGAACCA
GTTCAACTCACCAAACCACTTTGCTCTACCGTCGAACTACATTCCCCCTTCGTTCTACTTTCCTCCAGTCTCGGAAACCTCAAAGGCACCCATGGTTGAGAGCCAG
TCTCGCTCCTCCACGCCGCAGTCTGGAGGAGGTGGAGGCCCGGCATCTCCTCCCCTGTTTCAGTCTCGCTGCAGCTCACCCCTCAACCTGCTGGAGCTGGAGCTGT
CGGTGGACCGGCAGGACAGCACAGCGTTCTCCTTTGGAGGTCAAGGGAATAATATTGCAGAAAGGGAGAAGGGAGCCAGTGGAAACCAGGCCAAAGAGAGGGAGCT
GAAGCAGCCTTCACTCAGTCTCCTTGGTCCTCCTGGACCCTTGTATTGCGAGGGCTCGCCCTGTGTCTGTGATCGTTTGTCTTGGGATAAAGTGTGCTCTAGAAGG
TCTTGCAGCAAAGAGGCAAGTTCACGTGGCGATGGGAACAACAGTGATGCCAACTCTTTGTCCAGCGACATGTTGGACATTATTCTCCACGAGGACTCAGCCGACT
CGGGGTCCATGGGCTCTGGATCGAATGGTTGTAGTACTTCAGCCAGTGGGACCTCCAATAGCGGGACGTCTAACAGTAGGACATCAGACAGTGGTACATCAAAGAG
CAGGACATCAGCCAGTGGGACCTCCGGCAGTGGGACAGGAAGCAACAACAGTAGTAACTACTTTGGCAGCGTGGACTCGTCGCAGAACAGTCAGAAGGTCAAAGGT
CACCTGAGTGGCAGCGATGGCAGGCCTTTGGAGATGGACCACAGTGAACACTTTATCACTGACATCCAGAGAGTGCTCAGAGACGACAGAGAGAAGCTGAGGATGC
TGCAGAAGAGCCAGCCGAGCTTTTCAGAGGAGCAGAAGAAGGAGCTGATGGAGGTGCACACCTGGATTAAGAGAGGAGGTCTGCCAAAGGAGATAGACATCAAGGC
GTGCTCCTGTTGTAACAGCGTCTCGGAGGCAGCAGCGGTGGAGGAGGAACAGGCAAACCTGGACATCGGTGAAACGGAGACACTGGAGGAGGGCTGTCAGCGGAGG
CCCAGAGAGGAACCCTCAACCTCAAACTCTCTCTCAGATACTACCAGCTAGACAACAAACTCTTCAAGATCTTCAAAGTGTCATCAATTACCCGTCAGACTCTGTT
TCTGCACAACTGTGAAAAGTGGCCTTCTTTGCGTGAAGCACTTAACTCTTCTCACAACATTCATTGCCGTCTTAAGTCAAAACGTTTGTTGAATGTAAATATATGA
GTGCTGGATTCTGTGACTGCATATGTTCATTCACACATTCTGTTTAGCACTTAATATAGTGCAGCATCTGCATTGATAGGTCATAAGTGAAACGCAATGTGTGCCC
ATGCACCGAGCAGCACAGTAGATGTCGAACACACACAAACAAACCAGAGGCTGCATCATTCACATGGCAGGAAGTGGCTCTTTGGTGTTCAAACTGCTGCGGTAAA
ATAGTTGATTCAGCTTTTTGTGTCGTGATTTCCTCAGTTACAACAACAGCACGCTGATTCAGCTTGTGGTAAAAACAGGCGGAGTTCAATTTCAGTAACTCCTCTT
CAGTGTTGTATAGCCATCACGCACTAAATGAATGCACATTTGCACTTTGTGAGCTACACCAGATAAAAGATTAAATGTGTTGTGTTTAGTCACTAGTAGCAAATTA
TGTCCAAAACACAAACTTGCTCCCGAATGCAGCAGGCATGGACTCACAATCTGAAGCTTTTCACCACCTCCATCCTGAAAACATACTCCCATACCATGCAATACTA
AACGTACAATATTATATATTGAACTTAAGTGATGTAATACTATAATGTATCATAAATGGAAAAGTCCATATGTTACTGAACACCAGGTACGATGTAGGTGGTGAAA
CCATTATTGACTGAAAAAGATAAACAGCATTTAATCACGAGTGCATGTTATAATGTTTATGAGAATGTATCATTCAAAGGGAGCTTTGTGTCACTTGTAACCTTAT
GAAGTTTTAAGTTTTCAATGTTTATGCATATTTCTATTTAGCATCTTTGTCAGTATCACCTGTGAAACATGACTCTCATTTAAAGAAACTTTAATTTAATGCATTT
CAAAGCTTGTGTGAATGTAATGGTCTCTGAGCATATATTTATATTTTTTCATACATCAATTACTTCTTTTTGAAACAAATGTGTATTTTTCATTCATCTTAAAAGA
ACCAATAGTATTAATATTAAGCATACATGTGTTTCAAATGTGAGTTGTGAAATCACCATCAGGAGAAATCCCACCATGGACTGGAAAATACTGGTATGTTGTTTAT
GGTGGGTGGTAAATGTAAATTGGGGGGGAAAAATTGTTTTCCCCAGTTTTCAAAGTGTGTCACATAACACTTGTAATGGTCTGATGTTGACCCTCATAAATGTGGA
AAAGAGCCTCCTGAGCATCCTGTGGAAGTGTCTGAAGCTCATTACCGTTTTGTGTGGTCATCTCAGAAGTGGAAGTTTAGGGTTATTGCACGGGTATAAAGCACAC
AGATGTCAAGTGACTGTGGAGAGCTTTCCTTCACTTTGAAGCTCATCTGCTCTTCATGACAAATTGAACATGTTCCTGAATTTGTCATTTGTCTTTCTGTTATTTG
AACAGGCACTGACAACAATGCACCTTTTATTTCCCCCTTTTTTAAGTGGCGAATGTTCTCGAAATAAAACGTT

>emac_TRINITY_DN63430_c0_g6_i5_per3 mRNA_partial cds
AGTCCGCTGAATGAGGATGTGTTTGCTGCTCCTACTAAAGAGGATGTCCCCGTCAACCATGAGGAGATTAAAGATCTACAAGCAAAGATCTATAAACTATTCCTGC
AGCCGGTCCACAACAATGGTTCCAGTGGTTACGGCAGTCTGGGGAGTAACGGCTCTCACGAGCACTACATCAGCATAGCTTCTTCAAGTGACAGCAATGGGAACCT
GTGGGAGGACTCGCGCCGGGAACCGGTGACTTTGCACCAGATCTGTGCTGATGTAAAACGAGTCAAGAGTTGGCGCCAGCACGCTTATCTGGGTTGCAATCACAAA
AATGCTCTTCTTGGGAAACGAGCCACAGCACGTCTGCCCCCTGCTGTTTCCAGCCCTGAGGTTGGGGATCACGAGCAAAGCAGGAAGCAAACATACATTCCCTCCT
ATCAGCAGATAAACTGTGTGGACAATATCATCAGATATTTGGAGAGCTGTACAGGCCCAGCCCTCAAGAGGAAGAGTGACTCTCATTCCTTCTCTTCCTCATCCTC

TACCTCAGAAGACAACAAGCCTGCTGAAGCCACCGACACCGCTCGGGCCCGCTCGGATGTGGTGTTGGACAGTGCGGCCTCAGGCGCTCCGACATCAGCTGCAGTT
GTTGGAACACCTCTGACAGACATCACAATGTCTACTAAGGCCATGAGTGTAGTCTCTGTCACCAGCCAGTGTTCGTACAGCAGCACCATCGTCCATGTGCCACAGC
CTGAATCAGAGGCTACAGCACTGGAGGACACCCCAATGGGCAGTGAGCCAGCTGATGCTGCTCCGACCTCTGTCCGGCCCGCCTCGAGCCCCGCCACAGAGGAACC
AATGTTAATAGGTCTCACCAAGGAGGTGCTGTCAGCTCACACCCAGAAGCAGGAGCAAGAGTATGTGGATCGATTTCATCATCGTATCCTGCAGAGCCCCTACAGC
TCCTATCTTCAGCTGGACAACGGCTCCATGGCTCACTCCCACCAGCCAGGAGACTACCCACCTCCATTGAGCGCTGGAGGGATGAACCGCCCTCGGAGGGGGAAGC
CCAGACACAAGCACCACAAACCCCAGGGATCCTCAGACAGCTACGCTTCCCTAGCTGGCCCTCCTCGTCGGGTCCCAAACTCCTCCTGGCCCTATTCAGAGTCCTC
CCAGCCCAAGATAGGGGCACTCTACAGCCAAACATCCCCTCTCCAGGCGCAATATTTCCCCATGATTACCCAGCCTGGTCCGGGGCAAATTACCAGACAACAGCAG
CTGTCTGGAGGAACCCCCATGGTGCTGCAGCCTCCGGACCAAACCCAGCTCTGCTTCAACATCATGCAGGCTGCTCAGAGCCTGCCGGGCAGTCAACCATTCCAGA
TGGATCATTTCCATAACCTGCACAATATTCCCAACTTTCCCAACATGCAGCCCATGGCTCCAGCACTGGGCGTCAACCCGTACATGACTCCGGTCATGGCTGTCAT
CTTGCCCAGTTACCCACCATTCACTCCAGGGTACCCCCTGTCCACTTCTACTTTGCTGCCTCATGCACCCATCACCATGGCAGGCTTTGCTCCTGGTAACATCCCG
CTCCCTCAGCCCCCGTTCCAGGCCCAGCCAGGTCCCCTCACTCAGACCAGCCTCAGCCCTCTGCTTTTGTCACCCAGAGCCAGCTCCTCTGTCGGGGAGGAGGAGG
AGGTGGCTGGGCCTCAAGCTTTATTCTCCAGTTCTCGCTCCAGTTCTCCACTGCAGCTGAACTTGCTGCAGGAGGAGCTGCCAAAGCCGAATGAAGGGCAGAGCAG
TACCGGGCTCAACCATGCAGAAAGCCTCTGCCAACAGGATGCCAACCAGGGTGATAACCCCAGTGGGTCAGGGAACCATGATGCCCAGTCAACATCCAGTGAGCTG
CTCGACCTGCTGATGCAGGAGGATGCCAGGTCTGGGACCGGCTCCAACGCCTCTGGGTCTGGAGAGTCCAGAGGCTCCCTGGGATCTGGATCTGGCTCCAATGGAA
CCTCCACCTCACAAACTGGCAGCAGCAACAGTAGCAAATACTTTGCCAGCAACGATTCATCGGACACATCCCGAAAAGCCCGTAAGAGCCAGGAGGCACCGGCAGA
GCAGCAGTGCAGCTTCGACAGTCGGGCGGAGAACTCTCTGTGGGGAATGATCCAGCACACACCTGAGCGGGTCATGATGACGTACCAGATCCACAGCAGGGATCAA
AATGAGGTGTTGGCGGAGGACAGGGAGAAGCTGAGGGTGCTTCAGCCCCTGCAGCCATGGTTCAGCTCGGAGCAGAGACTGGAGCTGGCAGAAGTCCATCCCTGTA
TCCAACAGCACATTATCCCACAGGAGATAGACACACAGGGTTGTGTGAGTTGCAACGCGGGACAAGGGGTCAAAGTGTCCCCTCCTCCTACTGCCAATGAGAGCCC
ATCCTCTCTGGAGATCCCACAGCAGGACTCAATCGTACCTGTACTGGACTCTTGAGGGAACCCCTAAAAAGTACCAACCAGCTCTGCCAGAAACCTTAAAAAACTC
CCACCATGCGCCTCCCGAACCAGAGGAGTCGACCCCTATGCTTTAGTTTGATGTTTACCATGTATTGTTAAGGCTTTAGAGACAGGAACAATTTCTAAGGACCGGA
GGAGATAAAGGAGCTTTCCTCTCAGCAAGCCTAATGTGACTGCCTTTAACGGACCATAAAAATGATACCTCTCTCTTTGTAGTGTTCATAGGCCCTTGTAGGGGTT
AAGTTTGTATTTAAAACTGAACAGAAATCGGCACAATGAACCACCTGACACTCAATGGAGGAAGATGTTTGTTGACCCTAAGCATGAGCACTGTGTACACACAGAC
TCAGGTTGGATGTGTGACAAAGACAAAGACAGTGCAGGCCCTGTTTGAGGGTCAGAGCACACAACAGACTCTTCTTCAGAGACATAGAGTGCTGTACGGATACCAC
TTCAACTGCACAGAAACGTACCATGGTTGACCCATGGTAGACATTTTTGATGGACACTTAGACAAACACACATGTTCTGCTCTGTTCATAATGTTCCTGTCCTAGT
CCTGTTAAGTGATTGTATTTGAGATGTTTGTCTATTCCATAGGGTGGATTTTCTCATGTCTAGAACATCTGTTTTTATGTTCGTCCAACTCAGAAACAACATTAGG
AGTTCAAATTGAATTTATTGCTACAACATAATCTCAACCCGACGTCGTGTACCAGCACCCCACAGTTACTTCAACTTACCTTTAATGTAAAGACACAGCTTTGATC
CAAATGTCCTCAGATCCGGGGTTTACTCACTGGATTTATTTCCTCTTTAGACTCTGTTTCTGTTGACTGGCTGTTTGTTTGTGTGTGAAAGAGAAAGACCGAGGGG
ATTTAAATGACCCATTCTGTAAATGTTGTTGCTAATTTTTTCATACTGGCTTCAGATTGCAAGTGTGTGAGGTTTGATTGAGTGTGTGAGTGTGAGCGTGCAATCG
ACGATGCAGTTATCCAAAAGCGCAAGACAGAACAGGGTTTTCTACCATTTTTAGATTGAAAGAATATTACAGTACAAAGTATTTTATAGAAAAAGGACACAAAACC
AACTCCACAAAGAGTTTAAAAACGCCATTTGTCTTGAATTCAGGGACTTTTGTTTGTGACACAAAATAATGAATAAATGATGTCGTAACAAAAAA

>emac_roraa_fin_HQ_transcript/45962_roraa mRNA_complete cds
GGTGGACAATCAGACCTACACACCCACCTCCAGACACACTCGCATTTCTCATTTCTTACT
TCTAGTCTAAATTAGATAGCAAGACGTAGTGCCGGAGCTCACAGTTCGCATAGAATTAGT
TTGAAGTACTAGCGTTAGTTAAACAGCAAATCTGAAATTCAACATGGAGTCCCCTCCAGA
TCCAGCGAGCGACCCGGGCAACAGCGCCTCGGAGCCGGCTACCCCGGTCAGGGAAACCCC
GGTAAACCTGGAGACGCTCCGAAAAGCGGACCATCCAGCCCCGGTTCGGAGGCAGACCTG
CTCCAGCACCAACAGAGGTATCTCAGTAACAAAGAAGACACATACCTCTCAAATCGAAAT
AATTCCCTGCAAGATCTGTGGAGACAAATCATCAGGCATCCATTACGGTGTGATAACATG
CGAAGGCTGTAAGGGCTTCTTCAGGAGGAGTCAGCAGAGCAATGCAGCCTACTCCTGCCC
CCGTCAGAAGAACTGCCTGATCGACCGCACCAGCCGCAACCGCTGCCAGCACTGCCGGCT
GCAGAAGTGCCTGGCAGTGGGCATGTCACGAGATGCGGTGAAGTTTGGCCGAATGTCGAA
GAAGCAGCGAGACAGCCTGTACGCTGAGGTGCAGAAGCACCGGCTGCAGCAGCAGCAGCG
TGAACACCAACAACAGCCAGGAGAGGCGGAGCCACTCACACCTAGCTATGGCCTCTCAGC
CAATGGCCTCACAGAGCTCCATGACGACCTCAGCGGCTACATGGATGGTCACACTCCTGA
TGGCAGCAAACCGGACTCTGCAGTCAGCAGCTTCTACCTGGACATCCAGCCATCTCCTGA
CCAGTCAGGCCTGGACATCAACGGCATCAAGCCGGAGCCCATCTGCGACTTTGCCCCCGG
CTCTGGCTTTTTCCCTTACTGCTCCTTCACCAATGGAGAAACCTCCCCTACAGTGTCCAT
GGCTGAACTAGAGCACCTGGCCCAGAACATCTCCAAGTCCCACATGGAGACATGTCAGTA
CCTGAGGGAGGAGCTGCAGCAGATGACCTGGCAGGCCTTCCTGCAAGAGGAGGTGGAGAG
CTACCAGAGCAAGCCCCGGGAAGTCATGTGGCAGCTGTGTGCTATCAAAATAACGGAGGC
CATTCAGTATGTGGTGGAGTTCGCCAAGCGCATCGACGGCTTCATGGAGCTGTGTCAGAA
CGATCAGATAGTGCTGCTGAAAGCAGGCTCTTTGGAAGTTGTGTTTGTCAGAATGTGCCG
TGCCTTTGACTCGCAAAACAACACCGTCTATTTTGATGGAAAGTATGCCGGACCTGATGT
GTTCAAGTCATTAGGCTGTGACGACTTGATCAGCTCCGTCTTCGAGTTTGGGAAAAACTT
GTGTTCTATGCACCTGTCTGAGGATGAGATCGCCCTGTTCTCCGCCTTCGTGTTGATGTC
TGCTGACCGGTCTTGGCTCCAGGAGAAAGTGAAGGTGGAGAAACTCCAGCAGAAAATCCA
ACTGGCCCTCCAGCACGTCCTGCAGAAGAACCACAGAGAGGATGGTATTCTTACAAAGTT
GATATGCAAAGTGTCGACACTGCGAGCGCTGTGCAGTAGGCATACAGAGAAGCTTACCGC
TTTCAAAGCAATATACCCAGACATTGTGCGTGCCCACTTCCCTCCCTTATACAAGGAGCT
GTTCGGATCAGACTTTGAGCAGTCCATGCCCGTTGACGGGTAGCAGCCCTGCCAGGTGCC
AGTGTGCCCACCGTAGGGGAATGTGGAGGAGGAATCTGAGACACTTTATTGGTGCCCTGC
ACACACCAGAGCGGACAGATGGCACGCTGTTCAACTTCTTCTTTTACATCGGAGGGGAGG
GCTTAGGGAGTTGATTCTCAAGGTTTACTTTATTGAATTTAGTTCTCTTTGGAAGACTTG
TGGGACCCAACAACCCATGGGACCTGAAGAGGAAATAGGGACAAATAATTTCTGTCTGGT
TGAACTTGACCCTCTTTTGCGCATTTCTGACATAATTCACTTTATTCAATCATTTTCACC
AGTATTTATTGCATTGCCTTTATTATGATTTACTATATGACTCAGTAATAGTACAAACTA
TTACTGAGTCACTTGGAAGTAAAAAAGCTGAAGAAGTCTTCAATTAGAAGATAAATCTGC
AAAAATCAGTATTATTTTTCAAATTGCATTCCCAAAACGTTTAAGTGTTTTCTCCTTCAA

ACTTTGAATAGTCTTATCCCTTACCTCCCCACAACTGAAACAAAGTTAAAATGCGCACAT
TTTGGTCAAGTTCAGCCCCTTTTCTCTTGTGTTCAGTCGAATATATTTTGGACGCAGTAC
TGGGAATGAAAACTCACAGCGACTTTGAGGGAGGGTCTGTCGGCATGGTGATGCAATCGG
CACCTCTCTATGGACAATCGTT

>emac_TRINITY_DN57227_c3_g1_i1_rorb mRNA_partial cds
CCTGACGCCGGTGCCCAACCTGTTCAGCTATGGAGGCTACCAGGACAGCCAGCTGGGACC
CAACAATGTCAGCATGGGAGAGCTGGACCGTATTGCTCAGAATATCATCAAGTCTCACCT
GGAGACGTGTCAGTACACAACGGAGGAGCTGCAGCAGCTAGCCTGGCAGACGCACTCCTA
CGAAGAAGTCAAAATGTACCAGAGCAAGCCCCGGGACGTGCTGTGGCAGCAGTGTGCCAT
CCAGATCACCCATGCCATACAGTACGTGGTGGAGTTTGCAAAGCGCATCTCAGGGTTCAT
GGAGCTGTGCCAGAACGACCAGATCCTCCTGCTCAAGTCTGGTTGTTTGGAGGTAGTTTT
GGTGCGGATGTGCAGGGCATTCAACCCTCTCAACAACACTGTGCTCTTTGAAGGGAAGTA
CGGCGGCATGCAGATGTTCAAAGCTCTAGGTTGCGATGACTTAGTGAGTGCCGTGTTTGA
CTTTGCCAAGAGTTTGTGTTCACTGCAGCTGACAGAGGAGGAGATCGCTCTGTTCTCGGC
AGCTGTACTCATTTCCACAGATCGGCCTTGGTTAATGGAGCCTCGGAAAGTCCAGAAGCT
CCAGGAGAAGATCTACTTTGCTCTGCAGCACATTATGCAGAAGAACCACATGGACGAAGA
TGCACTGGCAAAGCTGATCAGCCGAATTCCAACGTTGTCAGCCCTGTGCACGCTCCACAC
CGAGGAGCTCCAGGCCTTCCAGCAGCTCCACCCAGAAACAGTCAACGTCCTCTTCCCTCC
GCTCTACAAAGAACTGTTCAACCCTGACCCCAACTCTGGGGTCATGGCCATACCCAAGTG
ACTGCCTGTGGTCCCAACGCAACAAACGGCATCAGCGGAGGACCAAGAAAATGAAAAGAC
AGCCACGTCTGACGAGACGACAGCAGCCATGTCGACTATGTCCTCATCGTGGCAATCACA
TATTCCTGAGCTCCCCGAGCTTCATTATGAGCAAGGGGGTGAGGAGCCATCTGCCAGGAA
CTCACAGTTACCGCCAACAATACTAGGAATGTCCAGCACTTATTCCATTCTGTTCACCGA
TACAGTCTGAAGAATGTATATATAAATGAAACGCGCCCGAGCCACAATTTAAGTGGGGCT
TGCTCCCGTCTCCTGAACTGAACTGAACTGACTGACCAGAAATGTACAGACTTTAAAACA
TCTAGGAACAATGTTAAGCTGTCAATCTTTTTAAAAAAATAGGTAAGTTAATTTGTTTTA
CCTTTGAACAAAATCCTATTGGTTTGTTCTTTTATTTTCGAGTGAAGAAATTGTGACGCT
GGACGGGGGAGGGAACTGTGGATTTAGAGAAGCTATTGTAGATATTCTTCTAGTGCAGAG
CGGATTCCAGTATTACTTCTTGTTCTGTTAAAATAAGTTAGTCCTAATTTAACTATACAG
CAGCCCACTGTGGCTGACTATACCTTGTCTATGTGTTTTTATTTTATTTGATAGTTTCTT
GTGTACAGAATTTGTAAAGTTGAGACATGTAAAAGAATATTGAGTAAAACCGGGAGACGA
TTCGGGTTTTGTTGATATATAGAAGCCTTGTCTGGATTTTGTTCATATGTAAAGAAGGTA
CTGCATCCTTCTGATGAGAACTTATAGAAATGTAAAGAGAGCATATGAAGGATTTAAAAA
GAAAATAATGCCTATTACTTTTCAAGGTGAAGATATATACATAAATATTCTATTTTGCAA
GAAAAAATAATCTGGCAGTTCTGCCATCCTATCAATCATTTTCTTCTCGCTGCAATAATA
AAGTCCAATAAAAGATGGACTGCTTTCGAACAATCAAACAGAAAAAAACAATCAGAAACT
AGGACCTCACAATCAACCCCACCCCTTCCCAAATTCCCTTCACCCTGCCCCTTATTTCTT
ATGTATTGCATCCTTGACATGAGATTTCAGCCGACAGAATAGTCTAGAAATCACAATGAA
GAAGTGGATAAAAGTTCCACCAAAAACACTAAAGACAAGCTTGAATTCAACTTTCCAATG
TGAAAAAAAACAACCTGTATGAGTGCTGTATGCCCTCCCATCACTATTTGCACAAATCAG
TCCGTGGATGTCAATCACTCCGGGCTGAGAATATGAAGGACCCTTTTTTCTCTAAAGAGC
AATAATAACTTTCCACTGCCTTTCCGATCTTTCCTTTTCCTTCTCTTCCTATGTCCAGCT
GTTCAAAACAGGATAGCAGTTTTAAATAGATTGCACTATATTTGATCATTTATTATGTCT
AACAATCATCTCTATTGATGTTACTCATTTCACCGCAGCATCTGCAGTGTTTTTAACGTG
ACGTAACAGGATCTGGGGTTAGTACACAACGCAGAGCACTGAAAAAAGAAACTTGTGAAG
TTATGAAGATTAGCCTCCGATGATCTTTAACACTTTAAATGCTGCAATATTACATTGACC
AGTTACACAGACGGATGTGAAAGACAATCACTTGTTCTGCTTTTCATTTAGTCATTTCAT
AATTATTAGCACTTACAGGTTTTTCTATACTAATGTTAGGTTGAATGTGATTTGGCCACT
GTGTAAAATCTGGGTATAAATCACTGTCTTTTTGCATTTCTCTATACGCTTACTTTTGTA
TACAGATCTTTGATGCGCTACGGAAAATCCCCTATTACAATCTCCACTTAAAAATAGCTA
TTCACCAAAATAAGTGACCCGTTAAAAAATAAGTGCACAGCATTAATCAGCTGATTGTGC
TTTTACAGACTGTTATCAATTTACTTGTTTGACATAATGATAATTTTGCAAAGAGGATTT
TCTAATTCTAAGACAGCACTTAGATTTTTAACCTAACTTCAATAATAATGATTAGTGGAC
AATGATCTATTGTTGTCCGTGCACTAAAAGTGGAGATATGATTTGTCTGTAGTTGCAAAT
AAACAGTGGCCAAATCCTCAATGTGCTCTAAGCATGGACACAATCAGACTAATTATCCTT
TCATTAACATTACCGTGTGCATCCTTTAATCTCCACATAGTCAACATAACGATGTGGACA
TCCTTCCTCCGACCCCTGACCCGACCCCAGATCAGAGGCCAGTGACTCTTAAGACAATCA
CTTGCTCATTAAGGCTCGATGCACTACAGCATCTCTCAGGGAGCAGCTCTTTTTCTGACT
TTGCATTAAAAGAGGGTTTAATATTTTTGCAAATTTACAAAAAAAAATGTAATTATAAAA
TA

>emac_fin_isoseq_HQ_transcript/31310_rorc mRNA_complete cds
AACTCTCTCCAGCCTGGGACGGTAAGGGCGCACGGAGGGCGCGGGGAGGGGGAGGCCGGG
GCGCGCGACTTGTGAACGGCCGGCCGGTCCAGCTGCACTCTCTCCCCTCTGCTGCTGCTG
CCTCGGGGGGAATTAAATTATTTCAAAACGACCTCACAAAAGGAGGTGTACTTCGGAACA
AACGGGAGGCTGACAGAGAGAGGTAACAACATGACGACCGAGCTGTGGCTCCCGCTGTAA
GGACAAGAGGGAGGCGAGATGGCAGTGCATGTTTTCAAATGCCTGCGCCGGCCCGGGCGC
TAGAGCAACCAACAGAGAGGTGAGTGTGTAGGACCACCCAGTGTTTTGACAGGAGAGTTA

TGGAATATGAGGAGCTCGACGTGCCCCCCACTGACAACCCTATCAAAAGAGAAGGAGCCA
TGTCCAAGAAGACTCATTTGACCCAGATTGAAGTTATTCCATGTAAGATCTGTGGGGATA
AGTCCTCTGGAGTCCATTATGGAGTCATCACTTGTGAGGGCTGCAAGGGATTTTTCCGGC
GTAGCCAGCTGCCTACTGTTTCCTACTCCTGCTCCAGGCAGAACAACTGTCAGATCGACC
GGGCCAGCCGCAACCGCTGCCAACACTGCCGCCTGCAGAAGTGCTTAGCACAGGGCATGA
GCAGAGATGCTGTCAAGTTCGGACGGATGTCCAAACGTCAGCGGGACTCCCTGATTGCTG
AAGTGGAGAGGCACCGACAGCAGCAGCAGCAGCAGCTTCAGGAAGACACCCCGTCTCTCT
TGTCCTTCCCCACCAAGGCCCGTCAAGACCGCTCAGCGCAACTCCTTCAACCCATGGCCT
CCACCTACTCCTTCACCGGGGAGTCCGAGCTGCTGTCCTACACGGCTGATGTCCACCCTT
ACCTGATGTGCTCCCCGAATGAGTCCCAGGTGTCGGGTATGATCTACCGAGGCTCCGCTG
TGTCTCCCACGTCGAGATCCCAGGGGAGGGGCGACAACAGCGGACTCCCTGAAAGCGGAT
TTGACTCCAGACAGCCAACTCATGATCTGGTGGCGATTCACCCCTACAGCCCTCTGGAGG
ATCCTTACAGCCTCTATCCTCACTCTTTGAGAAACATCGATGAGCTGTGTGCCAGCATTG
TGCGCTCCCACAGAGAGACCACTCAGTACAGGGCAGAGGAGCTGCAGGCTCTCAGATGGA
AAGTGCTCAGCAGAGAGGAGATCCAAGCCTACCAGAGCAAATCAGTGGATGACATGTGGC
AGCACTGTGCCATCCGACTGACTGATGCTGTCCAGTATGTGGTGGAGTTTGCGAAACACA
TCCCAGGTTTTCGTATGCTCAGCCAGAACGACCAGATAGCTCTCCTGAAGACCGGCTCTA
TGGAGGTGGTTCTAGTCCGGATGTGTCGCTACTTCAACACAGAGAACAACACCGTCTTTT
TCGATGGGAAATTTGCTGGAGTTGAAGTCTTCAAGTCTCTGGCATGTGGTGATTTAATCA
CAGCAGTGTTTGACTTTGCTCACAATATGTGTGCTCTCAAGCTCACTGAGCAGCAGATCG
CTCTCTTCAGTGCTCTGGTGCTGATCAACACAGAGCGTCCATGTCTGGAGGACAGAACCA
GAGTTCAACGAGTGCAAAGAAGCGTGGAGTTTGGACTCTCACACATTCTCCACCGAGACA
ATCAAGAAAGTCTATTGCACAAGCTGTACCAGAGGATGGCAGTGCTGCGTTCACTGTGCA
GTCTGCACATGGAGAAGCTGCGCTGGTTCAGTCAGCGTTACCCACTCACCGCTCACTCTC
TGTTCCCTCCTCTTTACAAGGAGCTGTTTGCCTGCGAGGCTGAGCAGCTGCCGGGAACCA
CTCACTGATGATTCACTAGATCATGCAAAGACACAAACTTATTTCAAGTCTTCTTTACTT
CCTTGACAGAATCTATTTTTCTAATAATTAAAATAAATGAGAAAAAGCACTGTAATCTTA
ACTTGTTTACTACTGTATTTTATGAATTCCAGCAGAGCTTTTTGATAGCTTGCATCAGAG
ATAAATATCAATATTTTCACAGTTTAAAAGTGTTTTGATTATTACAGGGAAACGTGTGCT
TTTGTTGTTGTTTGACAGTACATTGTTAGAGGGGTCCAGAATTAATTTTCTGAGTTACAG
TGCATTCGGACACAGACATTTAAAAAAGCAGAACACCTTTCATTGTTTAAAATGCTAGTT
CAGAAGAAGGAAAGTATGACTGTTTATGTATGTGTTAAATAAGCCCGAACAGAGACACAA
GGCACATAAGGAAGATGATGAAGGTTATGTTTTTTTCCAATTTTATCTGATGTTCAGAAT
TGACAATCAGTTGCTAATTGGCTTGTGGTGCTCTCTCTCTCTCAGTATTTCAGGAGCCTG
TAAAAAAATCCTACCTTCACATACCACATACACATTTGAAGGTTAATGTAGTGCGTAACA
AGTAGCCCTGCAAATAAATAAATATTGGTGTGTGACAATTACATACAGAGTGTTAAACAC
ACCCAGAAAACCGAGACAACCTGACCTTGTTTTTGTAGAGAAATGCATCTCCTCTTCAGG
TAACAGATCGACCATAAGTCCTACAAAGCAGCCGTACCTCTTAAATGTTATCCTTTGTGT
TTGAACAGATCCTAACAGGCTTTTGGTGTTTTTAAGCCTAGAAAAGATGTTGCATTAACT
GTATAAATTCAATTAAAATGTGAGCAATAAAGTAGT

>emac_emac_TRINITY_DN61725_c0_g14_i1_rorca mRNA mRNA_complete cds
CTGCCACACTTTCCCACGCACGCTGGAACCTCCTTCGAAGGCGCGCGGAGGGTGGCGCGC
CCGTCAGATCACGGGGCACGAGGCGGGCGGACCCGCGGGTGAGGATCATGTGTAGTCCAC
GCGCCGAAACTGAGTTAGCCTTCAGCGAGCTGCAGAGCAAGAAGGAGCTGAGAGAAGTGG
AGCTGCTGCTTTATCACCGTTTTATCACAGTAATATCGCTTAGACCGCACGGAAGTGACT
GACAGAGATTAACTAGTAAACACACACCGTTGCTTTTATTTGTTTTAAGGTCGGTGCATG
CAAAGAAAGGCTGGAAAATCCATATGAGGATTTGCTGTTGTGACAGTTTTCAGCTGCTCC
GTTTTGGAGCAAATAAGTGCGCGTCGCTGTGTTCTGGGCTTCATTTCCACCTGGAGTTGT
ATGACTTGAAGAGTTTGCGTAGACTTCACCGCCACCATGAGAGCTCAAATAGAGGTAATA
CCGTGTAAAATATGTGGGGACAAATCATCTGGGATTCACTATGGTGTCATTACCTGTGAA
GGCTGCAAGGGTTTTTTCCGCCGCAGCCAGCAGAACAATGCTATGTACTCCTGCTCGCGA
CAGAGGAACTGTTTAATTGACAGGACCAACCGTAACCGTTGTCAGCACTGCAGGCTGCAG
AAGTGTCTCGCTCTTGGCATGAGCCGTGATGCGGTCAAGTTTGGCCGAATGTCCAAAAAG
CAGCGTGACAGCCTGTATGCAGAGGTCCAGAAGCACCAGCAGTCCCAGGAGTGTGCGGCG
CTCGGAGTCCGCGAGGAGAATACCGACACGGCCGACCACAGCCGCACCTACAGAAGAGTC
TCCAGCACCACGCTCAGCGATCTGGACGACATTACTATGCTGCAAGAAGGCCTGCTTTTC
GACCTGCCGCTGACCCCGGAGGACGCGGGGAGAGAGTACTGTAACCTGGACATGCTGGGC
GGCAGTGCAGGCAGCAGCTCCTCCGCTCAGAGTTCACCAGAACAGACCAGCTTGGACTTT
GCAGAAGGCAACCACAGCATCAAGCATGAGTACCAGCTGTTGCACGACTCCGGACTCTTC
TCACATGCTATCCTCAACCCGCTGCCTGAGGGCTGCTCCATGCTCGAGATAGAGCGTATC
ACTCAGAGTGTCCTGAAGTCCCATATTGAGACGAGCCATTACAGCACAGAGGAGCTGAAG
AGGATGGCGTGGACCCTGTACAGCCCGGAGGAGACGCGCTCATTCCAGACCAAGTCAGCT
GAGGTGATGTGGCAACAATGTGCCGTTCACATCACTAATGCAATCCAGTACGTGGTGGAG
TTTGCCAAGCGCATCTCTGGCTTCCTGGACCTGTGTCAGAACGATCAGATCATCCTCCTC
AAAGCAGGCTGCATGGATGTTCTTCTAATCCGCATGTGTCGGGCCTATAACCCCATCAAC
AACACAATGCTGTTTGATGGAAAGTTTGCCACTGCACATCTCTTCAAAGCTCTCGGCTGT
GACGACCTTGTGAATGGAGTTTTCGACTTAGCTAAAAGCCTGAGTCGTATTCAGATGTCA
GAGGAAGAGATGGCTCTCTTCAGCGCTGCTGTGCTGCTCTCTCCAGACCGCCCCTGGCTG

ACAGATGTTCAGAAGATACAGAAGTTGCAGGAGAAAGTGTACGTGGCTCTGCAGCGCTGC
CTACAGAAAGAGGGAGCGTCAGAACAGAAACTAGCTAAGATGGTGTCTAAGCTTCCCGTT
CTGAAGTCCATTTGCAACCTTCACATCGACAAACTGGAGTTTTTCCGTCTGGTCCACCCC
GAGACAGCGTACACCTTCCCTCCTCTGTACAGGGAGGTATTTGGCAGCGAAATCACCTTC
CCAGACTCCACAGAGGGCTAGAACTATTTAGCTAAAAAAGCGGGATTTTAATTGAGACAC
TGAAGTGTAGGGAGGGGGAGATAAAAATGTGGCAACCAGCTGAGGCAAAAACCCCACTGC
TCTCACCTAAGAGCTCCCAAACTGTGTGGAGTAGCTCGCACTGCCCTGCGTCCGGTCCAT
TCAGCTTCAACCTAACACTACAGGAACTGCGGCTGGGCACTCAGTGAACGTTACCTCTTT
TTTTTAAGGAACTTATTGAAGAGTAGCGTATTTTATTTGTATACTCAAAAATATTTTTCT
ATGAGAAGACCTTAAACCTCTCATCAAGCTGAATTTGACTTTCACCATTCTCACAGTTCC
TGTAAGCTATCGTTGAATGTATCCCCTCACCACCCTAAGCTGGATATAGAATGTAAATAC
TTTAAATCCTCTCCCTCTCTGTGGAGGTCTGCAGTAATCATGCCTGTTTCAGTTCTCAGT
CTAGTAACAGTATATATTTTTCGGTAACGCCCATATTTTGCAGTTTTGTAACGTATGAAC
AAACTGTTTAAAGGGACCATGATCAAACTAACAAACAGTGTTTTTTTTTAAGACTGTGTG
AACCAAGGGCACCTATAAACTGTTGCACACAGCACAAAGCACTTATCTTTGTAGGTATTT
TTACTAGAAACCCACAGACACTGATGTAAGTATGCTTTGTTTTTTTGCAATCGGTGACCT
CAGCCTCAATACCTTTTCCCAGTGAAAATGCCACTCAGAAAGTGAGGGAATCACAGAGGT
ATTTGAAAGTAAGCTCCTTTTTATACACAAGTAATATATAAAACCTCATTTCATTTTAAG
AGCAGTCCAAGTGAGAAGCAGAAGAATATTCTGGTGCTCAGGTCTCTATACTGTCTGCCC
AAACTCTGGGGTTATGGGATGGAAAGATTGTATTCTAATTGGTAATTAGCATTTACAGTA
AGAGACCTATTCAGAAACCAAAACAAACTCTTGGCCCAAGAGGCTGCCATCTGTGAAAC

>emac_fin_isoseq_HQ_transcript/45003_rorcb mRNA_complete cds
GGGGGTTAGCAGGGTAGACTGAGAGGAAAGCAGGCGGAGAATCAGAGCGCAGAGGGGGGG
TCCGATCAATGGTCGTCCACTGTGACCGTGCGACTATTCTGTCAGAAAAGTTTTGCTTCA
CGTTTAAAACGATGCACTGAGAGATTGCGATGCAATCCGTTGCGACTTTTTTCACTCTTT
TATTTCGCTCGGACTTGACCGAGCGATAGGGGAGAGGAGTTTCGGGTAGTTATTTTTCCA
TAAATTTTTGGTCGTGCTTTGCGAAGGCTGTCTGCAGCCAGGACGAAGTTAATATTTACG
AAGCCAGTTCACTCTGGCTCCAGCGGCTTAATCTGCTCGACCGTGTCAACTACATTAACC
ACATGTAGCCGAGTTTTACCATCGTTCTGCCAAGAGAAAGAAAACATCAACATGAGAGCT
CAGATCGAGGTCATCCCCTGTAAGATTTGTGGGGACAAGTCCTCAGGAATCCACTATGGC
GTCATCACCTGTGAAGGCTGCAAGGGTTTTTTCCGTCGCAGCCAGCAGAACAATGCCATG
TACTCTTGTTCCCGCCAGAGGAACTGCCTGATCGATCGAACAAACCGTAACCGCTGCCAG
CACTGTCGCCTACAGAAGTGTCTGGCCTTAGGCATGAGCAGAGATGCTGTCAAGTTTGGT
CGTATGTCCAAGAAGCAGCGCGACAGTCTCTACGCTGAAGTGCAGAAACACCAGAAGTCC
CAGGAGTGTGCAGGCTCTGCAGGCGGTGGCACTACAGCTCTGTCCCAACACAAAGAGGAA
GGTGTTAAAGAGACTCTGAGCCGGTCCTACAGCAGCGGGGGCTCCAGCTCCACCCTCAGC
GACCTGGATGACATCGAAACGCTGCCGGACCTGTTTGACCTGCCACTGACCCCCGAGGAG
GCCAACGACTACTGTAGCATGGAACTGCTCGGCGGCGGAGCCAGCGCGGGGAACACCTCC
GCCTCATCCTCTTCTTCCTCTTCCTCCTCCCCATCCTTGTCCAATCAGAATTCCCCGCAG
CAGACTATGCTGGACGGGGCAGACAGCAACGGAGTCCAGCTTCTGCATTCTCACACACAT
GCACTGCTGGACCATCTGCCTAATGACTGCTCAACAACAGAACTTGAGCGCATCACTCAG
AGTATTGTTAAGTCTCATTTAGAGACGTGTCAGCACAGCGCTGAAGACATGAAGAGATTC
ACCTGGGTGCAATACGCGCCTGAGGAAACACGTGCTTTCCAAAACAAGTCAGCAGAGTGG
ATGTGGCAGCAGTGTGCCCACCACATCACCAATGCCATTCAGTATGTAGTGGAGTTTGCC
AAACGCATTGCTGGCTTCATGGACTTGTGCCAGAATGATCAGATTATATTACTGAAAGCA
GGCTGTCTTGAAGTCCTGCTGATACGTATGTGCCGAGCTTTCAACGTAAACAACAGCTCC
ATTTTCTTCAATGGAAAGTTTGCCCCGGCTCAGTTCTTCAAAGCACTTGGTTGTGAGGAC
CTGGTCAGTGCAGTGTTTGACCTGGGTAAAGGACTCTGTCGTCTACAGCTGTCTGACGAA
GAGATGGCTTTGTTCAGTGCTGCCGTTCTGCTTTCTCCTGATCGACCCTGGCTAACAGAA
GGCCCAAAGGTTCAGAAACTCCAGGAGAAGGTCTACCTGGCTCTGCAGCACAGTCTAAAC
AAGAGTGGGGCTTTTGAGGAGAAACTGGACAAGATGGTTTCGAAGCTGCCCATAATGAAG
TCAATCTGTAACCTCCATGCGGATAAACTGGAATTTTTCCGCTTGGTCCACCCGGAGACT
GCTTTCAGTTTTCCACCACTGTACCGGGAAGTGTTTGGCAGCGATATGACTCTGCCTGAC
TCCACCAACAGCTAGACAGACACAAATAGGAGAGATGGAGAGAGTGTATTAAGAAGTAGG
GAAACAAAAAAAAGGAGTGACAGAGAAGGATAAAGAAAGCTCAGCAACCAGCAGTTTAGA
GACAATCCAGGACTAAATACTCCGTCTCCCATGATCCCAAAGTAGTCCTCCACTTTCCCA
GCAGGCAAAGCACCTTACACTACAGACCACACATGCTTATGGTCTCTTCATATTGTAGCA
TTAATAACAATCGGATGGGCATCGTAATAATAAAAGCCCATTCAGCATGACACAGCTCCA
CGTTCTCAGACACTTATTTAACATCAGATCCACTGTGGATCACCCCACCAGTTATGAATG
TACCTCATCAATAAGCACCAGTGCACAGAGTAAGAACCCAACAACAGTGAATGTACAACC
CTGCCACTTTAATGCAGAGTTTTACACATGGTATTAGTCGTGTTGATCCATAGGTCTCAC
TATCATTACATCTCACAGTCAATATGTTGTGCCTGCGTATTCTTACAGACAATATCCTTG
TACTGACCATCACTTAGGGACCAAGATCATGAGTCTTTTTGTCTGATAT

>emac_fin_isoseq_HQ_transcript/74008_rorcb mRNA_complete cds
GAGATCATGGGAGAGTCCACCGCGCTTGGAGAAAGTGGGTTAGCAGGGTAGACTGAGAGG
AAAGCAGGCGGAGAATCAGAGCGCAGAGGGGGGGTCCGATCAATGGTCGTCCACTGTGAC
CGTGCGACTATTCTGTCAGAAAAGTTTTGCTTCACGTTTAAAACGATGCACTGAGAGATT

GCGATGCAATCCGTTGCGACTTTTTTCACTCTTTTATTTCGCTCGGACTTGACCGAGCGA
TAGGGGAGAGGAGTTTCGGGTAGTTATTTTTCCATAAATTTTTGGTCGTGCTTTGCGAAG
GCTGTCTGCAGCCAGGACGAAGTTAATATTTACGAAGCCAGTTCACTCTGGCTCCAGCGG
CTTAATCTGCTCGACCGTGTCAACTACATTAACCACATGTAGCCGAGTTTTACCATCGTT
CTGCCAAGAGAAAGAAAACATCAACATGAGAGCTCAGATCGAGGTCATCCCCTGTAAGAT
TTGTGGGGACAAGTCCTCAGGAATCCACTATGGCGTCATCACCTGTGAAGGCTGCAAGGG
TTTTTTCCGTCGCAGCCAGCAGAACAATGCCATGTACTCTTGTTCCCGCCAGAGGAACTG
CCTGATCGATCGAACAAACCGTAACCGCTGCCAGCACTGTCGCCTACAGAAGTGTCTGGC
CTTAGGCATGAGCAGAGATGCTGTCAAGTTTGGTCGTATGTCCAAGAAGCAGCGCGACAG
TCTCTACGCTGAAGTGCAGAAACACCAGAAGTCCCAGGAGTGTGCAGGCTCTGCAGGCGG
TGGCACTACAGCTCTGTCCCAACACAAAGAGGAAGGTGTCAAAGAGACTCTGAGCCGGTC
CTACAGCAGCGGGGGCTCCAGCTCCACCCTCAGCGACCTGGATGACATCGAAACGCTGCC
GGACCTGTTTGACCTGCCACTGACCCCCGAGGAGGCCAACGACTACTGTAGCATGGAACT
GCTCGGCGGCGGAGCCAGCGCGGGGAACACCTCCGCCTCATCCTCTTCTTCCTCTTCCTC
CTCCCCATCCTTGTCCAATCAGAATTCCCCGCAGCAGACTATGCTGGACGGGGCAGACAG
CAACGGAGTCCAGCTTCTGCATTCTCACACACATGCACTGCTGGACCATCTGCCTAATGA
CTGCTCAACAACAGAACTTGAGCGCATCACTCAGAGTATTGTTAAGTCTCATTTAGAGAC
GTGTCAGCACAGCGCTGAAGACATGAAGAGATTCACCTGGGTGCAATACGCGCCTGAGGA
AACACGTGCTTTCCAAAACAAGTCAGCAGAGTGGATGTGGCAGCAGTGTGCCCACCACAT
CACCAATGCCATTCAGTATGTAGTGGAGTTTGCCAAACGCATTGCTGGCTTCATGGACTT
GTGCCAGAATGATCAGATTATATTACTGAAAGCAGGCTGTCTTGAAGTCCTGCTGATACG
TATGTGCCGAGCTTTCAACGTAAACAACAGCTCCATTTTCTTCAATGGAAAGTTTGCCCC
GGCTCAGTTCTTCAAAGCACTTGGTTGTGAGGACCTGGTCAGTGCAGTGTTTGACCTGGG
TAAAGGACTCTGTCGTCTACAGCTGTCTGACGAAGAGATGGCTTTGTTCAGTGCTGCCGT
TCTGCTTTCTCCTGATCGACCCTGGCTAACAGAAGGCCCAAAGGTTCAGAAACTCCAGGA
GAAGGTCTACCTGGCTCTGCAGCACAGTCTAAACAAGAGTGGGGCTTTTGAGGAGAAACT
GGACAAGATGGTTTCGAAGCTGCCCATAATGAAGTCAATCTGTAACCTCCATGCGGATAA
ACTGGAATTTTTCCGCTTGGTCCACCCGGAGACTGCTTTCAGTTTTCCACCACTGTACCG
GGAAGTGTTTGGCAGCGATATGACTCTGCCTGACTCCACCAACAGCTAGACAGACACAAA
TAGGAGAGATGGAGAGAGTGTATTAAGAAGTAGGG

>emac_fin_Isoseq_HQ_transcript/3075_timeless mRNA_partial cds
GGTCTTGGCGCGAGACTTGTCCGACAGAAGAGGGCTGAAACCACTAAACAACAATAATAA
CTTCTGAAAAAGATTTACATTTATACACCGGCAGATGAACACGGAACCTGTCGAATAACA
GCGTTTAAACGATCTGGCTGTCTGATGTTGTTTGGACTAAGTCAGGATGAATTGCGATCT
TTTGGCAACGTGCAGTGCTCTCGGCTATCTGGAGGGAGACACCTATCACAAGGAAGCTGA
TTGCTTAGAGAGTGTGAAAGACTTGATCAGGTATTTACGCCATGAAGATGACACGCGTGA
CGTCCGCCAGCAACTCGGTGCTGGCCAGATTGTACAGAATGACCTTCTACCTATGATCGT
CCAGCATGGAGAAGACAAGGCCTTATTTGATGCCTGCATCAGGCTCATGGTCAACCTTAC
TCAGCCTGCCATGCTTTGCTTCGGTAAGGTCCCCGATGACCCAGTGTTCAGACATCACTT
TTTGCAAGTGACTGCTCATTTACAAGCCTGTAAAGAGGCATTTGCCAGTGAAGCAGTGTT
TGGCATTTTAAGTGAGACATTATACACGCTTCTACAACTGGACTGGGAGCAGAGACAGGA
GGAGGATAACCTGCTGATAGAAAGGATCCTGCTGCTGGTCAGAAATGTGCTTCATGTTCC
TGCGGACCCCTGTGAGGAGAAGAAAGTAGATGATGATGCCAGCATCCACGATAGGCTGCT
GTGGGCCATTCACATGAGCGGATTCGATGACCTCATCAAGTTCCTGGCGTCGGCCCAGAG
TGAGCAGCAGTGGAGTATGCACGTGTTAGAAATAATCTCCCTCATGTTCAGAGATCAGAC
GCCTGAAGCTTTGGTGAGCGCCGGTCAGGCTCGCTCAGCAGAGGAGAAGCAGAGAGACGG
TCTGGAGCTGGAGGCACTGAGGCAGAAAGAGCACGCTGCGAAACGCTCCCGCAAATTTCA
GAGGGGAACCAGACACTCTCGCTTTGGAGGCTCCTATGTTGTGCAAGGCCTCAAGGCGAT
TGGGGACAATGACGTGATTTATCACAGAAATATTCATAATTTCAAAAACTATACTCATGA
CACCGGCAAAGCAGTGAGACGTGTTCCTAAAAGGAATCGACAGGCTCGGGAATGTAAGGA
CAAACGCCGCTCAGCCCTAAACGTCCGACTCTTCCTGCGAGAGTTCTGTGTGGACTTCTT
GGAGAACTGCTACAATCGCCTCATGTACCTTGTCAAGGAAAGTCTAAATCGAGAGCAAAC
TCAGCAACACGATGAGACGTACTACCTCTGGGCGCTCAGCTTTTTCATGGCTTTCAACCG
CGGCAACGGTTTCCGTGCAGACTTAGTTTCTGAGACCATGTCCATCCGTGCTTTCCATTA
CATTGAGCGAAACATCACTAACTACTACGAGATGCTTCTTACAGACCGCAAAGAAGCCAC
GTCCTGGTCACGCAGGATGCACCTGGCACTTAAGGCATATCAGGAGCTGCTGCTAACTGT
GAACGAGATGGACCGTTCCCAGGATGAAAGCATCCGTCAGAGTTCCAGTGTTATTAAAAG
CAACATATTTTACCTGATGGAGTACAGGGAGATCTTCCTGACCCTGCTGAGGAAATATGA
TGAAACCAAGCAGCCTCACTCCTACCTCATAGACCTAGTGGAGTCAACTCATCTCTTCAT
GCGCATGTTGGAGCGCTTCTGCAAAGGGCGCAAGAACTTGATGGTTCAGAGAAAGAAAGT
GAGGCGGAGAAAGTCTAACAGTAAAAAAAAGCCCCCGGCACCAGATACTAGTCCGGAGGC
TCTGGCAGAGACCTGGAAGATTGTGGAGGAAGAGCTGAAGGCTTCAGGCTTTAAGTTGTC
AGAGACTTTAACTGAAAGCATCGTGCCATTTGATGCCACATCTGAAACTCCACTCGAGGA
CCAGAAGACGGAGACCATGGTGCGGGTGCAGGATGCGATGCTGGGTCGCATGGGACCGGA
GGCGCTGGGGATGCTGCGAGCTGCCAGGGAGGTGTGGCCAGAGGGAGATGTGTTTGGTTC
AGCTGACGTTGAACCTGAGGAGGAACTGGAACTCCTCAAGCAGATCCTGTTTGCGAATCT
GCCAAGGTCAGCTCCTCCTGAGCCTGTGGCAGAGGAGTATGAGGATGATGCTGAACTAGA
AGAAGAAGAGATGGAGTCCATCCAGGTGTCTGAAAAAGAGTTTGACTTTCTAGACTTTAT

```
CAAGAGGTTTTCCCAGCCCAGTATTGTGCGTCCATACCTCCTCTTATTAAAAACATACTC
CAAAAACACACCTCACACAAACCACTGCATCGCCCGAATGCTGCACCGCCTGGCTTTCGA
CCTCAAAATGGACGCCCAGCTTTTTCAGCTGTCAGTCTTCAACATTTTCAACAAGATCCT
GAGTGACCCCGCTGCAGCTGCTTACACGGAACTCGTGACCTTTGCCAAGTTTGTGCTGAA
CCGTTTCTTTTCCCTGGCAGCAAAAAACAATAAGGCATATGTTGAGCTACTGTTCTGGAA
AAATGTGGGCGCTGTACGTGAGATGACCGAAGGATACAGCAAAGACGGAGAGGAAACAAA
ACCATCATGGACTGAAGAGGAGGAAGAGGAGTTACGCATCCTCTTTGAGGAGCACCGTGA
TTCTGAAGTGCCAGATGTAGTGGAGGCGCTGCTGCCATTACTCAGCAAAAGAAACCGCAC
CCGGCGCCAGGTGGTGACGCAGCTGGTGCACATGGGTCTGGTGGCCAATGCTAAAGAACT
GAAGAAACAGAAGAAAGGTACCCAGATTGTTCTGTGGACTGAGGAGCAGGAGGAGGAGCT
GCAGATGCTCCATGAGGAGTACAAAGACTCTGATGATGTGCTGGGTAACATCCTGAAGAA
GCTCACAGCCAAACGGTCCCGTGCCCGTGTGGTGGACAAGCTGCTCAGTATGGGCTTAGT
GTCTGACAGAAGAGAACTCTATAAGAAGAGGAGTCGCGGCGCTCCAGGGAAAAGCTCTGG
AAAAAGCTCTGGAAAAAGAATGACTGAAGAAGATTTCCTTGAAGAACTAACACAAGGGTT
CCCAGATGATCCAGTGGACAGAGATGATGAAGATGAG
```
